# A high-coverage Neandertal genome from Chagyrskaya Cave

**DOI:** 10.1101/2020.03.12.988956

**Authors:** Fabrizio Mafessoni, Steffi Grote, Cesare de Filippo, Viviane Slon, Kseniya A. Kolobova, Bence Viola, Sergey V. Markin, Manjusha Chintalapati, Stephane Peyrégne, Laurits Skov, Pontus Skoglund, Andrey I. Krivoshapkin, Anatoly P. Derevianko, Matthias Meyer, Janet Kelso, Benjamin Peter, Kay Prüfer, Svante Pääbo

## Abstract

We sequenced the genome of a Neandertal from Chagyrskaya Cave in the Altai Mountains, Russia, to 27-fold genomic coverage. We estimate that this individual lived ~80,000 years ago and was more closely related to Neandertals in western Eurasia (1,2) than to Neandertals who lived earlier in Denisova Cave (3), which is located about 100 km away. About 12.9% of the *Chagyrskaya* genome is spanned by homozygous regions that are between 2.5 and 10 centiMorgans (cM) long. This is consistent with that Siberian Neandertals lived in relatively isolated populations of less than 60 individuals. In contrast, a Neandertal from Europe, a Denisovan from the Altai Mountains and ancient modern humans seem to have lived in populations of larger sizes. The availability of three Neandertal genomes of high quality allows a first view of genetic features that were unique to Neandertals and that are likely to have been at high frequency among them. We find that genes highly expressed in the striatum in the basal ganglia of the brain carry more amino acid-changing substitutions than genes expressed elsewhere in the brain, suggesting that the striatum may have evolved unique functions in Neandertals.

## Introduction

Neandertals and Denisovans are the closest evolutionary relatives of present-day humans. Analyses of their genomes showed that they contributed genetically to present-day people outside sub-Saharan Africa (4,5). However, to date, the genomes of only two Neandertals and one Denisovan have been sequenced to high quality. One of these Neandertal genomes (*Vindija 33.19*) comes from an individual found in Vindija Cave in Croatia (1) whereas the other Neandertal genome (*Denisova 5* or the *“Altai Neandertal”*) (3) and the Denisovan genome (*Denisova 3)*(6) both come from specimens discovered in Denisova Cave, in the Altai mountains in Siberia.

A number of archaic genomes of moderate quality (1- to 3-fold genomic coverage) have yielded additional insights into Neandertal history. For example, genome sequences from five late Neandertals from Europe have shown that they carried little genetic variation (2,7) and were more closely related to the *Vindija 33.19* Neandertal than to the *Denisova 5* Neandertal. A genome sequence from a morphologically undiagnostic bone from Denisova Cave, *Denisova 11*, belonged to the direct offspring of a Neandertal mother and a Denisovan father (8), indicating that the two groups met in the Altai region. The Neandertal mother of *Denisova 11* was more closely related to *Vindija 33.19* than to *Denisova 5*, indicating that a replacement of Neandertal populations in the Altai Mountains occurred (8).

Here, we present the high-coverage genome sequence of a Neandertal from Chagyrskaya Cave, located 106 km to the west of Denisova Cave (9–12) (Fig. 1). This genome provides insights into Neandertal population structure and history and allows the identification of genomic features unique to Neandertals.

**Figure 1:**
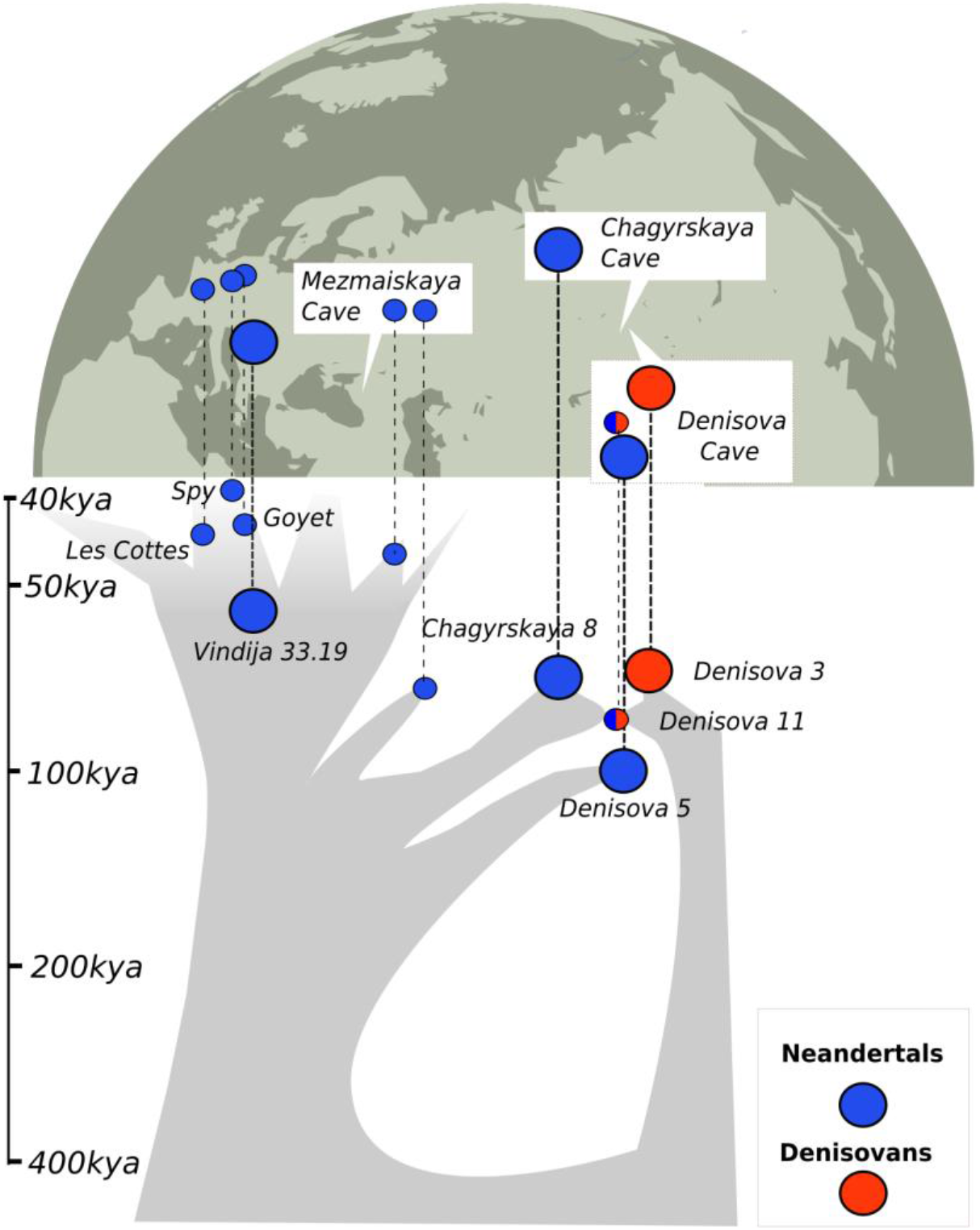
The *Chagyrskaya 8* Neandertal and its relationship with other archaic individuals. Above, locations of Chagyrskaya Cave and other sites where archaic specimens analyzed here were found are indicated. Below, schematic illustration of the relationship among archaic genomes. Split times were estimated using the F(A|B) statistics (1,3,8) (SI7). Neandertals are indicated in blue, Denisovans in red. Genomes determined to high genomic coverage (>27-fold) are indicated by large circles.

## Results

### Genome sequencing and age estimates

We sampled 100 mg of bone powder from *Chagyrskaya 8*, a phalanx found in 2011 at Chagyrskaya Cave in layer 6b (SI1). The DNA extracted (13) allowed the nuclear genome to be sequenced (SI2) to an average coverage of 27.6-fold (SI3). Less than 1% of the DNA fragments sequenced were estimated to originate from contamination by present-day modern human DNA (SI4-5).

We estimated the age of *Chagyrskaya 8* using two different methods (SI6). First, we counted the proportion of “missing” derived substitutions compared to present-day genomes (1,3,6,14). We also used a method similar to Fu et al. (14), that takes advantage of the shared evolutionary history of the three high-coverage Neandertal genomes. Under the assumption that Neandertals had the same mutation rate as present-day humans (1.45×10^−8^ mutations per generation per base pair) (14), both methods suggest that *Chagyrskaya 8* lived approximately 80 thousand years ago (kya), *i.e*. ~30 thousand years (ky) after *Denisova 5* and ~30 ky before *Vindija 33.19*. This estimate is older than the archaeological estimates of ~60 kya (10,12) for the archaeological layer in which *Chagyrskaya 8* was found, which are based on optical luminescence. As re-deposition from lower, older layers seems unlikely (10,12), this might indicate that the genetic dates, based on the current mutation rate in humans, is incorrect. Possible explanations could be that Neandertals had a lower mutation rate than modern humans or that that the modern human mutation rate decreased recently (15). Additional high-quality genomes determined from well-dated Neandertal remains are needed to address these possibilities. Nevertheless, the *Chagyrskaya 8* Neandertal and the *Denisova 3* Denisovan display similar proportions of “missing” mutations relative to present-day humans, suggesting that they lived approximately at the same time (SI6).

### Relationship to other Neandertals and Denisovans

*Chagyrskaya 8* shares more derived alleles with *Vindija 33.19* and other later Neandertals in the Caucasus and in Europe than with the older *Denisova 5* Neandertal from the Altai (Fig. 2). With the Denisovan *Denisova 3, Chagyrskaya 8* shares less derived alleles than does *Denisova 5*.

**Figure 2:**
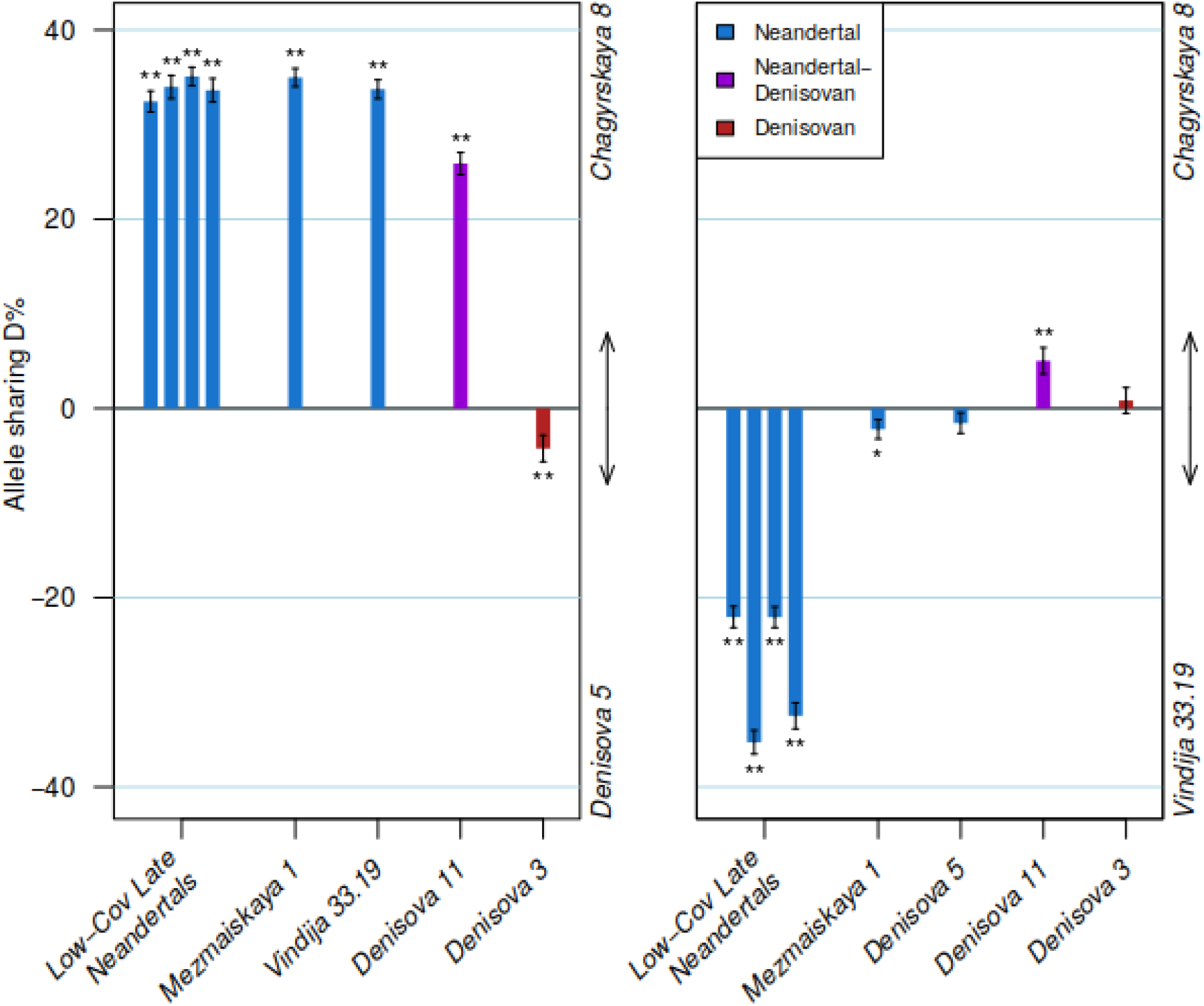
Relative sharing of derived allele among the *Chagyrskaya 8* and other archaic genomes. Positive values in the *D*-statistic indicate more allele sharing with *Chagyrskaya 8* than with *Denisova 5* (a) or than with *Vindija 33.19* (b). Error bars indicate one standard error. One asterisk indicates Z>2; two asterisks indicate Z>3. Late Neandertals of low genomic coverage refer to the genomes in Hajdinjak et al. (2).

When compared to *Vindija 33.19* (Fig. 2), *Chagyrskaya 8* shares less derived alleles with other Neandertals that lived in Europe ~50 kya, *i.e*. approximately at the same time as *Vindija 33.19*. However, *Chagyrskaya 8* shares more derived alleles than *Vindija 33.19* with *Denisova 11*, a first generation Neandertal-Denisovan offspring (8). Since *Vindija 33.19* and *Chagyrskaya 8* do not differ in their sharing of derived alleles with the Denisovan *Denisova 3*, this indicates that *Chagyrskaya 8* is most closely related among currently known Neandertal to the mother of *Denisova 11* found at Denisova Cave (Fig.1, SI7).

### Relationship with modern humans

Non-African present-day humans carry approximately 2% (1,4) Neandertal ancestry as a result of gene flow from Neandertals that occurred between 50 and 90 kya (4,16). Genome-wide, *Chagyrskaya 8* shares more alleles with present-day human populations outside Africa than *Denisova 5* does, and a similar proportion of alleles as *Vindija 33.19* (SI7). However, if the analysis is restricted to previously detected Neandertal haplotypes introgressed in present-day humans (17), or to derived alleles that occur at low frequencies in present-day non-African populations and that are therefore more likely to be introgressed from Neandertals, *Vindija 33.19* shares more alleles with present-day populations than *Chagyrskaya 8*. This indicates that *Vindija 33.19* is more closely related than *Chagyrskaya 8* to Neandertal populations that contributed the majority the DNA to present-day populations.

To test if any modern human population carries an additional genetic contribution from Neandertals more closely related to *Chagyrskaya 8* than to *Vindija 33.19*, we made use of previously published haplotypes inferred to come from Neandertals that are found today exclusively in East Asia, Europe, India or Oceania. Among 300 genomes from the Simons Genome Diversity Panel (18) and 89 Papuan genomes (19,20), the proportions of alleles shared with *Chagyrskaya 8* and with *Vindija 33.19* are similar (SI7), giving no indication that Neandertals closer related to *Chagyrskaya 8* than to *Vindija 33.19* contributed to the populations tested. Thus, within the limits of the resolution of these analyses, we conclude that if different Neandertal populations contributed to modern humans populations (21,22), these Neandertal populations were similarly related to the Neandertal genomes available to date.

### Small population size and inbreeding

The Neandertal genome from Denisova Cave, *Denisova 5*, carries a high proportion of long tracts that are homozygous by descent (HBD tracts) (3). Whereas tracts that are over 10cM long indicate that the parents of *Denisova 5* were closely related, HBD tracts between 2.5 and 10cM indicate that the population from which *Denisova 5* comes was of small size over approximately 100 generations before the individual lived (SI8).

Compared to *Denisova 5*, the *Chagyrskaya 8* genome carries fewer HBD tracts longer than 10cM, but more HBD tracts of intermediate length (SI8). In fact, all three high-coverage Neandertal genomes available carry more HBD tracts of intermediate size than almost all present-day and prehistoric modern human genomes, as well as the Denisovan genome (*Denisova 3*) (Fig. 3a). We show by coalescent simulations that this cannot be explained by an overall small but panmictic population. Rather, it suggests that Neandertal populations were subdivided (SI8).

**Figure 3:**
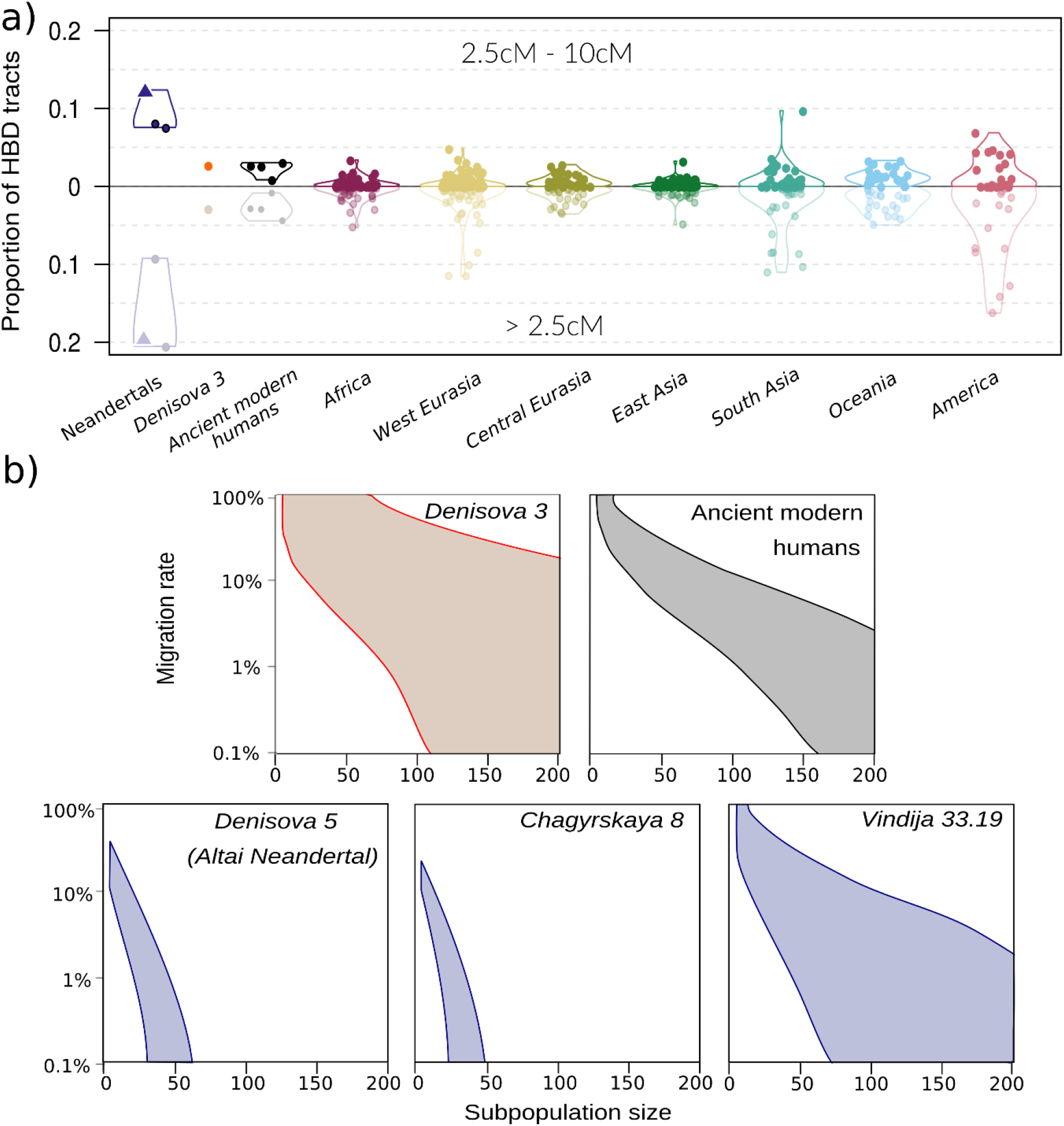
Homozygous tracts in archaic genomes. (a) Proportion of the genome spanned by HBD tracts of size 2.5 to 10 cM (dark – top panel) and all HBD tracts (light - bottom panel) for archaic genomes, ancient modern humans (“modern humans”) and present-day humans from the Simons Genome Diversity Project^18^. Values for *Chagyrskaya 8* are indicated by a triangle. The genetic size of the tracts was estimated assuming a uniform recombination rate of 1.3×10^−8^ recombinations/bp. (b) Estimated group size (x-axis) and migration rate (y-axis) for three Neandertal, one Denisovan and ancient modern human genomes based on HBD tracts >2.5cM long. The colored areas delimit 95% confidence intervals of likelihood ratios. Ancient modern humans give estimates for the genomes of four individuals dated to between ~45 kya and ~8 kya (“Ust’Ishim” (14), “Sunghir 2” (33), “Loschbour” (37) and “LBK-Stuttgart” (37)).

By coalescent modelling, we infer that *Chagyrskaya 8* and *Denisova 5* may have lived in subpopulations of 60 or fewer individuals. In contrast, current and past modern human populations as well as Denisovans (based on the *Denisova 3* genome) lived in subpopulations of more than 100 individuals, assuming a migration rate between subpopulations of 1% or less (Fig. 3b; SI8). Interestingly, the *Vindija 33.19* Neandertal seems to have lived in a subpopulation of larger size than the two Siberian Neandertals, although this difference is only marginally statistically significant when the proportion of the genomes covered by all HBD tracts longer than 2.5cM is considered (likelihood-ratio test, p=0.05).

### Derived genomic features in Neandertals

We used the three high-coverage Neandertal genomes to identify biological pathways where protein-coding genes show more derived non-synonymous substitutions fixed in the three Neandertals than expected from the silent and polymorphic changes. We identified 993 fixed non-synonymous substitutions among 889 genes, and 2,952 polymorphic non-synonymous substitutions in the three Neandertals.

No groups of genes associated with a specific known biological functions or phenotypes (23) show a higher ratio of non-synonymous to synonymous fixed changes relative to the ratio of non-synonymous to synonymous polymorphic changes (MacDonald-Kreitman ratio)(24), compared to other groups of genes (25) (FWER>0.1)(SI9).

However, when analyzing genes preferentially expressed in different brain regions according to the Allen Brain Atlas (26,27), we find that genes expressed in the striatum in individuals 12-19 years of age show a higher MacDonald-Kreitman ratio (1.02, FWER =0.029) than genes expressed in other brain regions and at other ages (0.53-0.83) (SI9). This may indicate that proteins encoded by these genes have been the targets of positive selection or evolved under relaxed constraints on the Neandertal evolutionary lineage. In addition, genes expressed in the prenatal striatum carry more substitutions in their untranslated regions than genes expressed elsewhere (FWER=0.049) and at other times. Among genes expressed in the striatum, those carrying fixed non-synonymous changes in Neandertals are more often present in genomic regions that carry little or no DNA introgressed from Neandertals in present-day humans than striatal genes not carrying such changes (Fisher’s exact test p=0.026). This pattern is not observed for all genes carrying fixed non-synonymous substitutions in the Neanderals (p>0.1), suggesting that some substitutions in Neandertal striatal genes might have been negatively selected in modern humans. Besides the striatum, genes expressed prenatally in the posterior parietal cortex, in the ventrolateral prefrontal cortex and in the primary somatosensory cortex carry more fixed substitutions in their regulatory regions in Neandertals than genes expressed in other brain regions and at other times (SI9).

In a further attempt to detect positive selection along the Neandertal lineage, we performed the Hudson, Kreitman, Aguadé (HKA) test (28) and the population branch statistic (PBS) (29) in 25kb sliding windows across the genome (SI10). We estimated the probability of obtaining the observed values of the different statistics by coalescent simulations, and retained windows with a false discovery rate <5% (Fig. 4). We identify a total of 35 separate candidate regions. One of these candidates, a 75kb long regions identified by PBS on chromosome 5, overlaps two separate windows identified by HKA. This overlap is lower than if expected by change (p-value<4×10^−4^, SI10). The candidate regions identified overlap genes involved in neural development (*EXOC6B*), immunity and wound healing (*HTN1, EVPLL*) and mitochondrial functions (*NSUN3, TIMM29*). For both statistics, we find an overlap with genomic regions previously identified as positively selected in modern humans (30) (enrichment test, p-values=0.010 and 0.056 for HKA and PBS, respectively).

**Figure 4:**
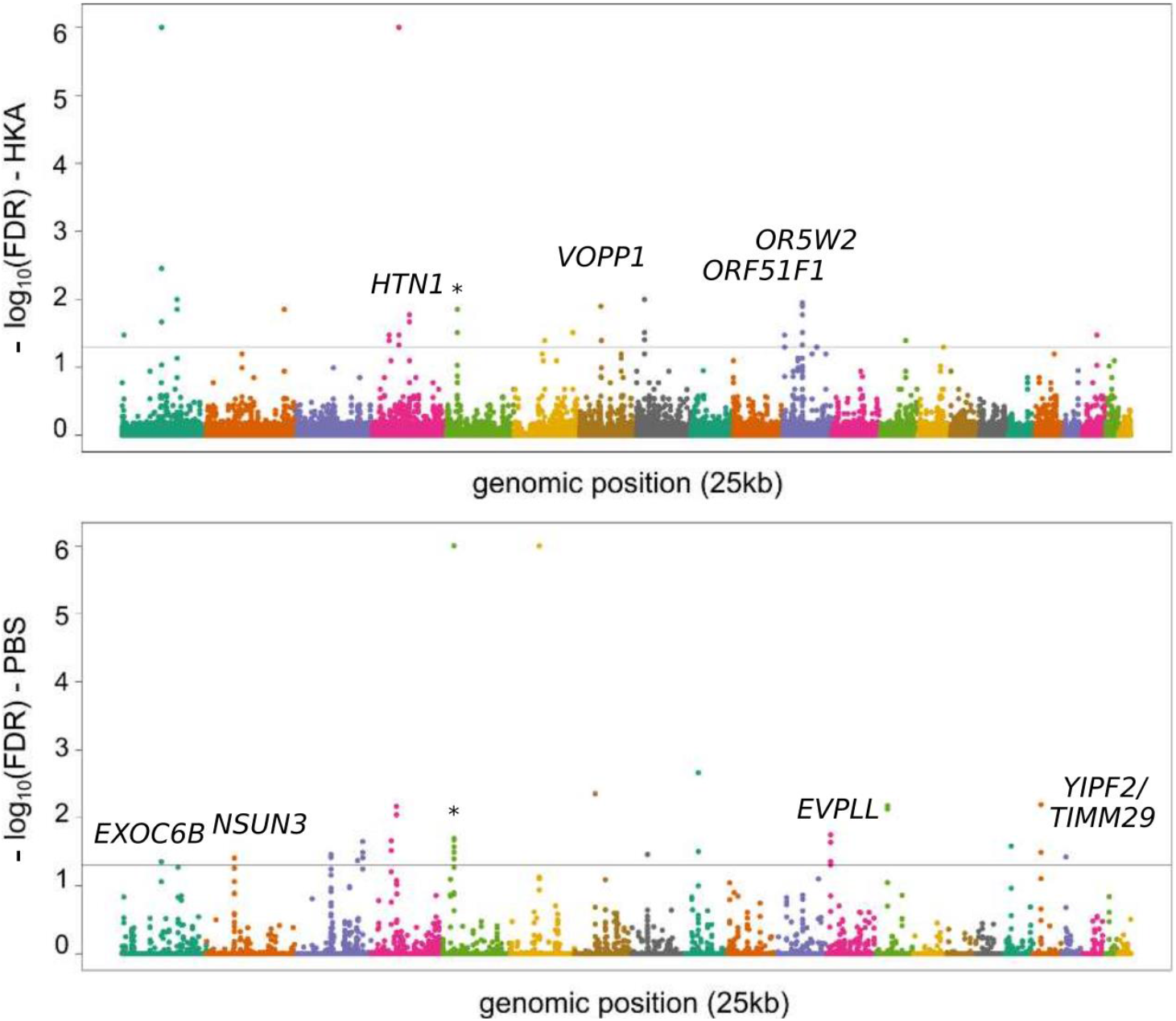
Positive selection on the Neandertal lineage. Manhattan plots of HKA (top) and PBS (bottom) tests for positive selection applied to three Neandertal genomes. Different colors indicate different autosomes, from 1 (left) to 22 (right). The y-axis indicates −log10 of false discovery rates (FDR). The gray line indicates FDR=5%. Candidate windows that overlap exons are indicated by the name of the corresponding gene. The regions that are significant for both tests are indicated by an asterisk.

## Discussion

*Chagyrskaya 8* is more closely related to *Vindija 33.19* and other late Neandertals in western Eurasia than to the *Denisova 5* Neandertal who lived earlier in the Altai Mountains (Fig. 2). *Chagyrskaya 8* is thus related to Neandertal populations that moved east sometime between 120 and 80 thousand years ago (8,31). Interestingly, the artifacts found in Chagyrskaya Cave show similarities to artifact assemblages in Central and Eastern Europe (10) (SI1), suggesting that Neandertal populations coming from western Eurasia to Siberia may have been brought their material culture with them (10,32). Some of these incoming Neandertals encountered local Denisovan populations, as shown by *Denisova 11*, who had a Denisovan father and a Neandertal mother related to the population in which *Chagyrskaya 8* lived.

In this regard, it is interesting that the *Chagyrskaya 8* and *Denisova 5* Neandertals lived in smaller populations than the *Vindija 33.19* Neandertal in Croatia, the Denisovan *Denisova 3*, and modern humans (33) (Fig. 3b). Neandertals in the Altai region may have lived in smaller and more isolated populations than Neandertals elsewhere as that region represented the periphery of their geographical distribution and may have been an area where Denisovans were more continuously present. However, more detailed studied of the Middle and Upper Paleolithic population history of Denisova Cave and other sites will be necessary to clarify this.

When analyzing genetic changes on the Neandertal lineage using the three Neandertal genomes, the number of changes in genes expressed in the striatum during adolescence stand out. Furthermore, genes expressed in the striatum overlap more frequently than expected with genomic regions where Neandertal introgressed fragments in modern human genomes are rare. We speculate that striatal genes may carry Neandertal-specific adaptation or other changes that were disadvantageous when introduced into modern humans. This, as well as positive selection for derived changes in the modern human lineage, may underlie so-called “Neandertal deserts” in present-day human genomes, *i.e*. regions that are depleted of Neandertal ancestry (19,34).

As more high-quality Neandertal genomes become available it will be possible to more comprehensively explore genes and groups of genes that carried functionally relevant changes in Neandertals. Currently, there is suggestive evidence that such findings may be forthcoming. For example, in addition to genes expressed in the striatum, untranslated regions and promoters of genes expressed in the posteroventral (inferior) parietal cortex, a brain region that has been associated with speech and mathematical cognition (35), carry more changes in the three Neandertals than expected by chance. In addition, among the top phenotypes associated with changes in regulatory regions in Neandertals are abnormalities in parts of the skeleton where Neandertal morphology stand out, such as the nasal bridge and the rib cage (36).

## Acknowledgments

This study was funded by the Max Planck Society; the European Research Council (grant agreement no. 694707 to S.P); and the Russian Science Foundation (project No. 14-50-00036 to A.P.D.). We thank Sarah Nagel, Birgit Nickel, Barbara Schellbach and Antje Weihmann for laboratory work; and Heiko Temming for CT scans.

## Data availability statement

The genome sequence of *Chagyrskaya 8* can be downloaded from http://ftp.eva.mpg.de/neandertal/Chagyrskaya/VCF/. Its mitochondrial DNA genome sequence has been deposited in GenBank (accession ID MK388903).

## Notes

http://ftp.eva.mpg.de/neandertal/Chagyrskaya/VCF/

