## Supplementary Information for "A high-coverage Neandertal genome from Chagyrskaya Cave"

### Supplementary Information 1

#### Archaeological context and morphology of *Chagyrskaya 8*

##### Archaeological, spatial and stratigraphical context of *Chagyrskaya 8*

Chagyrskaya Cave (51°26’32.99’’, 83°09’16.28’’) is located in the mountains of northwestern Altai, on the left bank of the Charysh River (Tigirek Ridge) (Fig. S1.1, A). The cave faces north and is situated at an elevation of 373 m ASL, 19 m above the river. The cave is relatively small and consists of two chambers, spanning a total area of 130 $m^2$ (Fig. S1.1, A, D). Investigations started in 2007 and approximately 35 $m^2$ have been excavated since that time by two teams (Fig. S1.1, D) (1,2).

The stratigraphic sequence contains Holocene (layers 1–4) and Pleistocene layers (5-7). The Neandertal artifacts and anthropological remains are associated with layers 6a, 6b, 6c/1 and 6c/2 (Fig. S1.1, E). Among the Upper Pleistocene layers, only the lower layer 6c/2 is preserved *in situ*, while the upper layers with Neandertal lithics have been re-deposited from that lower layer via colluvial processes (2). Additional evidence of re-deposition is seen in the size sorting of the paleontological remains and bone tools, with the largest bones found in the lower layers 6c/1 and 6c/2, and the smallest bones in the upper layers 5, 6a and 6b (1,2).

The *Chagyrskaya 8* specimen was found during wet-sieving of layer 6b (horizon 2) in the 2011 field expedition headed by Sergey Markin. Using field documentation, we reconstructed the spatial (Fig. S1.1, B, C) and stratigraphic (Fig. S1.1, E, F) location of the specimen. The spatial data shows that *Chagyrskaya 8* had been surrounded by lithic artifacts and paleontological remains which are the result of the re-depositional process (Fig. S1.1, B, C). The age of the layer 6b sediments has been estimated by post-infrared IRSL to 53.4 ± 3.3 ky. All optical dates obtained from Chagyrskaya Cave sediments correspond to the end of MIS 4 and the beginning of MIS 3 (1,2).

The sum total of palaeontological material obtained from horizon 2 of layer 6b is 7,520 items, most of them strongly fragmented. The faunal assemblage includes bones of bison, the Ovodov horse, reindeer, argali and Siberian ibex (1).

The total number of lithic artifacts from layer 6b horizon 2 (2011 excavations) is 1,362 items; 83,5% of them are debris and chips. No cores have been found. The primary knapping process has been characterized based on the dorsal scar pattern of flakes (radial and orthogonal knapping). Most of the blanks in the assemblage are flakes. A total of 32 tools have been recovered, including mostly single-edge longitudinal scrapers, convergent scrapers, retouched points and plano-convex bifacial scraper (Fig. S1.2, 1-8). The assemblage from layer 6b horizon 2 has the same typological and technological characteristics as the larger assemblage from layer 6c/1 that has been studied in detail. In general, the lithic assemblage from Chagyrskaya Cave is characterized by radial/orthogonal core reduction methods, plano-convex/plano-convex alternate bifacial methods and the domination of trapezoidal and leaf-shaped convergent scrapers. Bifacial tools include backed points and scrapers of the Keilmesser type (3). All characteristics are typical of the Micoquian assemblages described for Central and Eastern Europe. Attributive analysis of lithic assemblages demonstrated the complete sequence of raw material treatment (4) typical for a base consumption camp. The available zooarchaeological and paleontological data confirm this function (5). The Chagyrskaya Cave assemblage differs from the Denisova Middle Paleolithic (3,5,6) assemblages (located ~ 100 km away) while it is similar to the Okladnikov assemblage (~ 80 km away) (Fig. S1.1, A).

**The morphology of *Chagyrskaya 8***

*Chagyrskaya 8* is a distal manual phalanx of an adult individual based on its size and the fully fused epiphysis (Fig.S1.3). It is not possible to assign the specimen securely to ray or side, but its general morphology and size allows to exclude an attribution to the thumb. Neandertal distal phalanges of the fifth ray usually show a marked narrowing at the midshaft and small apical tufts, and neither of these features are visible here, so an attribution to ray 2-4 seems likely. Using the techniques of Case and Heilman (7), an assignment to the left side is possible, but as the accuracy of these methods is only between 50 and 70%, this is very tentative.

The specimen is complete and shows very good surface preservation (Fig. S1.3). The shaft is robust (4.7x6.9mm, dorsopalmar x radioulnar at the midshaft) and long (max. length 19.7 mm, articular length 18.7 mm). The apical tuft is large, round and radioulnarly expanded (maximum breadth 9.4), and is in lateral view somewhat offset towards palmar from the long axis of the shaft and quite thick (distal max. height 5.6 mm). The insertion of the M. *flexor digitorum profundus* in the proximal half of the shaft is roughly triangular, and about 6.3x4.5 mm (radioulnar x proximodistal). The interphalangeal articular surface is large (9.6x4.7 mm, proximal articular breadth x proximal articular height, 11.5x7.2 mm proximal max. height x breadth).

Metrical comparisons (Table S1.1) show that *Chagyrskaya 8* is relatively long compared to recent modern human phalanges, both with regards to its maximum and articular length, while being somewhat below the average for Neanderthals. Its midshaft height is close to the upper limit of the variability seen in modern humans, and its large midshaft breadth is outside the range of variation seen in either recent or fossil modern humans, but close to the Neanderthal average. The proximal articular surface is quite wide compared to modern humans. The apical tuft is very large, with a width and dorsopalmar thickness well above the range seen in recent modern humans.

We performed a principal component analysis of a subset of the measurements reported in Table S1.1, selected to maximize the available sample size (*Articular length, midshaft height and breadth, proximal maximum height and breadth, proximal articular height and breadth, distal maximum breadth*). The first two principal components explain about 90% of the variance, and separate modern humans and Neandertals quite clearly (Figure S1.4). PC1 is mostly driven by distal maximum breadth, articular length and proximal maximum breadth. Chagyrskaya 8 falls in the Neandertal group, closely clustering with the other two Chagyrskaya specimens, Krapina 206.5 and Moula-Guercy M-H1-21.

The wide and rounded apical tuft, combined with a long and robust shaft, as seen in *Chagyrskaya 8* has been since a long time recognized as a morphology characteristic for Neandertals (8-10). A similar morphology is seen in at least some of the Atapuerca SH phalanges (11), but interestingly not in the Denisovan distal phalanx *Denisova 3* (12). All in all, the morphology and metric features of *Chagyrskaya 8*, clearly support its attribution to Neandertals.

**Table S1.1.** Metric comparisons of *Chagyrskaya 8* to 2nd to 4th distal phalanges of recent and fossil modern humans, and Neanderthals.

| Group | Maximum length[a] | Articular length | Midshaft height | Midshaft breadth | Proximal maximum height | Proximal maximum breadth | Proximal articular height | Proximal articular breadth | Distal maximum breadth | Distal maximum height |
|---|---|---|---|---|---|---|---|---|---|---|
| Recent[b] | 17.34+-1.88 (n=31; 14.2-20.7) | 16.32+-1.75 (n=31; 13.5-19.4) | 3.53+-0.58 (n=31; 2.6-4.9) | 4.41+-0.76 (n=31; 3.1-6.0) | 6.16+-0.64 (n=31; 5.1-7.2) | 9.63+-1.14 (n=31; 7.7-12.2) | 5.11+-0.57 (n=31; 4.3-6.3) | 8.51+-0.90 (n=31; 7.1-10.7) | 6.83+-1.03 (n=31; 4.9-9.0) | 3.40+-0.55 (n=31; 2.5-4.8) |
| Fossil MH[c] | 19.74+-2.08 (n=5; 17.2-22.9) | 17.13+-1.90 (n=15; 14.0-22.6) | 3.84+-0.67 (n=15; 2.7-5.2) | 5.03+-0.84 (n=15; 4.1-6.6) | 6.45+-0.83 (n=13; 5.4-7.8) | 10.44+-0.95 (n=13; 9.0-11.8) | 5.69+-1.03 (n=12; 3.8-7.3) | 8.68+-1.10 (n=13; 7.0-10.6) | 6.86+-1.50 (n=14; 5.2-9.6) | No data available |
| Neandertals[d] | 20.02+-1.14 (n=16; 18.2-22.9) | 19.60+-1.63 (n=29; 17.2-23.2) | 4.19+-0.43 (n=28; 3.4-5.0) | 6.42+-0.67 (n=28; 5.0-7.7) | 7.01+-0.87 (n=28; 5.5-8.8) | 11.98+-1.05 (n=28; 10.0-14.1) | 5.66+-0.53 (n=23; 5.0-6.8) | 10.13+-0.94 (n=23; 8.5-11.9) | 9.95+-1.24 (n=29; 7.8-13.9) | 4.35+-0.75 (n=10; 3.2-5.5) |
| Chagyrskaya 8 | 19.7 | 18.7 | 4.7 | 6.9 | 7.2 | 11.5 | 4.7 | 9.6 | 9.4 | 5.6 |

[a] Mean+- standard deviation, (n, range)

[b] Own measurements on modern human phalanges (ray II-IV), collection of Dept. of Anthropology, U. of Vienna and U. of Toronto

[c] Specimens included: Dolni Vestonice DV3, 13, 15, 16 (13), Tianyuan 1 (14), Sunghir 1 (15), Sea Harvest PQ-S1708 (16)

[d] Specimens included: Krapina 206.1-6, 8-11 (17), Kebara (18), Shanidar 3,4,5,6 (19), Palomas 28, 96HH, 92Y, 96BBB, 96CCC (20), Moula Guercy M-K0-HNN1, M-H1-21(21), Combe Grenal 26 (22), Chagyrskaya 56b & 61 (own data)

**Additional hominin remains spatially associated with *Chagyrskaya 8***

All in all, 75 human remains have been recovered until now from Chagyrskaya Cave. Most of the remains (61 of the 75 total remains) derive from two clusters, a Southern cluster (SC) in the vicinity of square N10 (N=31) and a Northern cluster (NC) around squares K6, K7 and L6 (N=30).

*Chagyrskaya 8* is part of the SC, and like most of the material there derives from layer 6b. The remains from layer 6b of the SC include a partial left foot consisting of the 3rd, 4th and 5th metatarsals (*Chagyrskaya 52* a,b and 53), a calcaneus (*Chagyrskaya 36*), a medial cuneiform (*Chagyrskaya 27*), a middle pedal phalanx (*Chagyrskaya 22*) and the distal ends of tibia and fibula (*Chagyrskaya 24* a&b). Four vertebrae, including the C1 (*Chagyrskaya 37*), C2 (*Chagyrskaya 38*), L5 (*Chagyrskaya 35*) and a middle thoracic vertebra (*Chagyrskaya 26*), as well as rib and sternum fragments (*Chagyrskaya 30 & 54*) are also part of the cluster, and could be associated with the foot. Several hand remains were also found in this area. These include a third metacarpal (*Chagyrskaya 23*), an associated middle and distal phalanx (*Chagyrskaya 56 b* & *c*) and a distal thumb phalanx fragment (*Chagyrskaya 56 a*).

Based on the dental remains, at least three individuals are represented in layer 6b of the SC, and at least two in 6c/1. *Chagyrskaya 6*, a partial right mandible of a young adult preserving C to $M_2$, and *Chagyrskya 14*, a left lower I2 are likely associated. The left maxilla fragment *Chagyrskaya 57*, preserving $M^1$ and $M^2$ is too worn to belong to the same individual. An isolated left upper $M^1$ (*Chagyrskaya 10*), duplicates the $M^1$ of *Chagyrskaya 57*, and is also more worn than would be expected for *Chagyrskaya 6*, and thus likely represents a third individual.

The underlying Layer 6c/1 yielded several hominin fragments. A naturally exfoliated left deciduous $dm^1$ (*Chagyrskaya 18*) and a right upper deciduous canine (*Chagyrskaya 20),* also naturally exfoliated could represent the same individual. A left lower $P_3$, *Chagyrskaya 12* is more worn than *Chagyrskaya 6*, but could belong to the same individual as either *Chagyrskaya 57* or Chagyrskaya 10.

It is not completely clear with which of these elements (or whether any) *Chagyrskaya 8* is associated with. Based on preliminary ancient DNA analyses, the left lower $P_3$ *Chagyrskaya 12*, found in the neighboring square, but in a lower horizon (Sq. M10, layer 6c/1) than *Chagyrskaya 8* (Sq. N10, layer 6b), may be the same individual. This association of remains from two different layers is in agreement with the geological data indicating some reworking of the cave deposits, and the redeposited origin of horizons 6a, b, and c/1.

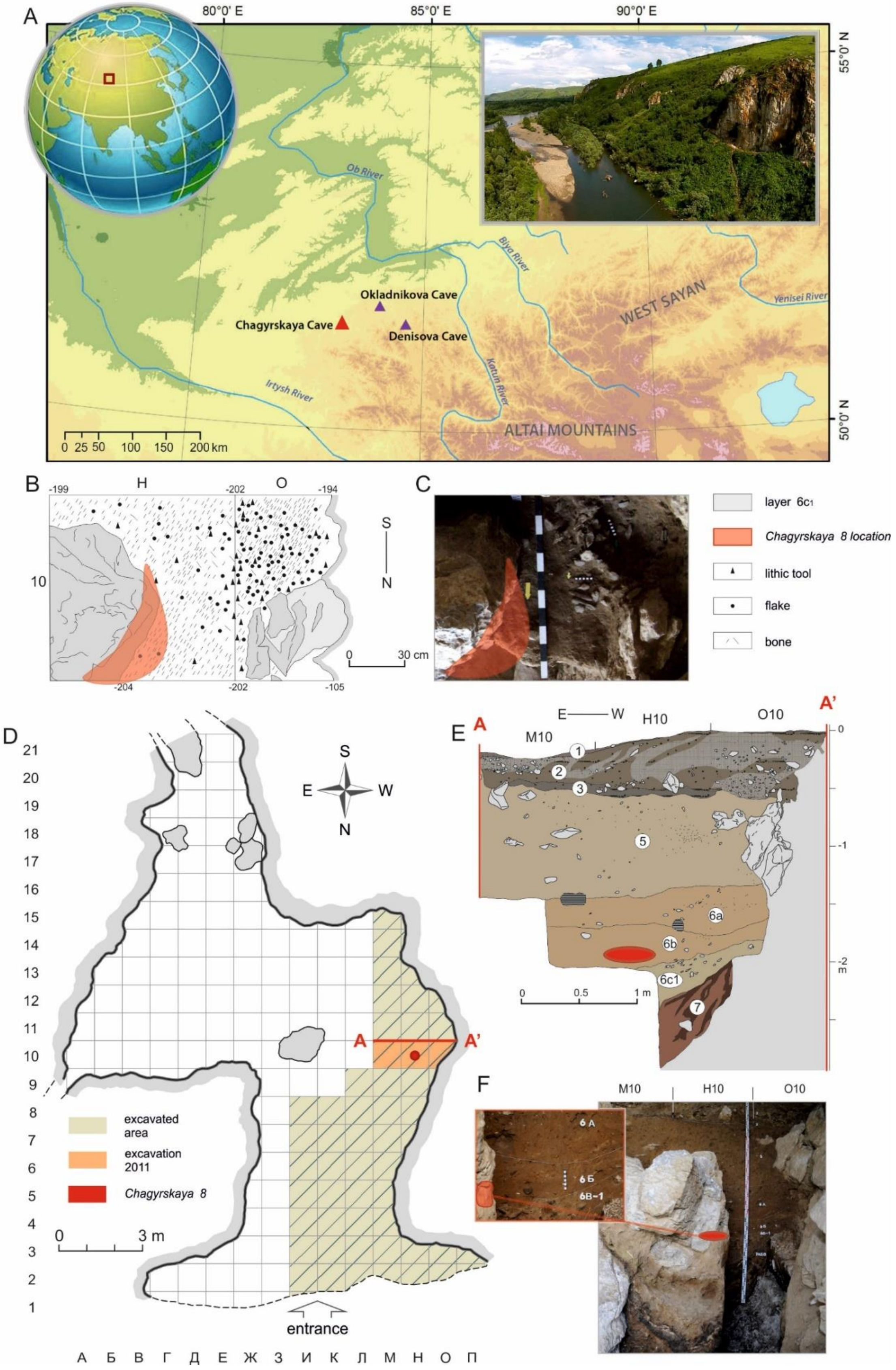
A
80°0' E
85°0' E
90°0' E
55°0' N
50°0' N
Ob River
Biya River
Okladnikova Cave
Chagyrskaya Cave
Denisova Cave
WEST SAYAN
Yenisei River
Irtysh River
Katun River
ALTAI MOUNTAINS
0 25 50 100 150 200 km
B
C
H
O
-199
-202
-194
10
-204
-202
-105
S
N
0 30 cm
layer 6c1
Chagyrskaya 8 location
lithic tool
flake
bone
D
S
E
W
N
A
A'
excavated area
excavation 2011
Chagyrskaya 8
0 3 m
entrance
А Б В Г Д Е Ж З И К Л М Н О П
E
E — W
M10
H10
O10
6a
6b
6c1
0 0.5 1 m
F
M10
H10
O10
6 А
6 Б
6В-1

**Figure S1.1.** Chagyrskaya Cave (A) Location of the Middle Paleolithic cave sites mentioned in the article and view of Chagyrskaya Cave. (B, C) Spatial location of the *Chagyrskaya 8* specimen. (D) Plan of the Cave chambers showing the location of the 2011 excavation area and the location of *Chagyrskaya 8*. (E, F) Stratigraphic location of *Chagyrskaya 8.*

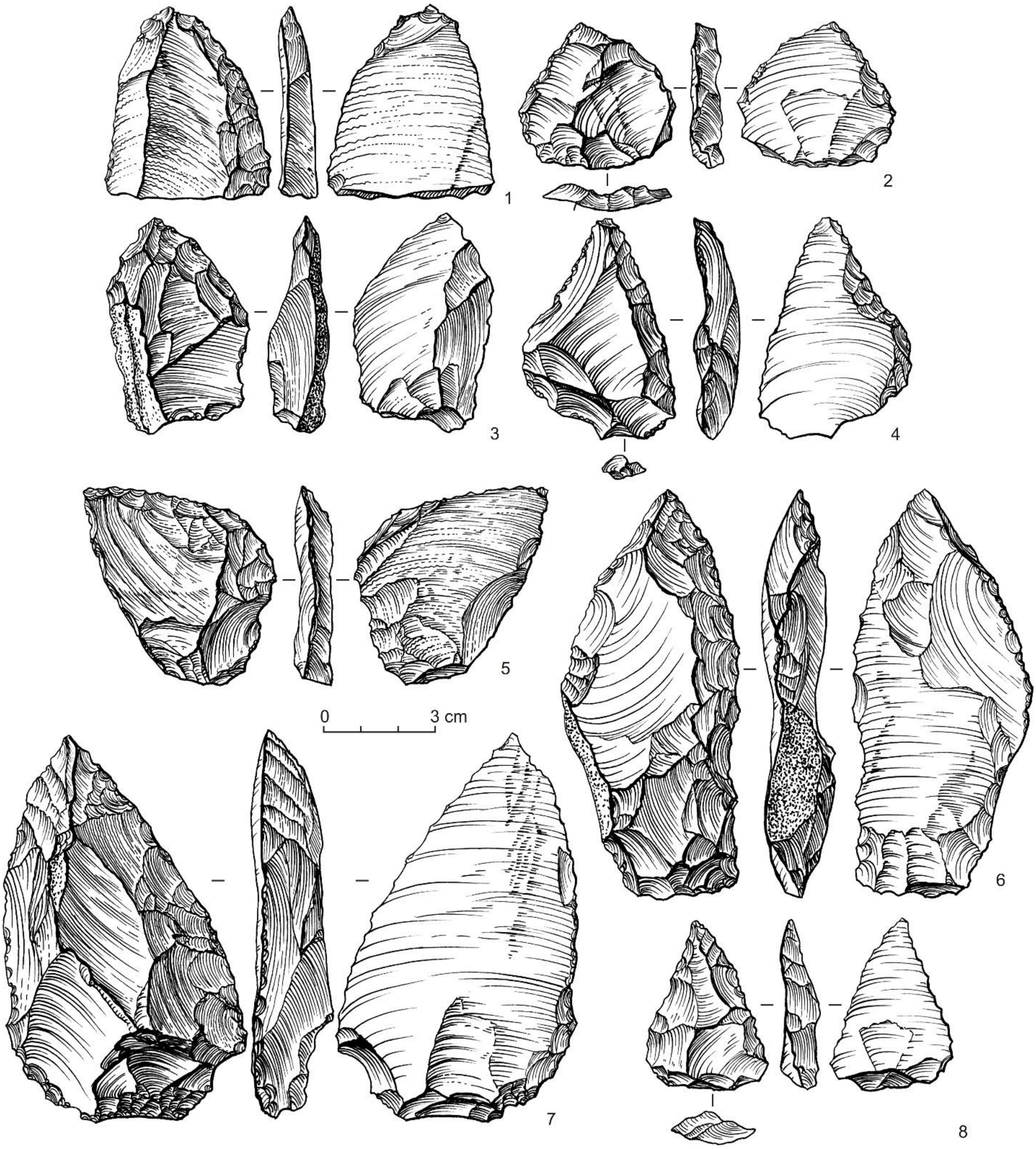


**Figure S1.2.** Archaeological context of *Chagyrskaya 8*: lithic tools from layer 6b (horizon 2).

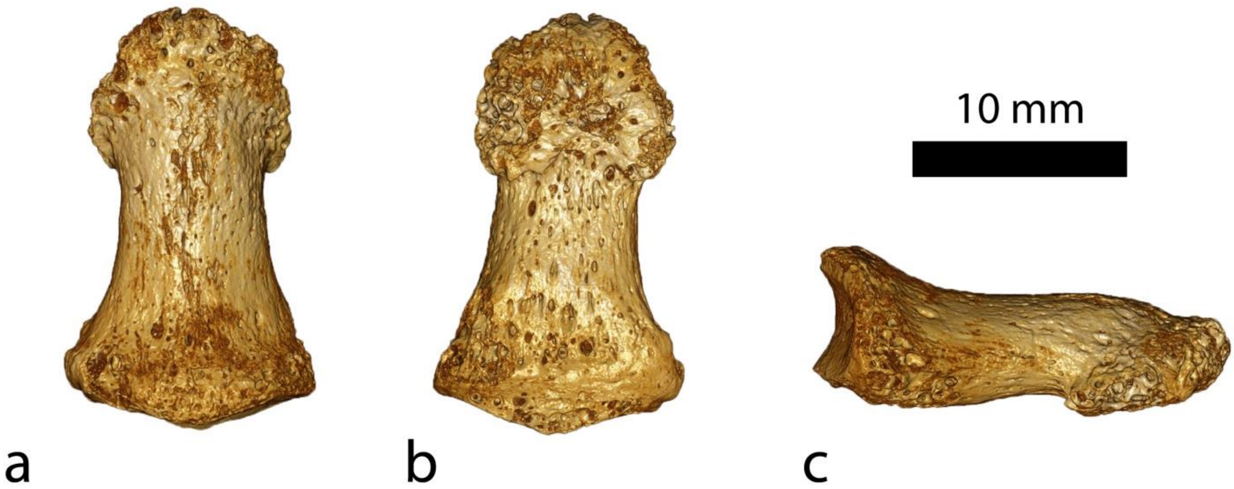


**Figure S1.3.** The *Chagyrskaya 8* phalanx (µCT based renderings). a: dorsal view, b: palmar view, c: lateral view (probably radial)

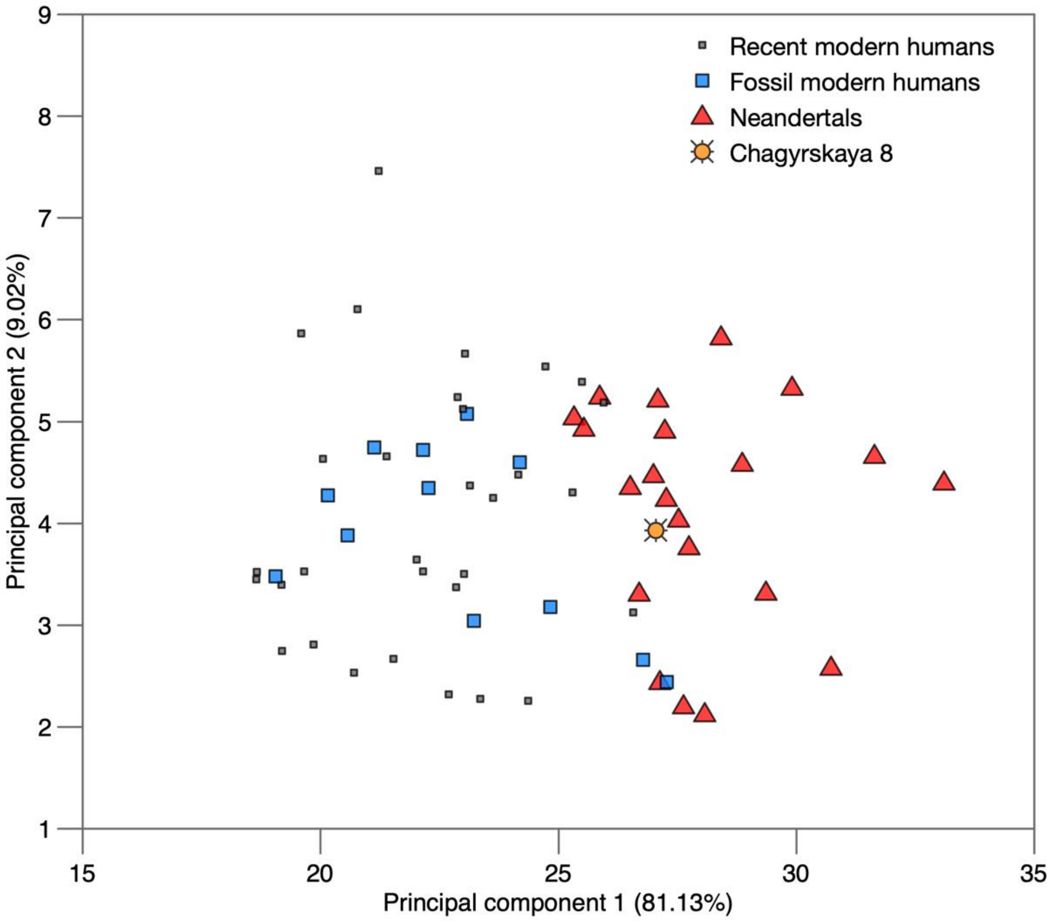

**Figure S1.4.** Principal component analysis of modern human and Neandertal distal phalanges. Data as in Table S1.1, measurements used are: articular length, midshaft height and breadth, proximal maximum height and breadth, proximal articular height and breadth, distal maximum breadth.

### Supplementary Information 2

#### DNA extraction and sequencing

##### Sampling, treatment of bone powder and DNA extraction

The *Chagyrskaya 8* specimen was µCT-scanned at the MPI-EVA prior to sampling. The proximal base of the phalanx was sampled in a clean room dedicated for ancient DNA work using 1.0 mm disposable dentistry drills (models H1S-010-RA and H1SX-010-RA, NTI-Kahla, Germany). Following the abrasion of the external surface of the bone, a total of 111 mg of bone powder was collected on two occasions. Of this, three aliquots of ~30 mg of bone powder each were treated with 0.5% sodium hypochlorite solution and one aliquot of ~15 mg was treated with 1.0% sodium hypochlorite (1). Three consecutive washes with water were carried out to remove the sodium hypochlorite from the bone pellet (1). DNA was extracted as described in (2), with modifications (1) and eluted in 50 µl TET buffer. One extraction negative control (no bone powder) was carried along each experiment.

##### DNA library preparation and sequencing

Aliquots of 5, 10 or 15 µl from the DNA extracts were converted into single-stranded DNA libraries as described in (3) with modifications (1). A short artificial oligonucleotide was spiked into each library to assess the efficiency of the library preparation (4). Each experiment included at least one library preparation negative control. The number of DNA molecules and spiked-in control oligonucleotides in the libraries were determined by digital droplet PCR (5) or quantitative PCR (3,4). Each library was tagged with a pair of unique indices (1,6). Following an amplification into plateau using AccuPrime Pfx DNA polymerase (Life Technologies) (7), purification was carried out using the MinElute PCR purification kit (Qiagen), or on an automated liquid handling platform (Bravo NGS, Agilent Technologies) using SPRI beads (8) as described elsewhere (9). The DNA content in each amplified indexed library was quantified using a NanoDrop ND-1000 (NanoDrop Technologies) photospectrometer. Table S2.1 shows a summary of the extracts and libraries generated for *Chagyrskaya 8*. Enrichment for human mitochondrial (mt) DNA fragments (10) was carried out in one round of hybridization capture on beads (11) as modified in (9), from 1 µg of library L5329. Libraries were sequenced either individually or in pools with other libraries originating from this or other projects, and an indexed ΦX 174 library was spiked into each sequencing run. Paired-end sequencing was carried out by 76 cycles (6) on Illumina platforms (HiSeq 2500 or MiSeq) (Table S2.2).

**Table S2.1. List of DNA extracts and libraries prepared from the *Chagyrskaya 8* specimen.** The extraction and library preparation negative controls are marked in gray.

| | Extract ID | *Bone powder [mg]* | *Sodium hypochlorite [%]* | *Input in library [µl]* | *Number of molecules in library* | *Number of control oligo in library* | *Amplified indexed library ID* | *Library enriched for human mtDNA* |
|---|---|---|---|---|---|---|---|---|
| **Chagyrskaya 8** | E2894 | 29.8 | 0.5 | 15 | 9.10E+08 | 5.55E+05 | L5329 | L5284 |
| | | | | 5 | 3.69E+08 | N/A | L5727 | - |
| | | | | 5 | 3.57E+08 | N/A | L5728 | - |
| | | | | 5 | 9.43E+08 | 7.45E+05 | L5831 | - |
| | | | | 5 | 4.88E+08 | 6.35E+05 | L5832 | - |
| | | | | 5 | 5.18E+08 | 7.10E+05 | L5833 | - |
| | | | | 5 | 6.70E+08 | 8.10E+05 | L5848 | - |
| | | | | 5 | 6.75E+08 | 7.30E+05 | L5849 | - |
| | E3658 | 27.3 | 0.5 | 10 | 1.22E+09 (*) | 2.44E+04 (*) | L5729 | - |
| | | | | 5 | 2.74E+09 | 8.05E+05 | L5800 | - |
| | | | | 5 | 2.62E+09 | 6.85E+05 | L5819 | - |
| | | | | 5 | 1.64E+09 | 6.70E+05 | L5820 | - |
| | | | | 5 | 2.38E+09 | 7.70E+05 | L5821 | - |
| | E3659 | 27.9 | 0.5 | 10 | 8.41E+08 (*) | 4.05E+04 (*) | L5730 | - |
| | | | | 5 | 1.34E+09 | 7.55E+05 | L5830 | - |
| | | | | 5 | 1.47E+09 | 6.95E+05 | L5822 | - |
| | | | | 5 | 1.01E+09 | 7.40E+05 | L5823 | - |
| | | | | 5 | 1.74E+09 | 7.30E+05 | L5824 | - |
| | | | | 5 | 1.83E+09 | 9.00E+05 | L5850 | - |
| | | | | 5 | 2.05E+09 | 6.95E+05 | L5851 | - |
| | E4972 | 14.0 | 1.0 | 10 | 1.14E+09 | 7.70E+05 | L5794 | - |
| ***Extraction negative controls*** | E2904 | - | 0.5 | 15 | 6.35E+07 | 3.95E+05 | L5267 | L5294 |
| | E3663 | - | 0.5 | 10 | 1.83E+06 (*) | 3.69E+04 (*) | R5791 | - |
| | E4980 | - | 1.0 | 10 | 1.69E+07 | 8.15E+05 | L5795 | - |

| | | | | | | | | |
|---|---|---|---|---|---|---|---|---|
| ***Library preparation negative controls*** | - | - | - | 15 | 2.86E+07 | 5.70E+05 | L5270 | L5296 |
| | - | - | - | 10 | 2.54E+06 (*) | 5.92E+04 (*) | R5792 | - |
| | - | - | - | 10 | 1.30E+07 | N/A | R9888 | - |
| | - | - | - | 10 | 1.40E+07 | 7.70E+05 | L5797 | - |
| | - | - | - | 5 | 1.27E+07 | 8.75E+05 | L5803 | - |
| | - | - | - | 5 | 4.15E+06 | 8.40E+05 | L5826 | - |
| | - | - | - | 5 | 1.18E+07 | 8.10E+05 | L5827 | - |
| | - | - | - | 5 | 1.12E+07 | 7.45E+05 | L5845 | - |

(*) – indicates that the quantification was carried out by quantitative PCR.

**Table S2.2. List of sequencing runs performed to generate data from *Chagyrskaya 8.*** Libraries were sequenced on Illumina platforms, either HiSeq 2500 or MiSeq. Libraries originating from negative controls are noted in parentheses, and the base-calling algorithm used is indicated.

| *Libraries* | *Sequencing platform* | *Number of lanes* | *Base caller* |
|---|---|---|---|
| *L5284 (L5294, L5296)* | MiSeq | 1 | Bustard |
| *L5329 (L5267, L5270)* | HiSeq 2500 | 1 | freeIbis |
| *L5329* | HiSeq 2500 | 4 | freeIbis |
| *L5729-L5730 (R5791-R5792)* | MiSeq | 1 | Bustard |
| *L5727-L5728 (R9888)* | HiSeq 2500 | 1 | Bustard |
| *L5727-L5730* | HiSeq 2500 | 19 | Bustard |
| *L5794, L5800, L5819-L5824, L5830-L5833 (L5795, L5797, L5803, L5826-L5827)* | HiSeq 2500 | 1 | Bustard |
| *L5329, L5830-L5833* | HiSeq 2500 | 17 | Bustard |
| *L5848-L5851 (L5845)* | HiSeq 2500 | 1 | Bustard |
| *L5848-L5851* | HiSeq 2500 | 8 | Bustard |

### Supplementary Information 3

#### Data Processing, Genotyping and General Filters

##### Initial processing

Data from 21 DNA sequencing libraries were used (Table S3.1). Base calling was carried out using freeIbis (1) or Bustard (Illumina) (Supplementary Information 2, Table S2.2) and adapters were removed and overlapping forward and reverse reads were merged by leeHom (2). Both adapters were required to perfectly match the expected combination from the double-indexing scheme (3) for sequences to be retained. Sequences were mapped to the human reference genome (hg19) with added decoy sequences (4) using BWA (5) with the parameters -n 0.01 -o 2 -l 16500 (6). Duplicates were removed separately for each library using bam-rmdup (https://github.com/mpieva/biohazard-tools/).

**Table S3.1: Average sequence length of merged sequences and genomic coverage in each of the *Chagyrskaya 8* library used.**

| Library | Sequence length | Coverage |
|---|---|---|
| L5329 | 73.3 | 3.49 |
| L5727 | 70.4 | 1.52 |
| L5728 | 71.4 | 1.67 |
| L5729 | 69.2 | 1.19 |
| L5794 | 62.7 | 0.03 |
| L5800 | 63.0 | 0.04 |
| L5819 | 62.9 | 0.03 |
| L5820 | 62.3 | 0.03 |
| L5821 | 62.8 | 0.03 |
| L5822 | 62.9 | 0.03 |
| L5823 | 63.7 | 0.04 |
| L5824 | 62.9 | 0.03 |
| L5831 | 63.8 | 3.64 |
| L5832 | 65.4 | 3.07 |
| L5833 | 64.7 | 2.95 |
| L5848 | 66.1 | 2.89 |
| L5849 | 64.1 | 2.90 |
| L5730 | 69.3 | 1.03 |
| L5830 | 64.7 | 1.89 |
| L5850 | 63.5 | 0.76 |
| L5851 | 63.8 | 0.37 |

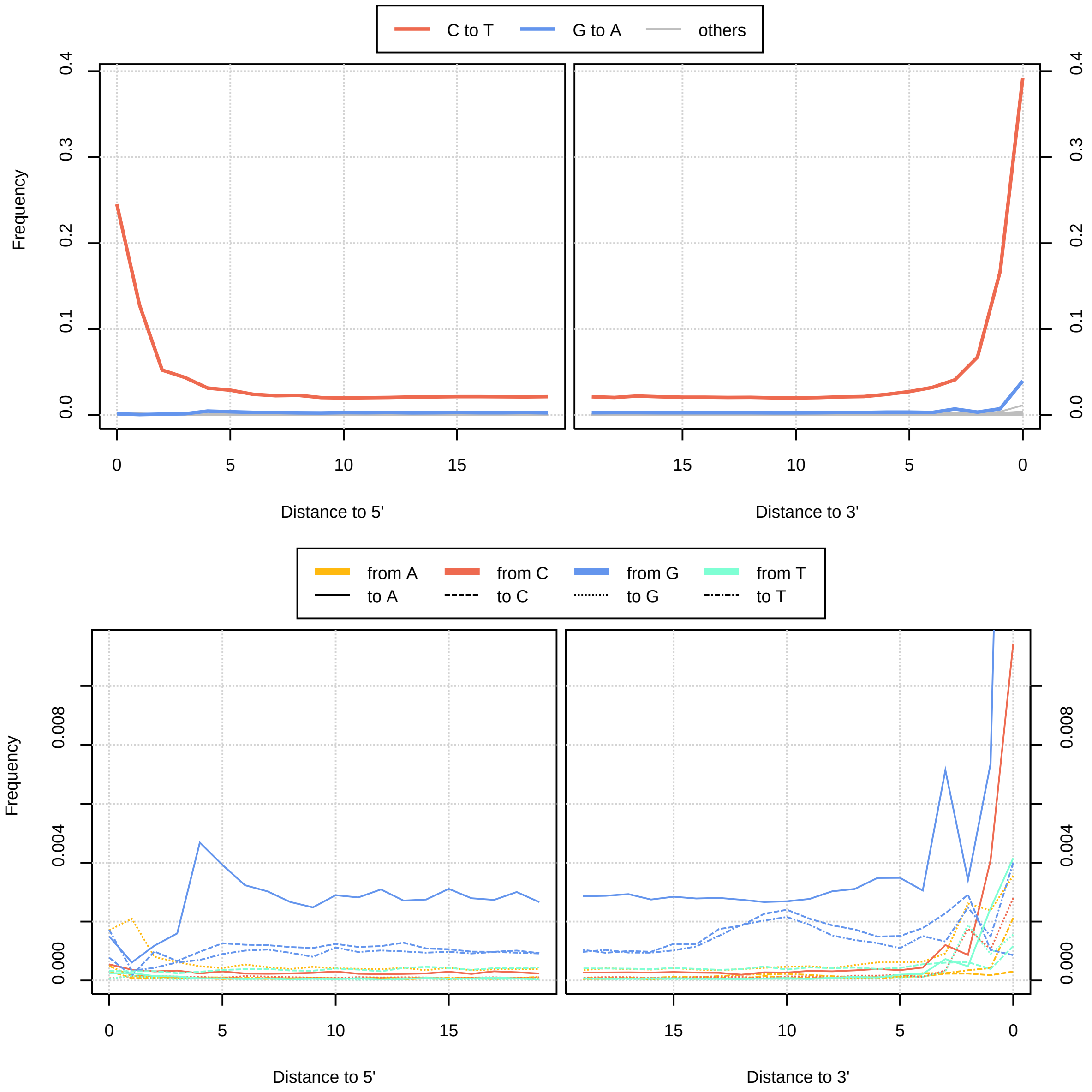


**Figure S3.1: Base substitutions in sequences used relative to the genome of *Denisova* 5.** Sequences from library L5831 are compared to the genotype calls of *Denisova* 5, and the frequency of substitution is shown on the y-axis. In the top panels the strongest substitution rate, C to T, are shown in comparison to the reverse complement substitution, G to A. Other substitutions, which occur at lower rates, are shown below at a finer scale. Sequences from other libraries show similar patterns.

Following previous approaches (4,7), sequences were excluded if they were shorter than 35bp and indels were realigned using GATK (version: 1.3-1.4) (8). Sequences of all libraries show patterns of ancient DNA damage in concordance with the age of the sample (9,10) (Fig. S3.1). The average length of sequenced DNA fragments is 67bp, ranging between 62 and 73bp among libraries (Table S3.1). The genomic coverage for each sequencing library is estimated by counting the number of bases with base quality ≥30 on autosomes that overlap regions of the human genome to which fragments of length 35 can be aligned uniquely (map35_100 track), divided by the total length of these regions. The average per base coverage is 27.6x. Table S3.2 shows the average coverage for each chromosome in regions of the genome retained by all filters. Chromosome X has a similar coverage to that of the autosomes, indicating that *Chagyrskaya 8* was a female.

**Table S3.2: Coverage and number of bases retained in regions passing all general filters.**

| Chromosome | Coverage (xfolds) | Bases retained |
|---|---|---|
| **1** | 27.2 | 136,520,807 |
| **2** | 26.8 | 150,617,216 |
| **3** | 26.9 | 124,517,822 |
| **4** | 27.4 | 117,549,348 |
| **5** | 27.1 | 111,404,161 |
| **6** | 28.0 | 105,405,915 |
| **7** | 27.7 | 90,715,332 |
| **8** | 26.9 | 90,679,087 |
| **9** | 26.4 | 68,750,317 |
| **10** | 26.4 | 80,138,252 |
| **11** | 26.2 | 81,131,730 |
| **12** | 26.6 | 79,848,567 |
| **13** | 27.5 | 61,892,869 |
| **14** | 26.6 | 54,694,181 |
| **15** | 26.6 | 47,827,596 |
| **16** | 25.6 | 44,383,122 |
| **17** | 25.0 | 43,493,333 |
| **18** | 27.0 | 49,143,097 |
| **19** | 25.0 | 25,470,168 |
| **20** | 26.6 | 38,417,872 |
| **21** | 26.7 | 21,155,819 |
| **22** | 26.1 | 19,487,080 |
| **X** | 27.5 | 80,456,000 |

#### Genotyping

Input files for the ancient DNA genotype caller snpAD (11) (version 0.2.1) were prepared using sequences that had a mapping quality of at least 25 and a length of at least 35 bases, and using bases with a quality of at least 30. Only positions within the 35mer mappability track were considered (12). Error rates and genotype frequencies were estimated independently for each chromosome (Table S3.3). These parameters were then used to call the most likely genotype at each site.

**Table S3.3: Estimates of genotype frequencies for each chromosome.**

| Chr. | AA | CC | GG | TT | AC | AG | AT | CG | CT | GT |
|---|---|---|---|---|---|---|---|---|---|---|
| **1** | 0.29 | 0.21 | 0.21 | 0.29 | 1.6E-05 | 6.0E-05 | 1.6E-05 | 1.6E-05 | 6.0E-05 | 1.7E-05 |
| **2** | 0.30 | 0.20 | 0.20 | 0.30 | 1.4E-05 | 4.8E-05 | 1.6E-05 | 1.5E-05 | 4.9E-05 | 1.7E-05 |
| **3** | 0.30 | 0.20 | 0.20 | 0.30 | 1.5E-05 | 5.3E-05 | 1.7E-05 | 1.5E-05 | 5.2E-05 | 1.6E-05 |
| **4** | 0.31 | 0.19 | 0.19 | 0.31 | 2.0E-05 | 6.8E-05 | 2.1E-05 | 1.7E-05 | 6.6E-05 | 2.0E-05 |
| **5** | 0.30 | 0.19 | 0.20 | 0.31 | 1.4E-05 | 4.9E-05 | 1.6E-05 | 1.4E-05 | 4.7E-05 | 1.4E-05 |
| **6** | 0.31 | 0.19 | 0.19 | 0.31 | 2.2E-05 | 8.0E-05 | 2.2E-05 | 2.1E-05 | 8.0E-05 | 2.4E-05 |
| **7** | 0.30 | 0.20 | 0.20 | 0.30 | 1.5E-05 | 5.0E-05 | 1.6E-05 | 1.4E-05 | 5.0E-05 | 1.5E-05 |
| **8** | 0.30 | 0.20 | 0.20 | 0.30 | 1.7E-05 | 5.7E-05 | 1.8E-05 | 1.9E-05 | 5.8E-05 | 1.8E-05 |
| **9** | 0.29 | 0.21 | 0.21 | 0.29 | 1.9E-05 | 6.8E-05 | 1.9E-05 | 2.1E-05 | 6.4E-05 | 2.0E-05 |
| **10** | 0.29 | 0.21 | 0.21 | 0.29 | 1.9E-05 | 6.7E-05 | 1.8E-05 | 1.9E-05 | 7.1E-05 | 2.0E-05 |
| **11** | 0.29 | 0.21 | 0.21 | 0.29 | 1.5E-05 | 5.1E-05 | 1.6E-05 | 1.3E-05 | 4.9E-05 | 1.5E-05 |
| **12** | 0.30 | 0.20 | 0.20 | 0.30 | 1.2E-05 | 4.0E-05 | 1.4E-05 | 1.2E-05 | 3.9E-05 | 1.3E-05 |
| **13** | 0.31 | 0.19 | 0.19 | 0.31 | 1.8E-05 | 5.8E-05 | 1.9E-05 | 1.5E-05 | 6.0E-05 | 1.8E-05 |
| **14** | 0.30 | 0.20 | 0.20 | 0.30 | 1.6E-05 | 5.6E-05 | 1.8E-05 | 1.5E-05 | 5.5E-05 | 1.6E-05 |
| **15** | 0.29 | 0.21 | 0.21 | 0.29 | 1.5E-05 | 5.3E-05 | 1.6E-05 | 1.6E-05 | 5.5E-05 | 1.6E-05 |
| **16** | 0.28 | 0.22 | 0.22 | 0.28 | 1.6E-05 | 5.4E-05 | 1.6E-05 | 2.0E-05 | 5.0E-05 | 1.7E-05 |
| **17** | 0.27 | 0.23 | 0.23 | 0.27 | 1.7E-05 | 6.9E-05 | 1.6E-05 | 2.0E-05 | 7.2E-05 | 1.8E-05 |
| **18** | 0.30 | 0.20 | 0.20 | 0.30 | 1.5E-05 | 4.5E-05 | 1.5E-05 | 1.3E-05 | 4.4E-05 | 1.4E-05 |
| **19** | 0.25 | 0.25 | 0.25 | 0.25 | 1.9E-05 | 5.8E-05 | 1.9E-05 | 2.1E-05 | 6.2E-05 | 1.7E-05 |
| **20** | 0.28 | 0.22 | 0.22 | 0.28 | 1.6E-05 | 6.1E-05 | 1.6E-05 | 1.8E-05 | 5.8E-05 | 1.7E-05 |
| **21** | 0.30 | 0.20 | 0.20 | 0.30 | 1.8E-05 | 5.9E-05 | 2.1E-05 | 1.8E-05 | 6.0E-05 | 2.0E-05 |
| **22** | 0.26 | 0.24 | 0.24 | 0.26 | 1.4E-05 | 5.2E-05 | 1.2E-05 | 1.8E-05 | 4.9E-05 | 1.4E-05 |
| **X** | 0.31 | 0.19 | 0.31 | 0.19 | 7.1E-06 | 1.9E-05 | 8.6E-06 | 6.3E-06 | 2.0E-05 | 7.7E-06 |

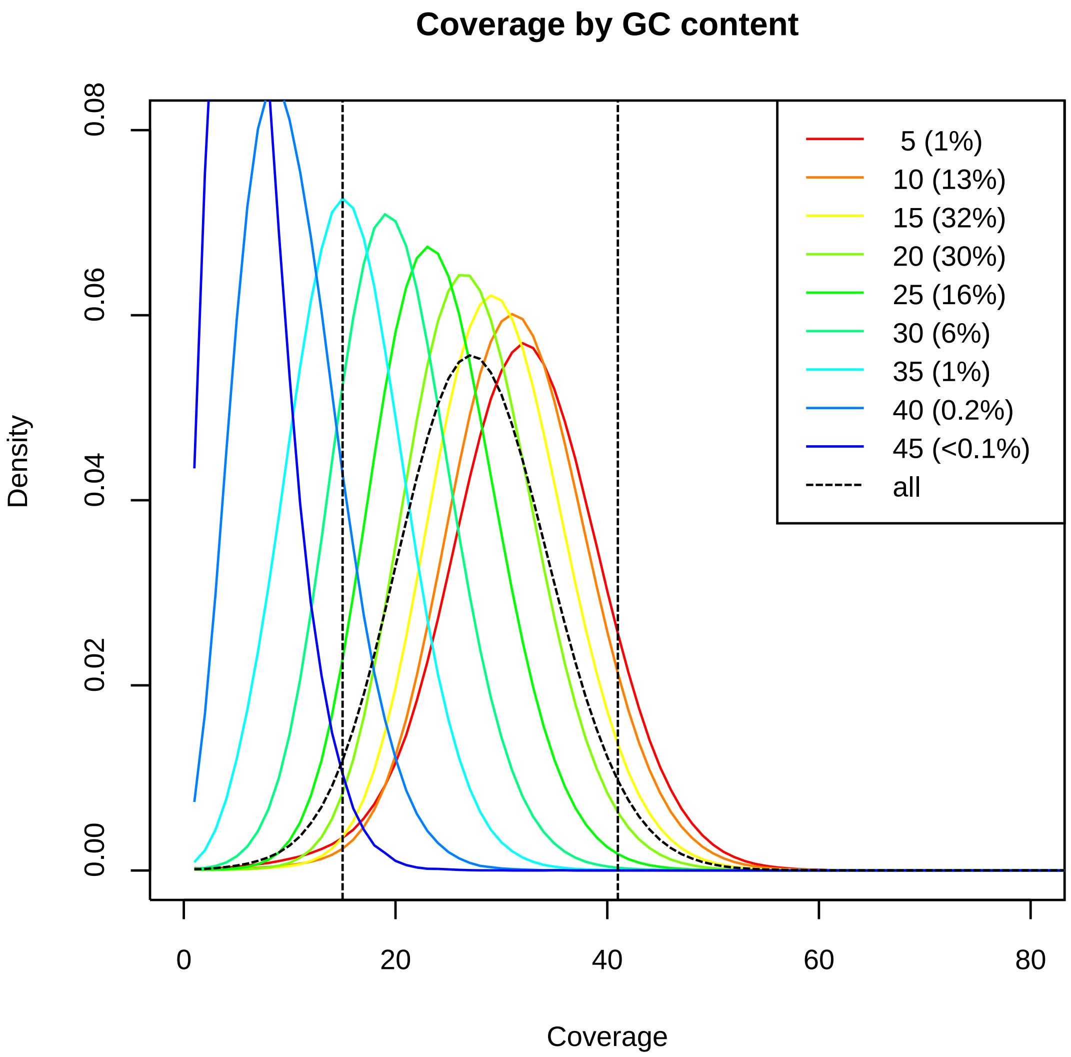


**Figure S3.2: Coverage in bins of GC content according to a sliding window of 51bp length along the genome.** Dashed lines show the average coverage over all bins and the 95% central part in this distribution. Legend: the number next to the line indicates the GC bin (5 indicates 5-9 G or C bases per 51bp window, 10 indicates 10-14, etc.); Percentages in brackets give the percentage of mappable bases that reside in each bin.

#### General Filters

As in previous analyses (4,7), we use a set of standard filters to exclude regions of the genome that are prone to alignment errors. In addition to the mapping quality cutoff of 25 and the 35mer mappability track, we exclude regions marked as simple repeats by TandemRepeatFinder (13,14), positions that are called as indels by GATK (version: 1.3-14) (8), and positions with a coverage that falls outside the central 95% of the coverage distribution per GC bin (Fig. S3.2). Finally, we require a minimum coverage of 10 for all sites, as described previously (4).

### Supplementary Information 4

#### Contamination estimates

**Summary:** We estimate the extent of modern human contamination in DNA fragments sequenced from *Chagyrskaya 8* using different approaches. Based on sites where the *Chagyrskaya 8* mitochondrial consensus sequence differs from 311 present-day humans, we estimate that 0.9% of mitochondrial DNA fragments originate from contamination. Based on fragments mapping to the Y-chromosomes, we estimate that 0.7% of fragments originate from male modern human contamination. A maximum-likelihood method to co-estimate contamination and error (1) arrives at a point-estimate for nuclear DNA contamination of 0.2%. When evaluating the state carried by DNA fragments from *Chagyrskaya 8* at a set of sites where a present-day human genome differs from two archaic hominins and four great apes, contamination is 0.7%.

##### Filters

All sequences were required to be at least 35 bases long and align with a mapping quality of at least 25. For nuclear DNA, we also required sequences to fall within the unique mappability track of 35 bp, i.e. "map35_100" as described previously (1) and as used for the general filters described in Supplementary Information 3. We only considered bases with a quality score of at least 30. To prevent deamination-derived nucleotide substitutions (2) from influencing contamination estimates, we disregarded sequences aligning in forward orientation (*i.e*. the sequenced strand is the strand in the reference sequence) at positions where one of the two possible bases is a C, and disregarded those in reverse orientation where one of the bases is a G ("strand-orientation filter").

##### Mitochondrial contamination

To assess the extent of contamination by present-day human mitochondrial DNA (mtDNA), we realigned a subset of sequences from each library (all sequences generated on the first sequencing run containing the library; see Table S2.2) to the revised Cambridge Reference Sequence (rCRS, NC_012920.1). The mtDNA genome of *Chagyrskaya 8* (see Supplementary Information 5) differs at 75 positions from the mtDNA genomes of 311 present-day humans.

The states of aligned sequences at these positions were recorded and we estimated contamination as the percentage of sequences carrying the present-day human variant with confidence given by the 95% binomial confidence interval. Contamination was estimated from all aligned sequences and from those sequences that show evidence for deamination in the original ancient DNA fragment, respectively (Table S4.1). We considered that sequences show evidence for deamination if they contained at least one C-to-T (or G-to-A) substitution compared to the reference genome within 3 bp of either sequence-end if they aligned in the forward (or reverse) orientation. Over all libraries, after removing duplicated reads, we estimate a present-day human DNA contamination of 0.9% (binomial 95% CI: 0.8-1.2%) for all sequences and 1.1% (0.6-1.8%) for putatively deaminated sequences.

**Y-chromosomal contamination**

Given that *Chagyrskaya 8* is a female (Supplementary Information 3), any contamination from male individuals will result in sequences aligning to the Y-chromosome. We estimate male contamination as Y-chromosomal sequence coverage divided by half the autosomal coverage. Table S4.1 shows that 0.7% of fragments originate from male contamination (0.5-0.8% over all libraries), a similar level to that found in *Vindija 33.19* (4).

**Autosomal contamination estimate based on the sharing of derived alleles with a present-day human genome**

We estimated the level of present-day human DNA contamination based on the proportion of sequences from *Chagyrskaya 8* that share a derived allele with a modern human genome from Africa (Mbuti, HGDP00456) (5) at positions where this modern human genome differs from the genomes of four Great Apes (4,6), a Denisovan (*Denisova 3*) (7) and a Neandertal (*Denisova 5*, or *"Altai Neandertal"*) (1).

As previously described (8), we identified positions where the African genome is different from the two archaic genomes after randomly drawing one allele whenever one of the three genomes carried a heterozygous genotype. The ancestral state was determined by requiring at least three great apes to match the archaic genomes while the remaining genome could either contain missing data or a third allele that did not match neither the African allele nor the archaic human allele.

Positions were excluded when they fell outside the published filter track for *Denisova 3* and *Denisova 5*. Over all position, we computed the proportion of *Chagyrskaya 8* sequences that carry the derived allele (*p*). This proportion *p* can be modeled as a mixture of the expected proportion of derived alleles for contaminant sequences $p_c$ and endogenous sequences $p_e$: $p = c \times p_c + (1 - c) \times p_e$.

We use the genotypes of the *Vindija 33.19* high-coverage Neandertal genome as a proxy for the genetic relationship of *Chagyrskaya 8* to the African genome ($p_e$=0.015) and assume that the present-day human DNA contamination is of non-African origin ($p_c$=0.3316 for a French, HGDP00521 (7)). With these assumptions, we can then solve the equation for *c* and estimate the contamination rate with 95% binomial confidence intervals (Table S4.1).

To test whether the contamination rate of *Vindija 33.19* and *Chagyrskaya 8* is comparable after genotype calling, we counted the number of derived alleles shared with the Mbuti genome at the positions described above. Whereas *Chagyrskaya 8* shares derived alleles less often with the Mbuti genome than *Vindija 33.19* does (Table S4.2), this difference is not significant (Fisher-exact test, p-value=0.0587). There is therefore no evidence for more human contamination in the genotypes of *Chagyrskaya 8* than in *Vindija 33.19*.

**Maximum-Likelihood estimate of autosomal contamination**

We estimated autosomal contamination using a maximum likelihood approach that co-estimates contamination and error using fixed derived sites between humans and the human-chimpanzee ancestor as previously described (1). The approach yields an estimate of 0.14% contamination for all *Chagyrskaya 8* sequence data, similar to the estimates reported for the other high-coverage archaic genomes (0.3%, 0.8% and 0.2% for *Vindija 33.19*, *Denisova 5* (Altai Neandertal) and *Denisova 3* (Denisovan), respectively) (1,4,7) . Estimates using only putatively deaminated sequences and estimates per library are reported in Table S4.1.

**Table S4.1 Present-day human contamination.** Contamination estimates were calculated using several approaches, as described in the text. The coverage per library is estimated in all uniquely mappable regions (filter map35_100). The column “Autosome” indicates the contamination estimates based on the sharing of derived alleles with a modern human (HGDP00521), while “Autosome MLE” gives the Maximum-Likelihood estimates of autosomal contamination.

| | Percentage of contamination (95% C.I.) | | | | Coverage | | |
|---|---|---|---|---|---|---|---|
| | mtDNA | Y chr. | Autosome | Autosome MLE# | X-chr | Y-chr | Autosome |
| **All sequences** | 0.9 (0.7-1.2) * | 0.7 | 0.7 (0.7-0.7) * | 0.14 (0.12-0.16) | 28.53 | 0,10 | 27,45 |
| **Sequences with terminal C-T** | 0.9 (0.5-1.6) * | 0.6 * | N/A | N/A | | | |
| **Library ID** | | | | | | | |
| **L5329** | 0.8 (0.2-2.0) | 0.7 | 0.7 (0.6-0.7) | 0.0 (0.0-2.9) | 3,59 | 0,01 | 3,46 |
| **L5727** | 0.6 (0.1-2.0) | 0.7 | 0.6 (0.5-0.7) | 0.3 (0.0-0.6) | 1,57 | 0,01 | 1,51 |
| **L5728** | 0.3 (0.0-1.8) | 0.7 | 0.5 (0.5-0.6) | 0.0 (0.0-0.2) | 1,75 | 0,01 | 1,68 |
| **L5729** | 0.0 (0.0-9.3) | 0.7 | 0.5 (0.4-0.6) | 0.4 (0.0-0.9) | 1,23 | 0,00 | 1,18 |
| **L5730** | 2.7 (0.1-14.2) | 0.7 | 0.6 (0.5-0.7) | 0.0 (0.0-0.6) | 1,07 | 0,00 | 1,02 |
| **L5794** | 0.8 (0.0-4.6) | 0.8 | 1.6 (0.9-2.4) | 11.1 (0.0-100.0) | 0,03 | 0,00 | 0,03 |
| **L5800** | 0.0 (0.0-2.3) | 0.7 | 0.6 (0.0-1.2) | 0.0 (0.0-100.0) | 0,04 | 0,00 | 0,04 |
| **L5819** | 0.8 (0.0-4.1) | 0.6 | 0.3 (0.0-1.0) | 10.1 (0.0-100.0) | 0,03 | 0,00 | 0,03 |
| **L5820** | 1.4 (0.2-5.1) | 0.5 | 0.4 (0.0-1.1) | 26.9 (0.0-100.0) | 0,03 | 0,00 | 0,03 |
| **L5821** | 1.9 (0.2-6.5) | 0.7 | 0.8 (0.1-1.4) | 0.0 (0.0-100.0) | 0,03 | 0,00 | 0,03 |
| **L5822** | 3.8 (0.8-10.8) | 0.5 | 0.3 (0.0-1.1) | 9.3 (0.0-100.0) | 0,03 | 0,00 | 0,03 |
| **L5823** | 1.5 (0.2-5.4) | 0.7 | 0.7 (0.0-1.3) | 0.0 (0.0-100.0) | 0,04 | 0,00 | 0,04 |
| **L5824** | 1.4 (0.2-5.1) | 0.7 | 0.5 (0.0-1.1) | 0.0 (0.0-100.0) | 0,03 | 0,00 | 0,03 |
| **L5830** | 0.7 (0.0-4.0) | 0.7 | 0.8 (0.7-0.9) | 0.2 (0.0-0.4) | 1,96 | 0,01 | 1,87 |
| **L5831** | 1.0 (0.2-3.0) | 0.7 | 0.6 (0.6-0.7) | 0.1 (0.0-0.2) | 3,74 | 0,01 | 3,61 |
| **L5832** | 1.2 (0.3-3.0) | 0.8 | 0.7 (0.6-0.7) | 0.0 (0.0-0.6) | 3,18 | 0,01 | 3,05 |
| **L5833** | 1.5 (0.5-3.4) | 0.7 | 0.7 (0.6-0.8) | 0.1 (0.0-0.2) | 3,04 | 0,01 | 2,93 |
| **L5848** | 0.6 (0.3-1.2) | 0.8 | 0.7 (0.7-0.8) | 0.1 (0.0-0.2) | 2,99 | 0,01 | 2,87 |
| **L5849** | 0.7 (0.3-1.3) | 0.7 | 0.8 (0.7-0.9) | 0.1 (0.0-0.2) | 2,99 | 0,01 | 2,88 |
| **L5850** | 1.4 (0.7-2.6) | 0.7 | 0.6 (0.5-0.8) | 0.0 (0.0-1.1) | 0,78 | 0,00 | 0,75 |
| **L5851** | 1.1 (0.4-2.4) | 0.7 | 0.8 (0.6-1.0) | 0.0 (0.0-2.9) | 0,39 | 0,00 | 0,37 |

* To reduce the impact of C-to-T changes, the strand orientation filter was applied.

### This method is aimed at high-coverage data, yielding large C.I.'s for individual libraries.

**Table S4.2: Number of derived and ancestral alleles in the genotypes of *Chagyrskaya 8* and *Vindija 33.19* at positions where a Mbuti genome carries a derived allele that is absent from the genomes of *Denisova 3, Denisova 5* and four great apes.** We randomly sampled one allele at heterozygous positions.

| | *Chagyrskaya 8* | *Vindija 33.19* |
|---|---|---|
| **n° derived** | 13,583 | 13,895 |
| **n° ancestral** | 899,978 | 899,688 |

### Supplementary Information 5

#### The mitochondrial genome of *Chagyrskaya 8*

**Summary:** We reconstruct the complete mitochondrial (mt) genome of *Chagyrskaya 8* and show that it falls within the variation of previously-determined Neandertal mitochondrial genomes. Using a Bayesian analysis, we estimate the tip date of the mtDNA genome to be between 58.3 and 121.8 kya.

##### The mtDNA genome of *Chagyrskaya 8*

DNA fragments from the library enriched for human mtDNA (library L5284, see Supplementary Information 2) were aligned to a modified version of the reference Neandertal mtDNA genome (NC_011137.1) (1), where the first 1,000 bases were copied to the end of the sequence to take into account that the mtDNA genome is circular (2). Sequenced fragments shorter than 35 bp and/or that mapped to the reference genome with a mapping quality lower than 25 were discarded. A total of 232,213 sequences were retained, yielding a 994-fold average coverage of the mitochondrial genome. To call a consensus, only bases with a quality of at least 30 were considered, and Ts on forward strands and As on reverse ones (relative to the reference sequence) were ignored to reduce the influence of cytosine deamination. We required each position to be covered by at least five fragments, and called a base by majority vote if at least 80% of fragments carried an identical base (2). At only two positions less than 80% of sequences carried the same base – positions 310 and 16,177 of the reference genome, which are in or adjacent to poly-C stretches. To call the base at these positions, we visually inspected a randomly-chosen subset of fragments in reverse orientation, noted the sequence of bases before and after the position in question to ensure that the fragments were accurately mapped, and determined the state carried by the majority of them. The fully-reconstructed mtDNA genome of *Chagyrskaya 8* is deposited in GenBank (accession ID MK388903).

##### Relationship to previously-determined mtDNA genomes

We computed the pairwise number of differences between the mtDNA genome sequence of *Chagyrskaya 8* and those of 27 archaic hominins, 22 ancient and present-day humans, and one chimpanzee (Table S5.1), aligned to each other using MAFFT (3). Among Neandertal-like mtDNA genomes, the *Chagyrskaya 8* mtDNA has the fewest differences to *Denisova 11* and *Okladnikov 2* (8 and 14 bases difference, respectively) from Siberia (4,5). We note that these mtDNAs have 67 and 88 uncalled bases, respectively, which may result in lower numbers of observed differences between them and the mtDNA of *Chagyrskaya 8*. The highest number of differences to a Neandertal-like mtDNA is to that of *Hohlenstein-Stadel* (99 differences), an individual from Germany previously shown to have a highly divergent mtDNA genome (6). *Chagyrskaya 8*'s mtDNA genome has 187-208 nucleotide differences to modern human mtDNA genomes; 280 differences to the mtDNA of a Middle Pleistocene hominin from Sima de los Huesos (Spain); 337-369 differences to Denisovan mtDNA genomes; and 1,432 differences to a chimpanzee mtDNA (Table S5.1). We also compared the mtDNA genome of *Chagyrskaya 8* to a partial mtDNA genome sequence reconstructed from a sediment sample collected at the site (7). We observe 3 base differences to this sequence, which may be a composite of more than one individual.

We reconstructed a maximum-likelihood tree relating the mtDNA of *Chagyrskaya 8* to other Neandertal mtDNA genomes (Fig S5.1) in MEGA 7.0.14 (8), using the mtDNA genome of a present-day Mbuti individual as outgroup. We used the Tamura-Nei model and allowed a proportion of the sites to be invariable (TrN+I) (9), a model determined using the Bayesian Information Criterion (BIC) implemented in jModelTest 2 (10). Only bases called in all mtDNA genomes were considered, and 500 bootstrap repetitions were carried out to evaluate the support for each branch.

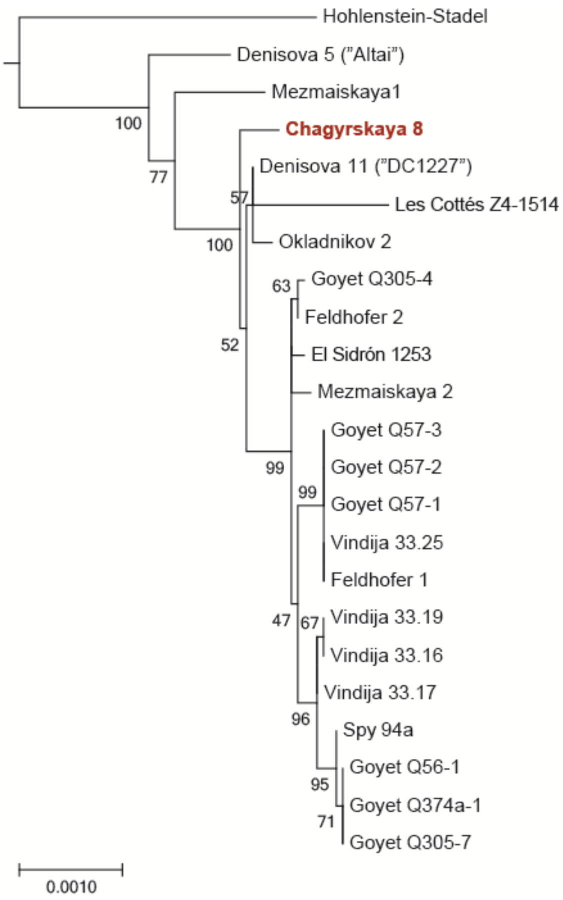


**Figure S5.1. Maximum-likelihood tree relating the mtDNA genome of *Chagyrskaya 8* to other Neandertal mtDNA genomes.** A present-day human mtDNA used as outgroup is not shown. The lengths of the branches are scaled by the number of observed substitutions per site.

#### Dating the mtDNA genome of *Chagyrskaya 8*

We estimated the tip date of the *Chagyrskaya 8* mtDNA using BEAST v.1.8.4 (43). To do so, we aligned all coding parts (positions 577-16,023 in the coordinates of the revised Cambridge Reference Sequence, as in (11)) to 22 other Neandertal mtDNAs and 22 present-day and ancient modern human mtDNA genomes using MAFFT (3). The reference Denisovan mtDNA genome (12) was used as an outgroup. For the 18 ancient individuals whose remains were directly radiocarbon-dated, we used the published dates calibrated using OxCal 4.3 (13) with IntCal13 (14) as prior on the tip dates of their mtDNA genome sequences. For the other ancient individuals, including *Chagyrskaya 8*, we set an initial value of 55kya, with the date allowed to vary between 30 and 300kya (except for two ancient individuals whose remains were found to be beyond the limit of radiocarbon dating, for whom the dates could vary between 50 and 300kya). The tip dates of present-day mtDNA genomes were allowed to vary between 0 and

100 years (Table S5.1). An initial mutation rate of $2.53 \times 10^{-8}$ (38) was used, and was allowed to vary between $1.0 \times 10^{-6}$ and $1.0 \times 10^{-10}$. The substitution model was chosen using jModelTest 2 (10) to be the Tamura-Nei model with invariable sites (TrN+I) (9). We computed the log marginal likelihood of models combining two possible tree priors (constant population size or Bayesian skyline) and two possible clock models (strict or uncorrelated relaxed lognormal) in 30,000,000 iterations with additional 1,000,000 iterations (stepping-stone sampling, frequency every 1,000 steps), and used the scale defined by Kass and Raftery (15) to evaluate the support for each one. The Bayesian skyline prior was better supported than the constant population size ($log_{10}BF=3$ and $log_{10}BF=0.8$ in combination with the strict or the relaxed clock, respectively). As there was no notable difference between the clock models ($log_{10}BF=0.3$), the simpler strict clock model was retained. With these parameters, we combined four runs of 75,000,000 iterations each (discarding the first 10% of each as burn-in). The resulting estimated tip date of the *Chagyrskaya 8* mtDNA is 89.4 kya (95% highest posterior density: 58.3-121.8kya).

**Table S5.1. Mitochondrial genomes used in the analyses.** The number of observed differences between each mtDNA genome and the mtDNA of *Chagyrskaya 8* is noted. Accession IDs from the NCBI database, the priors used for the tip dating analysis and references are given.

| mtDNA type | Individual | Number of differences | Prior on date | ID | References |
|---|---|---|---|---|---|
| Neandertal | Denisova 5 (“Altai”) (Russia) | 38 | 55,000 (30,000-300,000) | KC879692 | ref.(16) |
| | Denisova 11 (“DC1227”) (Russia) | 8 | 55,000 (50,000-300,000) | KU131206 | ref. (4) |
| | El Sidrón 1253 (Spain) | 22 | 55,000 (30,000-300,000) | FM865409 | ref.(17) |
| | Feldhofer 1 (Germany) | 24 | 43,707 (42,670-44,744) | FM865407 | refs.(17,18) |
| | Feldhofer 2 (Germany) | 20 | 43,268 (42,193-44,342) | FM865408 | refs.(17,18) |
| | Goyet Q56-1 (Belgium) | 30 | 42,515 (42,063-42,967) | KX198082 | ref.(19) |
| | Goyet Q57-1 (Belgium) | 20 | 44,696 (43,834-45,558) | KX198082 | ref.(19) |
| | Goyet Q57-2 (Belgium) | 25 | 41,185 (40,595-41,775) | KX198088 | ref.(19) |
| | Goyet Q57-3 (Belgium) | 25 | 42,407 (41,946-42,867) | KX198083 | ref.(19) |
| | Goyet Q305-4 (Belgium) | 21 | 44,236 (43,386-45,085) | KX198087 | ref.(19) |
| | Goyet Q305-7 (Belgium) | 30 | 55,000 (30,000-300,000) | KX198086 | ref.(19) |
| | Goyet Q374a-1 (Belgium) | 30 | 55,000 (30,000-300,000) | KX198085 | ref.(19) |
| | Hohlenstein-Stadel (Germany) | 99 | 55,000 (30,000-300,000) | KY751400 | ref.(6) |
| | Les Cottés Z4-1514 (France) | 34 | 43,230 (42,720-43,740) | MG025536 | ref.(20) |
| | Mezmaiskaya 1 (Russia) | 36 | 55,000 (30,000-300,000) | FM865411 | ref.(17) |
| | Mezmaiskaya 2 (Russia) | 23 | 43,834 (42,038-45,630) | MG025537 | ref. (20,21) |
| | Okladnikov 2 (Russia) | 14 | 55,000 (30,000-300,000) | KF982693 | ref.(5) |
| | Spy 94a (Belgium) | 29 | 40,463 (39,840-41,085) | MG025538 | refs.(20,22) |
| | Vindija 33.16 (Croatia) | 26 | 43,707 (39,234-48,179) | NC_011137 | refs. (1,23) |
| | Vindija 33.17 (Croatia) | 24 | 55,000 (30,000-300,000) | KJ533544 | ref.(24) |
| | Vindija 33.19 (Croatia) | 26 | 55,000 (50,000-300,000) | KJ533545 | refs.(24,25) |
| | Vindija 33.25 (Croatia) | 24 | 55,000 (30,000-300,000) | FM865410 | ref.(17) |
| Modern human | Australian | 203 | 0 (0-100) | AF346964 | ref.(26) |
| | Chinese | 193 | 0 (0-100) | AF346973 | ref.(26) |
| | Filipino | 196 | 0 (0-100) | AY289070 | ref.(27) |
| | Finnish | 196 | 0 (0-100) | AY195773 | ref.(28) |
| | Indian | 190 | 0 (0-100) | AF382013 | ref.(29) |
| | Italian | 201 | 0 (0-100) | AY882393 | ref.(30) |
| | Japanese | 190 | 0 (0-100) | AF346990 | ref.(26) |
| | Mandenka | 202 | 0 (0-100) | AF346995 | ref.(26) |
| | Mbuti | 199 | 0 (0-100) | AF346998 | ref.(26) |
| | Native American | 191 | 0 (0-100) | AY195748 | ref.(28) |
| | Pakistani | 203 | 0 (0-100) | AY882380 | ref.(30) |
| | Papuan (Coast) | 203 | 0 (0-100) | AY289082 | ref.(27) |
| | Papuan (Highlands) | 194 | 0 (0-100) | AY289090 | ref.(27) |
| | San | 206 | 0 (0-100) | AF347008 | ref.(26) |
| | Spanish | 208 | 0 (0-100) | AY882392 | ref.(30) |
| | Boshan 11 (China) | 187 | 8,234 (8,152-8,316) | KC521454 | ref.(11) |
| | Iceman (Austria/Italy) | 200 | 5,300 (5,275-5,325) | EU810403 | refs.(31,32) |
| | Kostenki 14 (Russia) | 194 | 37,473 (36,262-38,684) | FN600416 | refs.(32,33) |
| | Loschbour (Luxembourg) | 200 | 8,054 (7,948-8,160) | KC521455 | refs.(11,34) |
| | Saqqaq Eskimo (Greenland) | 195 | 4,504 (4,423-4,585) | EU725621 | refs.(35,36) |
| | Tianyuan (China) | 190 | 39,008 (37,761-40,254) | KC417443 | refs.(37) |
| | Ust‘-Ishim (Russia) | 191 | 45,045 (43,212-46,878) | - | ref.(38) |
| Denisovan | Denisova 2 (Russia) | 337 | N/A | KX663333 | ref.(39) |
| | Denisova 3 (Russia) | 369 | 55,000 (30,000-300,000) | NC_013993 | ref.(12) |
| | Denisova 4 (Russia) | 369 | N/A | FR695060 | ref.(40) |
| | Denisova 8 (Russia) | 349 | N/A | KT780370 | ref.(41) |
| SH | Femur VIII (Spain) | 280 | N/A | NC_023100 | ref.(2) |
| Chimpanzee | - | 1,432 | N/A | NC_001643 | ref.(42) |

SH – Sima de los Huesos; N/A – this genome was not included in the tip dating analysis.

### Supplementary Information 6

#### Demography and molecular dating

**Summary:** We estimated the demographic history of *Chagyrskaya 8* using the Pairwise Sequentially Markovian Coalescent (PSMC) (1), and find that the population size gradually declined over time, similarly to what is observed for the two previously sequenced high-coverage Neandertal genomes. Leveraging the common/overlapping time intervals of these inferred demographic histories, we estimate that *Chagyrskaya 8* lived approximatively 30,000 years before *Vindija 33.19* (2), and 50,000 years after *Denisova 5* (3). These times are consistent with age estimates based on the number of derived substitutions in the *Chagyrskaya 8* genome, suggesting that *Chagyrskaya 8* lived ~80,000 years ago.

##### Population demography estimates

We applied PSMC to the genotype data of *Chagyrskaya 8*, *Vindija 33.19* and *Denisova 5,* filtered as described in Supplementary Information 3. As PSMC parameter estimates are affected by low coverage data and filtering of data (2,4), we calibrate our estimates using simulations, similarly to Prüfer et al.,2017 (2). Specifically, we first applied PSMC on filtered data to obtain a demography *D*, consisting of the population size history $N_t$, relative to the population size at time 0, and two global scaling parameters $\theta$ and $\rho$, that reflect the effective population size and the mutation rate at time 0, i.e. when the individual lived, and the recombination rate, respectively. Our aim is to find the combination of parameters $\theta'$ and $\rho'$ that would result in the demography *D* when estimated from filtered data. We thus generate multiple whole-genome coalescent simulations by fixing $N_t$ but varying the values of $\theta$ and $\rho$ to $\theta'$ and $\rho'$. The simulated genomes are then filtered analogously to the original data, and PSMC is applied to infer new demographies *D'*. The corrected combination of $\theta'$ and $\rho'$ parameters is thus inferred by selecting that which better explains the observed data, i.e. that resulting in the demographic history *D'* that provides the highest likelihood when applied to the original data. This likelihood can be computed directly via PSMC, using the –i flag. This last step differs from the approach used in Prüfer et al.,2017 (2), where the best fit was assessed with a Least Squares quantification and only $\theta'$ was estimated. We performed 400 simulations in a 20x20 grid of *θ'* and *ρ'* values obtained by rescaling the original *θ* and *ρ* parameters by factors between 1 to 1.6, and 1 to 3, respectively. These ranges were shown in preliminary explorations of the data to best fit the data. Maximum likelihood estimates were then obtained after interpolating the grids of values with the R package *mgcv*. Likelihood surfaces are shown in Fig.S6.1. The estimated correction factors are broadly similar for the three Neandertal genomes (Fig.S6.1, black markers). For *Vindija 33.19* and *Denisova 5*, the *θ* correction factors are between 130% and 140%, as estimated in Prüfer et al.2017 (2). The demographic histories obtained after the correction are plotted in Fig.S6.2.

Genomes following these demographies can be generated via coalescent simulations using the code reported in *ms* (5) format in Fig.S6.3.

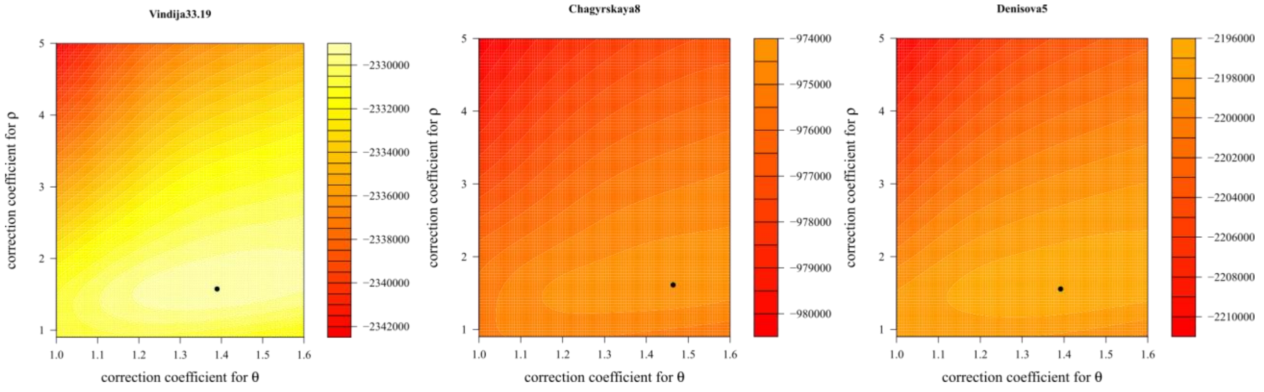


**Fig.S6.1. Correction estimates for the demographic histories of *Vindija33.19*, *Chagyrskaya 8* and *Denisova 5*.** Correction factors for the recombination ratio *θ* and the *ρ* parameters are shown on the y-axis and x-axis, respectively. Lighter colours indicate higher likelihood values calculated by PSMC, when applied to genomes generated via simulation using values of *θ* and *ρ* corrected by the values reported on the x and y axis, respectively. Black markers indicate the combinations of parameters for which PSMC provides the highest likelihood.

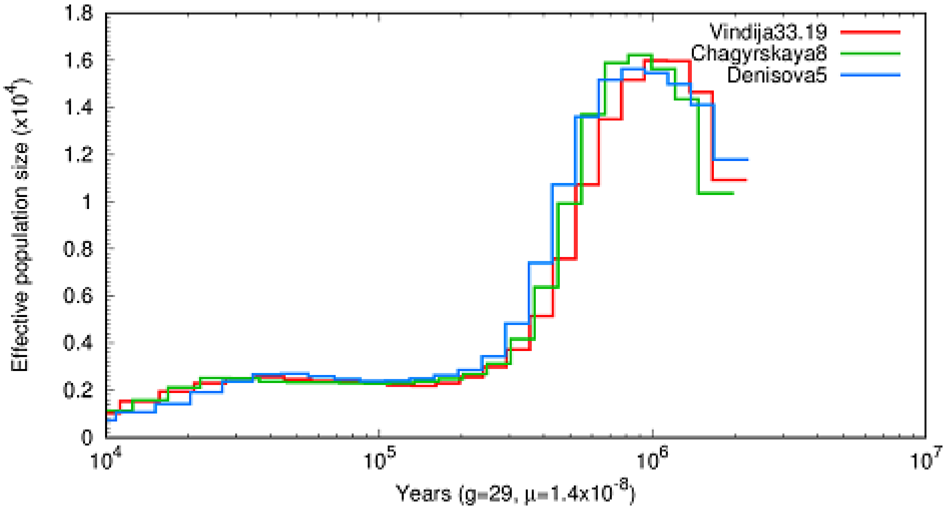


**Fig.S6.2. Demographic histories for three Neandertal genomes estimated by PSMC.** Time is reported on the x-axis for each Neandertal from up to $10^4$ years before the time of death, assuming a mutation rate per year of $0.5 \times 10^{-9}$ years$^{-1}$ bp$^{-1}$, corresponding to $1.45 \times 10^{-8}$ gen$^{-1}$ bp$^{-1}$ for a generation time of 29 years. Note that the time of death differs for the three individuals. Thus, the x-axis does not reflect an absolute time in the past, and the demographic histories of the more recent Neandertals are expected to be shifted rightwards compared to those of the more ancient individuals.

*Vindija 33.19:*

ms 2 1 -t 13041.14772872183 -r 3048.50564922064 249230605 -en 0.0436 1 0.4496 -en 0.0724 1 1.1386 -en 0.1073 1 1.6976 -en 0.1494 1 2.1414 -en 0.2004 1 2.5273 -en 0.2620 1 2.7677 -en 0.3365 1 2.8236 -en 0.4266 1 2.7121 -en 0.5355 1 2.6260 -en 0.6671 1 2.6174 -en 0.8264 1 2.5544 -en 1.0189 1 2.4373 -en 1.2516 1 2.4051 -en 1.5331 1 2.5286 -en 1.8733 1 2.8193 -en 2.2848 1 3.2856 -en 2.7822 1 4.1041 -en 3.3837 1 5.6754 -en 4.1109 1 8.3609 -en 4.9902 1 11.8388 -en 6.0533 1 14.8836 -en 7.3388 1 16.7310 -en 8.8930 1 17.6350 -en 10.7722 1 17.6137 -en 13.0443 1 16.1529 -en 15.7915 1 12.0577 -en 23.1293 1 44.8768

*Chagyrskaya 8:*

ms 2 1 -t 2748.38749554598 -r 653.111605863336 249230605 -en 0.1728 1 2.0395 -en 0.2880 1 4.2353 -en 0.4282 1 6.3325 -en 0.5986 1 8.8150 -en 0.8058 1 11.6664 -en 1.0577 1 13.8983 -en 1.3640 1 13.8425 -en 1.7363 1 13.0453 -en 2.1891 1 12.8561 -en 2.7396 1 12.8571 -en 3.4090 1 12.6969 -en 4.2228 1 12.5652 -en 5.2123 1 12.6974 -en 6.4153 1 13.1013 -en 7.8781 1 13.7268 -en 9.6566 1 14.8088 -en 11.8190 1 17.2506 -en 14.4482 1 23.0490 -en 17.6449 1 35.2529 -en 21.5317 1 54.9353 -en 26.2574 1 75.7921 -en 32.0032 1 87.9151 -en 38.9893 1 89.7266 -en 47.4833 1 86.4670 -en 57.8109 1 79.3633 -en 70.3677 1 57.3675 -en 104.1979 1 206.3010

*Denisova 5:*

scrm 2 1 -t 5484.46414949352 -r 1273.698712986202 249230605 -en 0.0997 1 1.3763 -en 0.1659 1 1.9359 -en 0.2462 1 2.8299 -en 0.3434 1 3.7586 -en 0.4613 1 5.0165 -en 0.6041 1 6.2030 -en 0.7773 1 6.9742 -en 0.9871 1 7.0692 -en 1.2415 1 6.7771 -en 1.5497 1 6.4688 -en 1.9233 1 6.2562 -en 2.3760 1 6.2782 -en 2.9247 1 6.5188 -en 3.5898 1 6.8857 -en 4.3958 1 7.4881 -en 5.3727 1 8.9953 -en 6.5566 1 12.6641 -en 7.9916 1 19.4238 -en 9.7306 1 28.1659 -en 11.8384 1 35.6656 -en 14.3929 1 39.7922 -en 17.4889 1 40.9909 -en 21.2412 1 40.5491 -en 25.7889 1 39.3030 -en 31.3005 1 37.0012 -en 37.9805 1 30.9487 -en 55.8887 1 101.6739

**Fig.S6.3. Code to simulate the demographic history of the three Neandertal genomes.** The code is reported in *ms* format for a genomic tract of length similar to that of chromosome 1.

**Age estimates using demographic histories**

We used the inferred demographic histories to estimate the differences in age among the three Neandertal genomes by a strategy similar to that used in Fu et al., 2014 (6) to estimate the age of an ancient modern human. This method has the advantage of not relying exclusively on private mutations, which can be enriched in errors (6). Assume that two genomes, $G$ and $G_{old}$, are sampled from a population with a demographic history $D$ at different times in the past, $\tau$ and $\tau_{old}$, respectively. Using PSMC, we can estimate the demographic history from these genomes: in this case, while for the younger genome $G$ we would estimate a demography $D$ up to time $\tau$, i.e. $D(\tau)$, for the more ancient genome we would estimate a demography $D_{old}(\tau_{old})$. Note that since $G$ and $G_{old}$ share the same demographic history up to time $\tau_{old}$, $D_{old}(\tau_{old})$ is expected to be approximatively equal to $D\ (\tau-\ \tau_{diff})$, where $\tau_{diff}=\tau-\tau_{old}$ is the difference in age between $G$ and $G_{old}$. We take advantage of this to estimate $\tau_{diff}$, by calculating the value of $\tau_{diff}$ that maximizes the likelihood of $D(\tau-\tau_{diff})$ given the genome of the more ancient individual. This can be computed by truncating the demography $D(\tau)$ estimated from the younger genome at a time $\tau-\ \tau_{diff}$ in the past, and using PSMC to compute the likelihood of this demography on the more ancient genome $G_{old}$.

Note that this approach cannot be performed directly on the corrected demographies described in the previous section, since filtering might affect the estimates for different genomes differently. Thus, we performed whole genome simulations following the corrected demographic histories of genome $G$, and then filtered them as the target observed genome $G_{old}$. We thus applied PSMC on this simulated genome to obtain a demographic history $D(\tau)$ matching the same filter as the observed data $G_{old}$. This PSMC demographic history is then used to estimate $\tau_{diff}$ as described above and in Fu et al.,2014 (6). We provide a step by step protocol of this procedure in Fig.S6.4

To validate this approach, we perform simulations under the demographic history estimated from *Vindija 33.19* using filters learned for the *Denisova 5* genome. We simulated 49 pairs of genomes differing in age between 0 and 150 ky, assuming a mutation rate of $0.5\text{x}10^{-9}$ $\text{years}^{-1}$ $\text{bp}^{-1}$. For all age differences, the correct demographic history provided the highest likelihood (Fig. S6.5).

We then applied this method to real data. Compared to the *Denisova 5* genome, we estimated that *Chagyrskaya 8* and *Vindija33.19* are ~50,000 and years ~80,000 younger, respectively (Fig.S6.6, left and middle panels). When using the *Chagyrskaya 8* genome, the highest likelihood for $t_{diff}$ when using the *Denisova 5* demography is equal to 0, confirming that

*Chagyrskaya 8* is younger than *Denisova 5*; $t_{diff}$ is instead estimated to be ~8,000 when using the demographic history of *Vindija33.19.* Note, however, that the posterior distribution for this estimate is bimodal, suggesting that the relatively smaller age difference between *Chagyrskaya 8* and *Vindija 33.19* could affect our estimates. Note how this ordering in the age of the samples is reflected visually in the leftward shift of the PSMC demographic history of *Denisova 5* relative to *Chagyrskaya 8*, and of *Chagyrskaya 8* relative to *Vindija 33.19* (Fig.S6.2).

Step 1) Truncation of the PSMC demographic history $D(\tau)$ estimated from genome *G* at time $\tau$-$\tau_{diff}$ to obtain a demographic history $D(\tau-\tau_{diff})$, using the script *psmc_trunc.pl* included in the PSMC package.

Step 2) Whole genome coalescent simulations following demography $D(\tau-\tau_{diff})$ (software *ms*). The resulting genome is then filtered as genome $G_{old}$.

Step 3) PSMC is performed on the simulated genome to obtain a newly estimated demography $D'(\tau-\tau_{diff})$ on the filtered data. The PSMC options used for the estimates were:

*psmc -N25 -t15 -r5 -p "4+25*2+4+6"*

Step 4) The likelihood of $D'(\tau-\tau_{diff})$ given genome $G_{old}$ is computed with PSMC, using the options:

*psmc -N1 –I* $G_{old}$

Step 5) The value of $\tau_{diff}$ maximizing the likelihood of $D'(\tau-\tau_{diff})$ corresponds to the estimated age difference between genome *G* and genome $G_{old}$. Uncertainty was quantified by repeating the estimation for 2000 simulated genomes and retaining the 2.5% top simulations with highest likelihood.

**Fig.S6.4. Procedure to estimate the difference in age between two genomes G and $G_{old}$.**

#### Age estimates using branch shortening

We calculated the proportion of missing mutations in Neandertals compared to modern human genomes as for the two previously published Neandertal high-coverage genomes (2,3). *Chagyrskaya 8* accumulated 0.62±0.09% less mutations than an African genome from the SGDP dataset (7) (S-Mbuti-2) from the separation from chimpanzees, compared to 0.40±0.09% for *Vindija 33.19*, 0.94±0.1% for *Denisova 5*, and 0.55±0.9% for *Denisova 3*. By assuming a mutation rate of $0.5 \times 10^{-9}$ years$^{-1}$ bp$^{-1}$ we estimate that *Chagyrskaya 8* lived about 80 ky ago, *Vindija 33.19* about 55 ky ago, and *Denisova 5* 115 ky ago. These ages are broadly consistent with age differences estimated using PSMC. However, *Denisova 5* is estimated to be approximately 60ky older than *Vindija 33.19* based on branch shortening, whereas the PSMC point estimate is at 80ky. However, the confidence interval for the latter estimate is 50-110ky.

To assess the extent to which this discrepancy could be due to reference bias (derived mutations specific to Neandertals might be lost during mapping), we mapped the *Chagyrskaya 8* reads to an ancestralized reference, following the scheme described in Peyrégne et al.,2019 (8). We then generated genotypes data in the same way as for the hg19 mapped data. No significant difference in the age estimates could be detected (Fig S6.7).

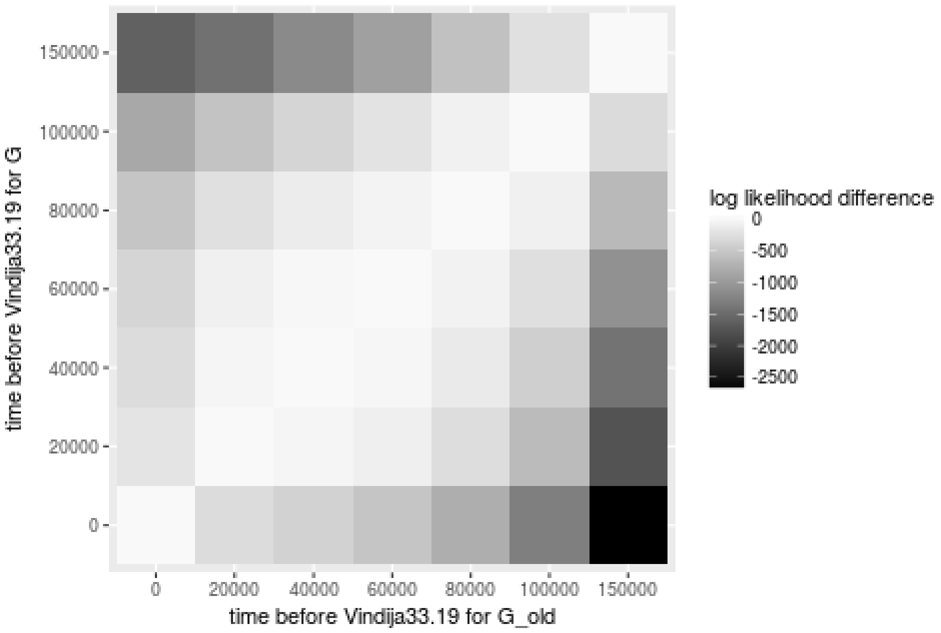


**Fig.S6.5. Validation of the PSMC based dating method.** Two independent genomes *G* and $G_{old}$ were simulated using *ms* following the estimated demographic history of *Vindija 33.19* and sampled at different times before *Vindija 33.19*, shown on the y-axis and x-axis, respectively.

Both genomes were filtered as *Denisova 5*. We then followed the procedure described in Fig.S6.4 and re-estimated the demographic history using PSMC for the genome *G*. The likelihood of this model given genome $G_{old}$ is reported with lighter shades of gray the higher the likelihood, after subtracting the maximum likelihood value for each column. The correct values of $\tau_{diff}$ are found along the diagonal, and always show the highest likelihood among those tested. Note that in this plot $G_{old}$ can be younger than *G* (upper triangle).

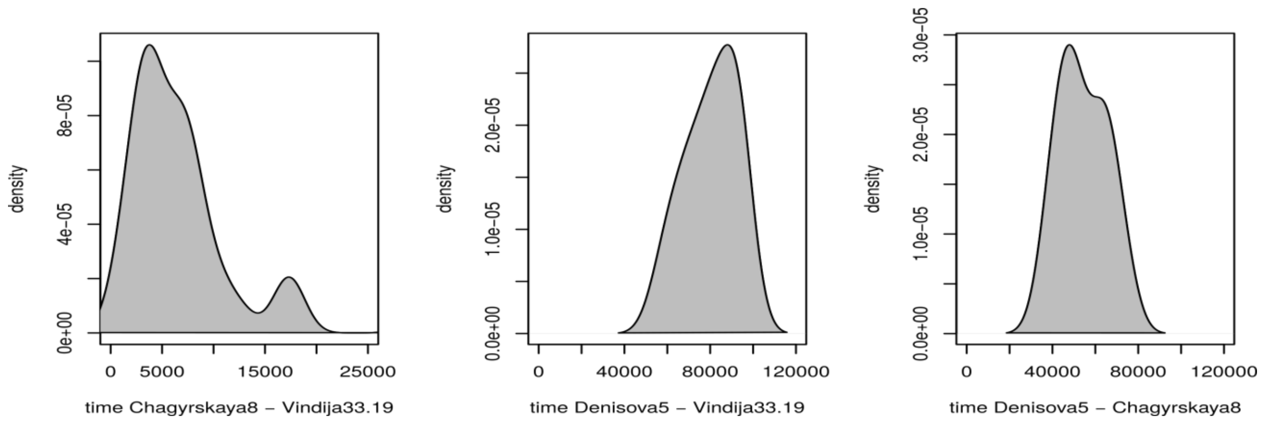


**Fig.S6.6. Dating of Neandertal genomes using the inferred demographic histories.** Posterior probabilities for the estimated age difference between two Neandertal genomes $t_{diff}$ (x-axis) was estimated by calculating the likelihood of the demography $D'(t\text{-}t_{diff})$ estimated for the younger Neandertal given the genome of the more ancient Neandertal (y-axis). The more recent and more ancient Neandertals are in order *Chagyrskaya 8* and *Vindija 33.19* (left), *Denisova 5 and Vindija 33.19* (center), *Denisova 5* and *Chagyrskaya 8* (right).

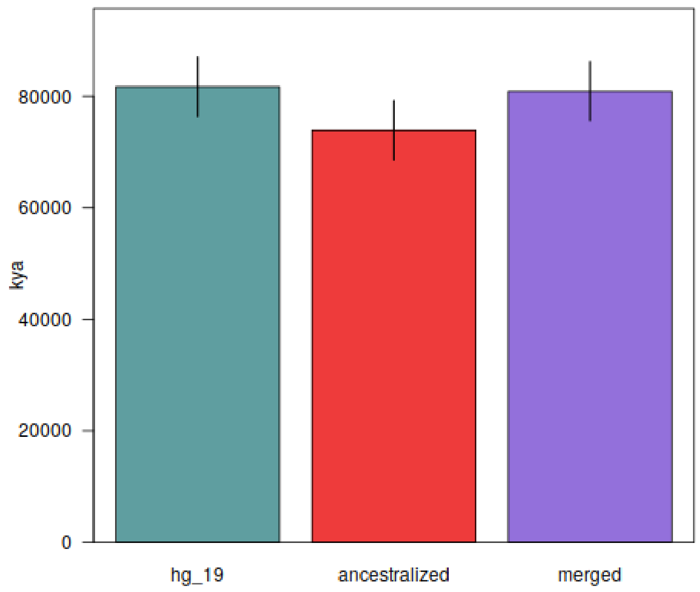


**Fig.S6.7. Dating of *Chagyrskaya 8* using branch shortening.** We calculated missing mutations in respect to a present-day African human genome (S-Mbuti 2 from the Simons Genome Diversity Project) (7) in 5Mb blocks, used to estimate the 95% confidence intervals via block jacknife (black lines). We calculated missing mutations for genotypes obtained by mapping DNA fragments to either the human reference hg19 (left, green), an ancestralized reference including derived positions in at least one of three previously sequenced archaic genomes (*Vindija 33.19, Denisova 5 and Denisova 3*)(center, red), or both reference genomes using the scheme adopted in Peyrégne et al.,2019 (8) (right, purple).

### Supplementary Information 7

#### Relationships to other archaic hominins and to modern humans

**Summary:** We estimate that *Chagyrskaya 8* was more closely related to *Vindija 33.19* and other Neandertals living in Europe ~45-55 kya than to *Denisova 5* (the "*Altai Neandertal*") and that the most recent common ancestor of *Chagyrskaya 8* and *Vindija 33.19* lived ~100 kya. There is no significant difference in the numbers of derived alleles that *Chagyrskaya 8* or *Vindija 33.19* share with modern humans. However, compared to *Chagyrskaya 8*, *Vindija 33.19* shares more derived alleles that segregate at low frequencies in present-day human populations with modern humans, suggesting that *Vindija 33.19* was more closely related to Neandertal populations that mixed with modern human populations.

*Chagyrskaya 8* is the most closely related among currently available Neandertals to the mother of *Denisova 11*, the offspring of a Neandertal mother and a Denisovan father, discovered in Denisova Cave. We estimate that the last common ancestor of the Neandertal mother of *Denisova 11* and *Chagyrskaya 8* lived ~10 kya before *Chagyrskaya 8*.

##### *D*-statistics

For the high-coverage genomes *Chagyrskaya 8*, *Vindija 33.19* (1), *Denisova 5* (2) and *Denisova 3* (3), we used snpAD genotype calls for sites that pass recommended filters (http://cdna.eva.mpg.de/neandertal/Vindija/FilterBed) (1). For the low-coverage Neandertal genomes (2,4), as well as for *Denisova 11* (5), at each position we randomly sampled a base from fragments that had a mapping quality ≥ 25, base quality ≥ 30 and presented evidence for deamination. For the low coverage Neandertal *Mezmaiskaya 1*, which has a coverage of ~2-fold, we also compare results obtained with random reads with genotype calls obtained using snpAD. Modern human genotypes were extracted from 271 individuals of the Simons Genome Diversity Panel (SGDP, filtered for genotype quality ≥ 1) (6) and from 2,504 individuals of the 1000 Genomes Project Phase III release (7). The genotypes of the ancient modern humans *Stuttgart* (~7kya, Germany) (8), *Loschbour* (~8kya, Luxembourg) (8) and *Ust'-Ishim* (~45kya, Siberia) (9), as well as randomly sampled deaminated fragments from the low-coverage *Oase 1* genome (~40kya, Romania) (10) were also used to estimate the genetic contribution of different Neandertals to modern humans.

By default, we used the chimpanzee genome (*panTro4*) as outgroup. We additionally used the rhesus (*rheMac3*) reference genome (11) as outgroup to estimate F(A|B) statistics.

*D*-statistics (12,13) of the form *D*(A,B,C,O) were calculated across all bi-allelic transversions on the autosomes as

$$D = \frac{\sum\big(p(BABA) - p(ABBA)\big)}{\sum\big(p(BABA) + p(ABBA)\big)},$$

with

$$p(ABBA) = (1 - f_A)f_B f_C(1 - f_O) + f_A(1 - f_B)(1 - f_C)f_{O,}$$
$$p(BABA) = f_A(1 - f_B)f_C(1 - f_O) + (1 - f_A)f_B(1 - f_C)f_O$$

where $f_X$ denotes the allele frequency in genome/group $X$ at one position. Heterozygous calls in individual genomes were assigned a frequency of 0.5. Standard errors (SE) were calculated using a weighted block jackknife procedure (14) over autosomes divided into 5 Mb blocks. *Z*-scores are defined as $Z = D/SE$.

**Relationship of *Chagyrskaya 8* to other archaic hominins**

We first computed *D*(*Chagyrskaya 8*, *Denisova 5/Vindija 33.19*, archaic hominin, chimpanzee) to test which archaic hominin genomes share more derived alleles with *Chagyrskaya 8* than with the other two high-coverage Neandertals. We find that all Neandertals tested and the Denisovan-Neandertal offspring *Denisova 11* share significantly more derived alleles with *Chagyrskaya 8* than with *Denisova 5* (the "*Altai Neandertal")* (25% ≤ *D* ≤ 36%, *Z* > 22) (Fig.S7.1a). The Denisovan *Denisova 3*, however, shares more derived alleles with *Denisova 5* than with *Chagyrskaya 8* (*D* = -4.2%, *Z* = -3.0), consistent with gene flow from Neandertals related to *Denisova 5* into the ancestors of *Denisova 3* (2).

When comparing *Chagyrskaya 8* and *Vindija 33.19*, we find that the Neandertals *Mezmaiskaya 2*, *Goyet*, *Les Cottés*, *Spy* and *Vindija 87* share significantly more derived alleles with *Vindija 33.19* than with *Chagyrskaya 8* (-74% ≤ *D* ≤ -22%, *Z* < -19) (Fig.S7.1b). As expected, this signal is strongest for *Vindija 87*, a bone that was inferred to belong to the same individual as *Vindija 33.19* (4). *Denisova 5* and *Mezmaiskaya 1* share a similar number of derived alleles with *Vindija 33.19* as they do with *Chagyrskaya 8* (*Z* = -1.4 for *Denisova 5*, *Z* = -2.2 for *Mezmaiskaya 1* genotypes and *Z* = -1.8 for randomly drawn deaminated reads for *Mezmaiskaya 1*) (Fig.S7.1b). These and earlier results suggest that *Denisova 5* diverged first from the ancestors of *Chagyrskaya 8*, *Mezmaiskaya 1* and later Neandertals. However, *Mezmaiskaya 1* shares slightly more derived alleles with *Vindija 33.19* than it does with *Chagyrskaya 8.* This suggests that *Chagyrskaya 8* might have diverged earlier from the remaining Neandertals. However, we emphasize that this signal is observed only when genotypes calls are used for *Mezmaiskaya 1*, and it is not observed when a closer outgroup (*i.e*. *Denisova 5*rather than a chimpanzee) is used (Fig S7.2).

The Denisovan genome (*Denisova 3*) shows no difference in the sharing of derived alleles with *Vindija 33.19* and with *Chagyrskaya 8* ($Z$ = 0.6). However, the Denisovan-Neandertal offspring *Denisova 11* shares more alleles with *Chagyrskaya 8* than with *Vindija 33.19* ($D$ = 5.0%, $Z$ = 3.6), suggesting that *Chagyrskaya 8* is more closely related to the Neandertal mother of *Denisova 11* than *Vindija 33.19* is (Fig S7.1b).

**Figure S7.1: Relative sharing of derived alleles between *Chagyrskaya 8* and other archaic individuals.** Positive values indicate a higher allele sharing of X with *Chagyrskaya 8* than with *Denisova 5* (a) or *Vindija 33.19* (b). NEA: Neandertals, ND: Neandertal-Denisovan offspring, DEN: Denisovans. Error bars indicate 1 SE. For all low coverage genomes, random reads were used. For *Mezmaiskaya 1* genotype data are also shown, indicated by GT.

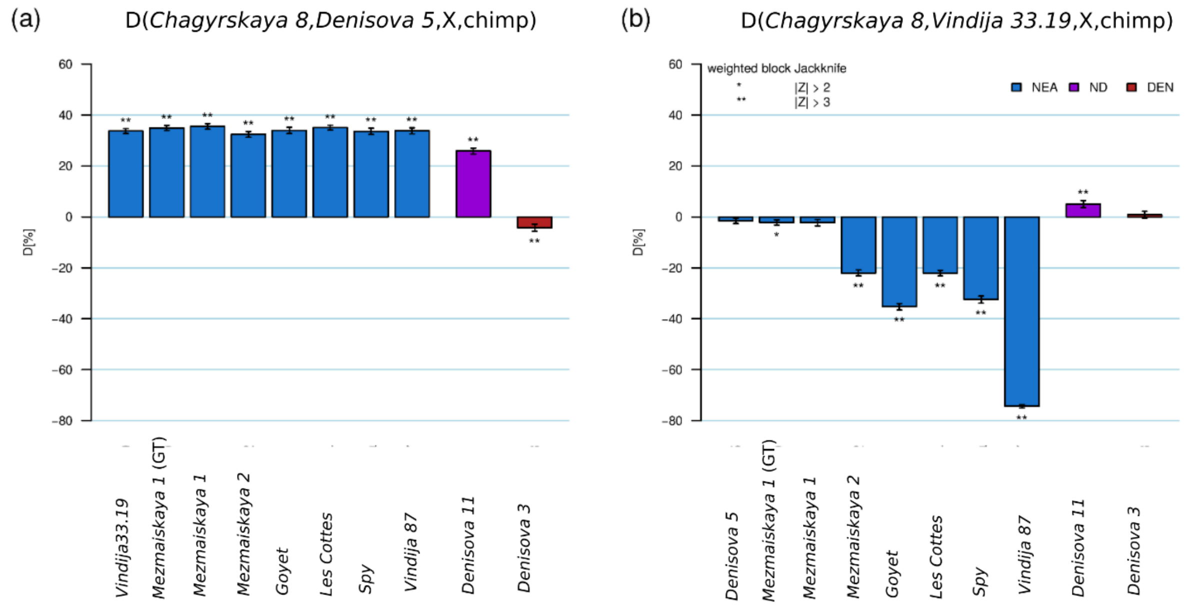

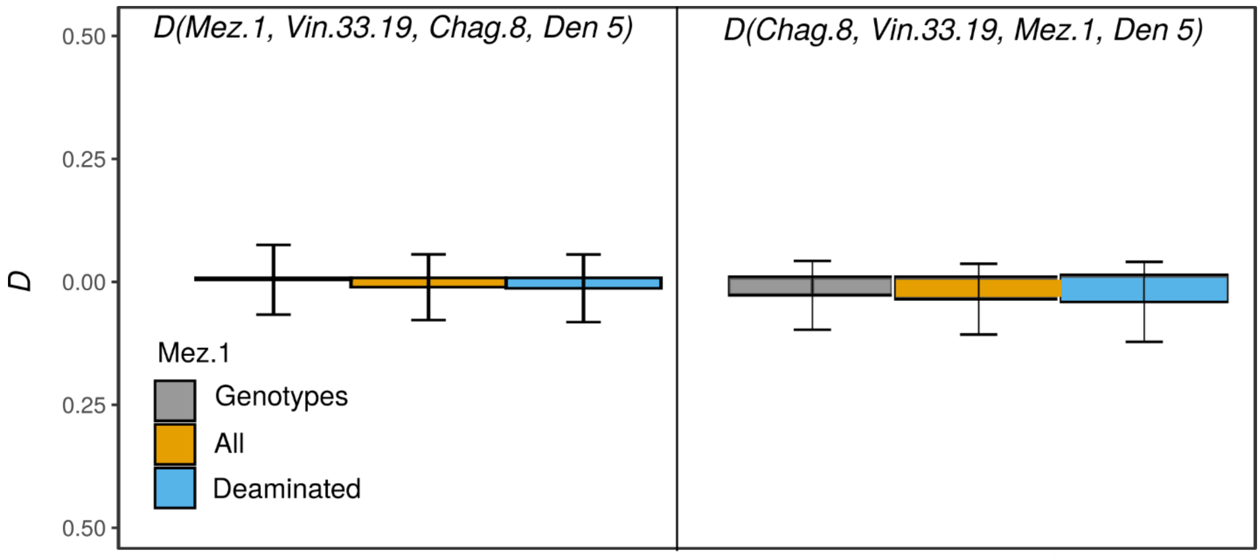


**Figure S7.2: Relative sharing of derived alleles between *Chagyrskaya 8* and other archaic individuals using *Denisova 5* as outgroup.** *D* statistic computed using snpAD genotype calls, random reads and random deaminated reads only are shown in gray, brown and blue, respectively.

**Population split time estimates**

To estimate population split times between *Chagyrskaya 8* and other Neandertals, we computed the F(A|B) statistic. The F(A|B) statistic is the proportion of alleles from population A carrying the derived state at sites where a genome from population B is heterozygous. We compute it as described in ref. (2). To translate the F(A|B) statistic to population split times, we use the demography histories of population B estimated with *PSMC* in Supplementary Information 6.

To correct for reference bias which may be introduced during mapping, data from low coverage Neandertal genomes were mapped to the human reference genome and to a modified reference that contains archaic alleles that are not present in the human reference as described in Peyrégne et al.,2019 (15). We also restrict our analyses to transversion polymorphisms to minimize potential biases introduced by recurrent mutations (1,2).

We find that the *Chagyrskaya 8* population split from populations ancestral to Neandertals in Europe occured ~80-100 kya (Table S7.1). Consistent with *D*-statistics, we estimate that the *Chagyrskaya 8* population separated from European Neandertals, represented by low-coverage late Neandertal genomes, more recently than the *Denisova 5* population but earlier than the *Vindija 33.19* population (Table S7.2). *Denisova 5's* population separated approximately at the same time (~20ky before she lived) from *Chagyrskaya 8* and *Vindija*

*33.19*. When using *Chagyrskaya 8* as lineage B, the confidence intervals for the F(A|B) obtained with *Vindija 33.19* and *Mezmaiskaya 1* overlap; similarly, when using *Vindija 33.19* as lineage B, the F(A|B) of *Chagyrskaya 8* and *Mezmaiskaya 1* overlap. Thus, we conclude that these three lineages separated approximately at the same time from each other, between 80 and 100 kya, consistent with that the statistics *D*(*Chagyrskaya 8, Vindija 33.19, Mezmaiskaya 1, Denisova 5*) and *D*(*Mezmaiskaya 1, Vindija 33.19, Chagyrskaya 8, Denisova 5*) do not depart significantly from 0 (Fig. S7.2).

We also estimate that *Chagyrskaya 8* lived 11.3 ky after the split from the Neandertal mother of *Denisova 11 (*Table S7.3), and ~20 ky after the split of *Chagyrskaya 8* and *Vindija 33.19* (Table S7.2). Note, though, that our estimates of population split times are associated with substantial uncertainty.

**Table S7.1: Population split times estimates between *Chagyrskaya 8* and other archaic and present-day individuals whose genomes have been sequenced to high coverage.** The split times of *Chagyrskaya 8* from other lineages are computed by using *Chagyrskaya 8* as population A (pop A) and vice versa. Uncertainty in F(A|B) is computed with a 5 Mb block-jackknife. The F(A|B) statistic results in split time estimates in terms of proportion of human-chimpanzee divergence (div %HC). Absolute times are then obtained by assuming a human-chimpanzee separation time of 13 My, corresponding to a mutation rate of ~0.5 per base per year. Times before present are obtained by adding the branch shortening estimates for population B, here reported in terms of percentage of the human-chimpanzee divergence (bs %HC).

| popA | popB | F(A\|B) (%) | ±CI (%) | div (% HC) | split from B (ky) | bs (%HC) | split (kya) | + CI (kya) | - CI (kya) |
|---|---|---|---|---|---|---|---|---|---|
| *Chagyrskaya 8* | *Denisova 5* | 36.5 | 2.0 | 0.2 | 20.0 | 0.9 | 142.5.1 | 132.8 | 161.2 |
| | *Vindija 33.19* | 32.4 | 0.6 | 0.2 | 29.5 | 0.4 | 81.2 | 79.4 | 82.8 |
| | *Denisova 3* | 12.8 | 1.1 | 2.5 | 330.9 | 0.6 | 402.8 | 378.5 | 444.8 |
| | Mbuti | 17.6 | 0.2 | 4.0 | 520.8 | 0.0 | 520.8 | 507.7 | 534.0 |
| *Denisova 5* | *Chagyrskaya 8* | 30.3 | 0.6 | 0.4 | 50.0 | 0.6 | 129.6 | 127.5 | 131.7 |
| *Vindija 33.19* | | 35.7 | 0.6 | 0.2 | 22.1 | | 101.6 | 90.4 | 102.7 |
| *Denisova 3* | | 13.4 | 0.9 | 3.1 | 403.6 | | 483.1 | 396.6 | 502.2 |
| Mbuti | | 11.1 | 0.7 | 3.5 | 453.2 | | 552.7 | 516.8 | 568.6 |

**Table S7.2: Split times of *Chagyrskaya 8* and other high-coverage Neandertals from low coverage genomes of Neandertals.** See Table S7.1 for explanations. For the low-coverage Neandertal genomes, either all reads (all) or only those presenting signs of deamination (deam) were used.

| popA | reads | popB | F(A\|B) (%) | ±CI (%) | div (% HC ) | bs (% HC ) | Split from B (ky) | Split time (kya) |
|---|---|---|---|---|---|---|---|---|
| *Mezmaiskaya 1* | all | *Denisova 5* | 36.4 | 0.7 | 0.16 | 0.94 | 20.5 | 142.9 |
| | deam | | 36.2 | 0.8 | 0.16 | | 20.9 | 143.4 |
| *Mezmaiskaya 2* | all | | 37.0 | 0.7 | 0.14 | | 18.4 | 140.9 |
| | deam | | 37.4 | 1 | 0.10 | | 13.3 | 135.7 |
| *Spy* | all | | 35.4 | 0.7 | 0.26 | | 33.8 | 156.2 |
| | deam | | 36.0 | 1 | 0.17 | | 21.5 | 143.9 |
| *Les Cottés* | all | | 36.0 | 0.6 | 0.17 | | 21.6 | 144.0 |
| | deam | | 36.6 | 0.8 | 0.15 | | 19.7 | 142.1 |
| *Goyet* | all | | 36.4 | 0.6 | 0.16 | | 20.4 | 142.8 |
| | deam | | 36.9 | 0.9 | 0.14 | | 18.8 | 141.2 |
| *Mezmaiskaya 1* | all | *Chagyrskaya 8* | 36.2 | 0.7 | 0.09 | 0.62 | 11.0 | 91.7 |
| | deam | | 36.7 | 0.9 | 0.06 | | 8.3 | 89.0 |
| *Mezmaiskaya 2* | all | | 36.9 | 0.7 | 0.06 | | 8.0 | 88.8 |
| | deam | | 36.9 | 0.9 | 0.06 | | 8.1 | 88.8 |
| *Spy* | all | | 35.6 | 0.8 | 0.17 | | 22.2 | 102.9 |
| | deam | | 36.4 | 1.1 | 0.07 | | 8.6 | 89.3 |
| *Les Cottés* | all | | 36.0 | 0.6 | 0.09 | | 11.3 | 92 |
| | deam | | 36.1 | 0.8 | 0.09 | | 11.2 | 92 |
| *Goyet* | all | | 36.1 | 0.6 | 0.09 | | 11.1 | 91.9 |
| | deam | | 36.3 | 1.1 | 0.08 | | 10.8 | 91.5 |
| *Mezmaiskaya 1* | all | *Vindija 33.19* | 33.2 | 0.7 | 0.21 | 0.34 | 26.9 | 78.6 |
| | deam | | 33.2 | 0.9 | 0.21 | | 27.1 | 78.9 |
| *Mezmaiskaya 2* | all | | 37.2 | 0.7 | 0.07 | | 9.6 | 61.4 |
| | deam | | 37.4 | 1.0 | 0.07 | | 9.4 | 61.2 |
| *Spy* | all | | 38.1 | 0.7 | 0.06 | | 7.6 | 59.3 |
| | deam | | 39.2 | 1.1 | 0.04 | | 5.0 | 56.7 |
| *Les Cottés* | all | | 36.5 | 0.6 | 0.09 | | 11.7 | 63.5 |
| | deam | | 36.4 | 0.8 | 0.09 | | 12.0 | 63.8 |
| *Goyet* | all | | 38.5 | 0.6 | 0.05 | | 6.8 | 58.6 |
| | deam | | 39.2 | 0.9 | 0.04 | | 5.0 | 56.7 |

**Table S7.3. Relationships between high-coverage archaic genomes and the Neandertal and Denisovan parents of *Denisova 11*.** Split times of the Neandertal or Denisovan parent of *Denisova 11* from population B are shown. Population A is used to compute the corrected F(A|B) as described in Slon et al.2018 (5). Briefly, we take advantage of the fact that the proportion of Neandertal and Denisovan ancestry in *Denisova 11* is known (~50%) to correct the F(A|B) statistic and estimate the split times for the Neandertal and Denisovan component separately. F(A|B) is computed in 5 Mb windows and uncertainty is computed with a 5 Mb block jackknife. To illustrate the effect of the uncertainty in age estimates, we used two different branch-shortening estimates to arrive at split time estimates before present: branch-shortening based on transversions ("transv" in column sites) and based on all sites ("all" in sites).

| popB | popA | parent | F(A\|B) | ±CI | split from B (ky) | bs | sites | split (kya) | + CI (kya) | - CI (kya) |
|---|---|---|---|---|---|---|---|---|---|---|
| *Denisova 3* | *Chagyrskaya 8* | Denisovan | 37.6 | 0.7 | 6.8 | 0.65 | all | 91.3 | 89.6 | 93.0 |
| | | | | | | 0.55 | transv | 78.8 | 77.1 | 80.5 |
| *Denisova 3* | *Vindija 33.19* | Denisovan | 37.3 | 0.7 | 7.4 | 0.65 | all | 91.9 | 90.2 | 93.6 |
| | | | | | | 0.55 | transv | 79.4 | 77.7 | 81.3 |
| *Denisova 5* | *Denisova 3* | Neandertal | 36.2 | 0.7 | 21.0 | 0.95 | all | 134.7 | 132.3 | 147.4 |
| | | | | | | 0.94 | transv | 133.1 | 140.7 | 147.9 |
| *Chagyrskaya 8* | *Denisova 3* | Neandertal | 35.8 | 0.5 | 11.7 | 0.64 | all | 94.4 | 87.5 | 97.4 |
| | | | | | | 0.61 | transv | 91.2 | 85.2 | 95.3 |
| *Vindija 33.19* | *Denisova 3* | Neandertal | 31.9 | 0.5 | 30.8 | 0.44 | all | 87.6 | 75.8 | 90.2 |
| | | | | | | 0.40 | transv | 82.6 | 70.9 | 85.2 |

#### Relationship of *Chagyrskaya 8* to present-day humans

Comparisons of *Chagyrskaya 8* and *Denisova 5* to 271 present-day human genomes from the SGDP show that modern human populations outside Africa share significantly more derived alleles with *Chagyrskaya 8* than with *Denisova 5* (*D*(*Chagyrskaya 8*, *Denisova 5*, non-African population, chimpanzee) > 2.3%, *Z* > 2.1) (Fig.S7.3a).

When comparing *Chagyrskaya 8* with *Vindija 33.19*, *D*(*Chagyrskaya 8*, *Vindija 33.19*, modern human population, chimpanzee) ranges from -2.9% to 0.4%, and with the exception of six genomes that are marginally significant with -3 < *Z* < -2, there is no significant difference between *Chagyrskaya 8* and *Vindija 33.19* in allele sharing with modern humans, although a non-significant higher derived allele sharing with *Vindija 33.19* is consistently seen (Fig.S7.3b).

Similarly, a non-significant higher derived allele sharing with *Vindija 33.19* than *Chagyrskaya 8* can be observed for the non-African populations of the 1000 Genomes dataset (Fig.S7.4).

Neandertal introgressed alleles are expected to segregate at lower frequencies in recipient modern human populations, due to the fact that the fraction of Neandertal ancestry is low and the time since admixture is insufficient for a large fraction of alleles to drift to higher frequencies. Therefore, the power to detect which of two Neandertals, N1 and N2, is more closely related to the introgressing population can be increased by limiting the calculating *D*(N1, N2, non-African, outgroup) to low-frequency derived alleles in the non-African population (1).

We pooled the 1000 Genomes data per geographical region (super population in Fig.S7.4). The statistic *D*(*Chagyrskaya 8, Vindija 33.19,* modern human super population, chimpanzee) was then calculated for bins of derived allele frequency, by pooling the 1000 Genomes data per geographical region (super population in Fig.S7.4). At frequencies below 0.15, *D* is significantly negative for all non-African populations ($D < -7.0\%$, $Z < -2.7$, Fig.S7.5 and Fig.S7.6), a signal that is most pronounced at frequencies below 0.05 ($D < -13.0\%$, $Z < -7.7$, Fig.S7.5). This is consistent with *Vindija 33.19* being more closely related to the introgressing Neandertal populations than *Chagyrskaya 8*. At the lowest frequency bin, *Vindija 33.19* also shares more derived alleles with Africans than *Chagyrskaya 8* does. This signal is significant when including African-Americans to the group of populations with African ancestry (labeled "AFR", Z=-4.90), but not when African-Americans are excluded ("AFR2", Z=1.97). The tendency of African populations to share more low-frequency alleles with *Vindija 33.19* than *Chagyrskaya 8* may be explained by gene flow from Neandertal-admixed western non-Africans to Africans[1].

To explore whether a population related to *Chagyrskaya 8* contributed additional DNA to some present-day populations, we subdivided previously detected introgressed fragments into fragments detected only in individuals from East Asia, Europe, India and Oceania (16). We find no differences in the allele sharing to *Vindija 33.19* and different Neandertal genomes using such fragments and the statistic *D*(*Vindija 33.19*, Neandertal, introgressed fragments, chimpanzee) (Fig.S7.7a) These results are stable also when *Denisova 5* is used as outgroup. Similarly, inintrogressed fragments shared between East Asian and European populations and fragments found in only one of the two populations, no difference in allele sharing to available Neandertals is seen (Fig.S7.8).

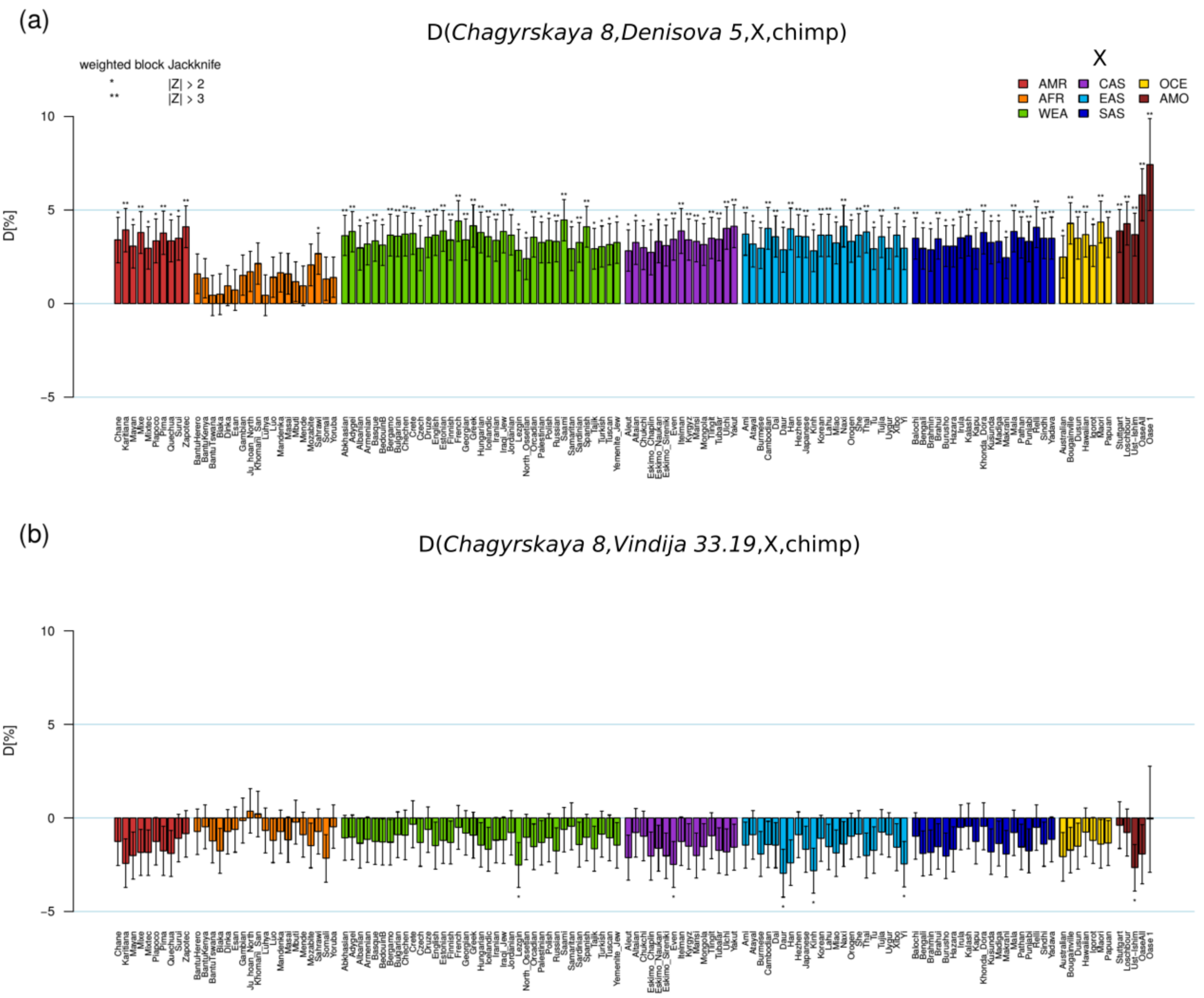


**Figure S7.3: Relative derived allele sharing (*D*-statistic) with present-day humans for *Chagyrskaya 8* and *Denisova 5* (a); and *Chagyrskaya 8* and *Vindija 33.19* (b).** Population X is one of the SGDP populations (x-axis): AMR: American; AFR: Africans; WEA: Western Eurasians; CAS: Central Asians/Siberians; EAS: East Asians; SAS: South Asians; and four ancient modern humans (AMO). Error bars indicate standard errors (1 SE).

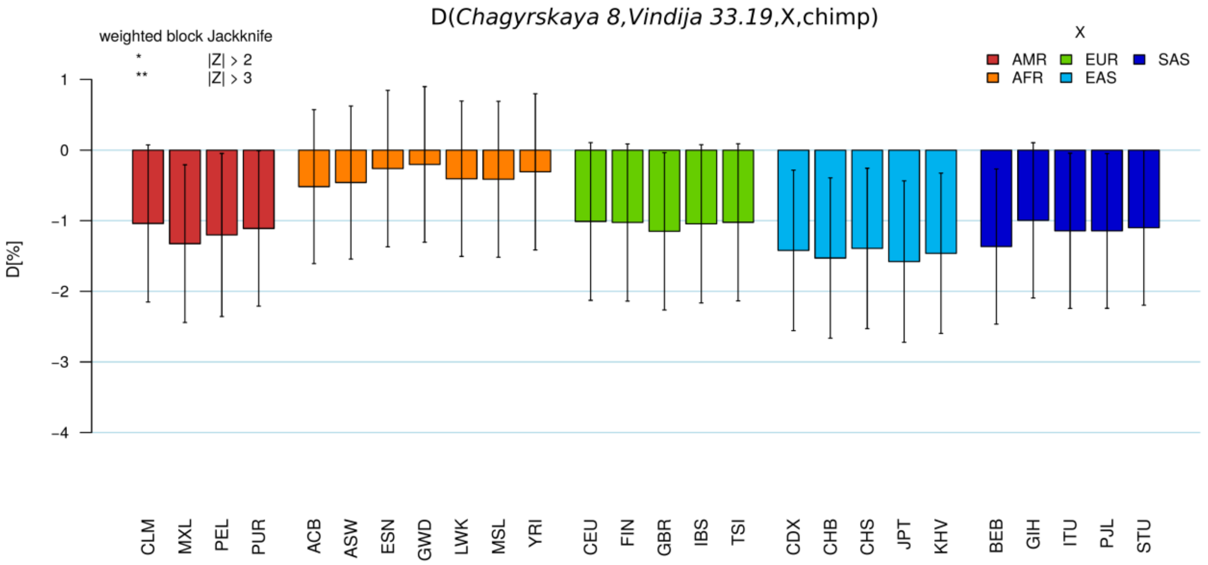


**Figure S7.4: Relative derived allele sharing (*D*-statistic) with present-day humans for *Chagyrskaya 8* and *Vindija 33.19*.** 1000 Genomes populations: AMR: American; AFR: Africans; EUR: Europeans; EAS: East Asians; SAS: South Asians. For abbreviations of individual populations, see http://www.1000genomes.org/category/population/. Error bars indicate standard errors (1 SE).

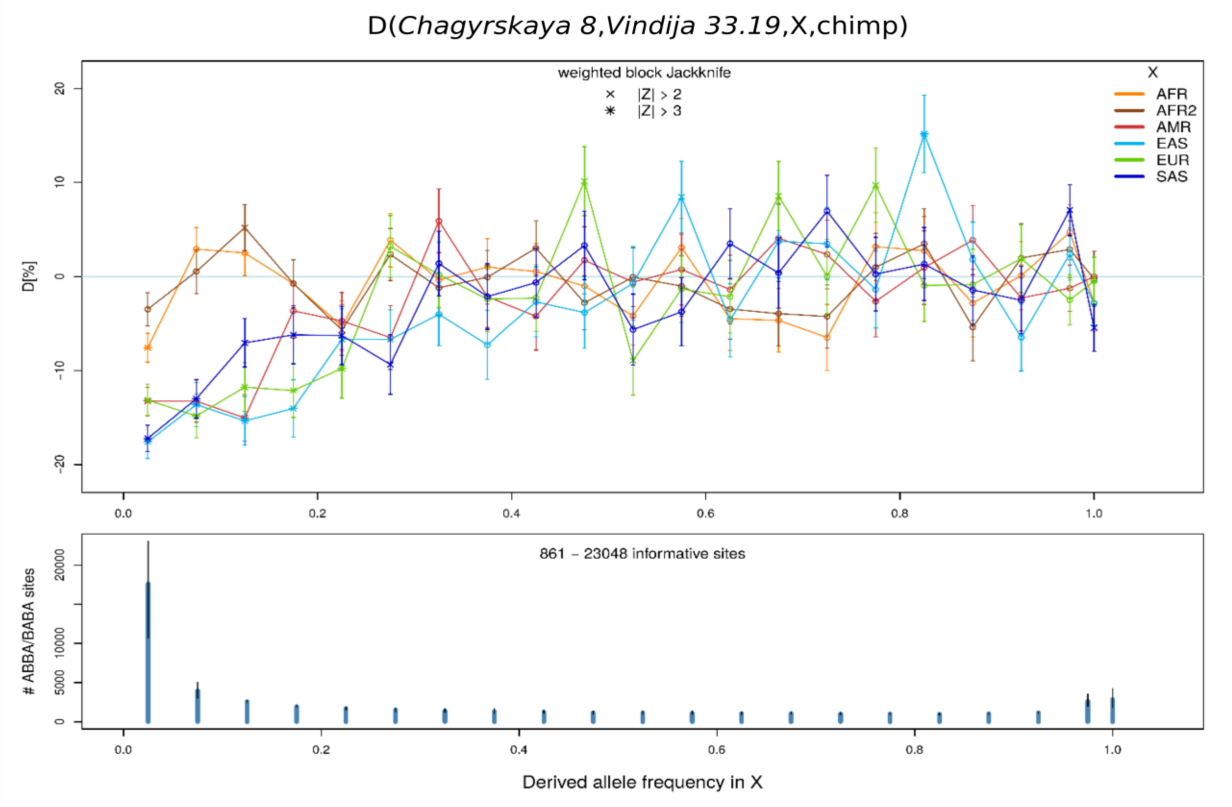


**Figure S7.5: *D*-statistic stratified by the derived allele frequency in 1000 Genomes population X.** In the top panel, error bars indicate 1 SE. Population abbreviations for X: AMR: Native Americans; AFR: Africans; AFR2: Africans without African-Americans; EUR: Europeans; EAS: East Asians; SAS: South Asians. The bottom panel shows the number of informative sites, *i.e.* sites where *p(ABBA)* > 0 and/or *p(BABA)* > 0. The blue bars show the average number across all populations X, and the black error bars show the whole range.

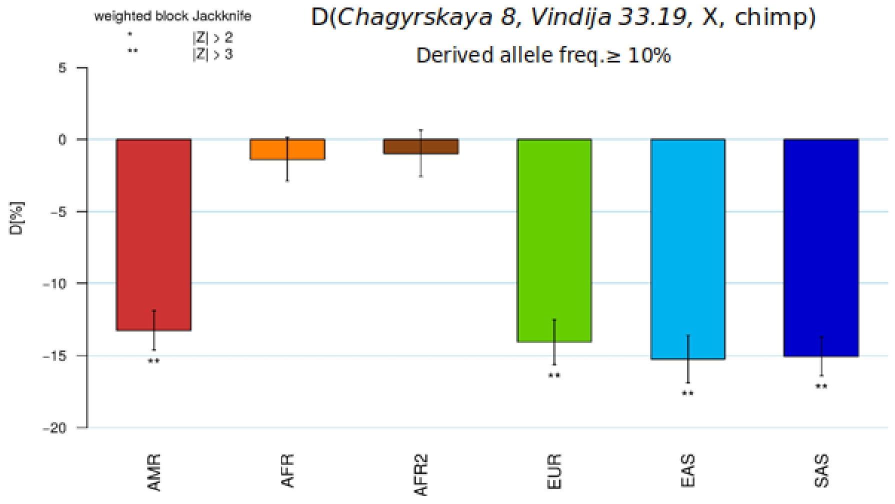


**Figure S7.6: *D*-statistic for derived allele frequencies ≤ 10% in 1000 Genomes populations.**

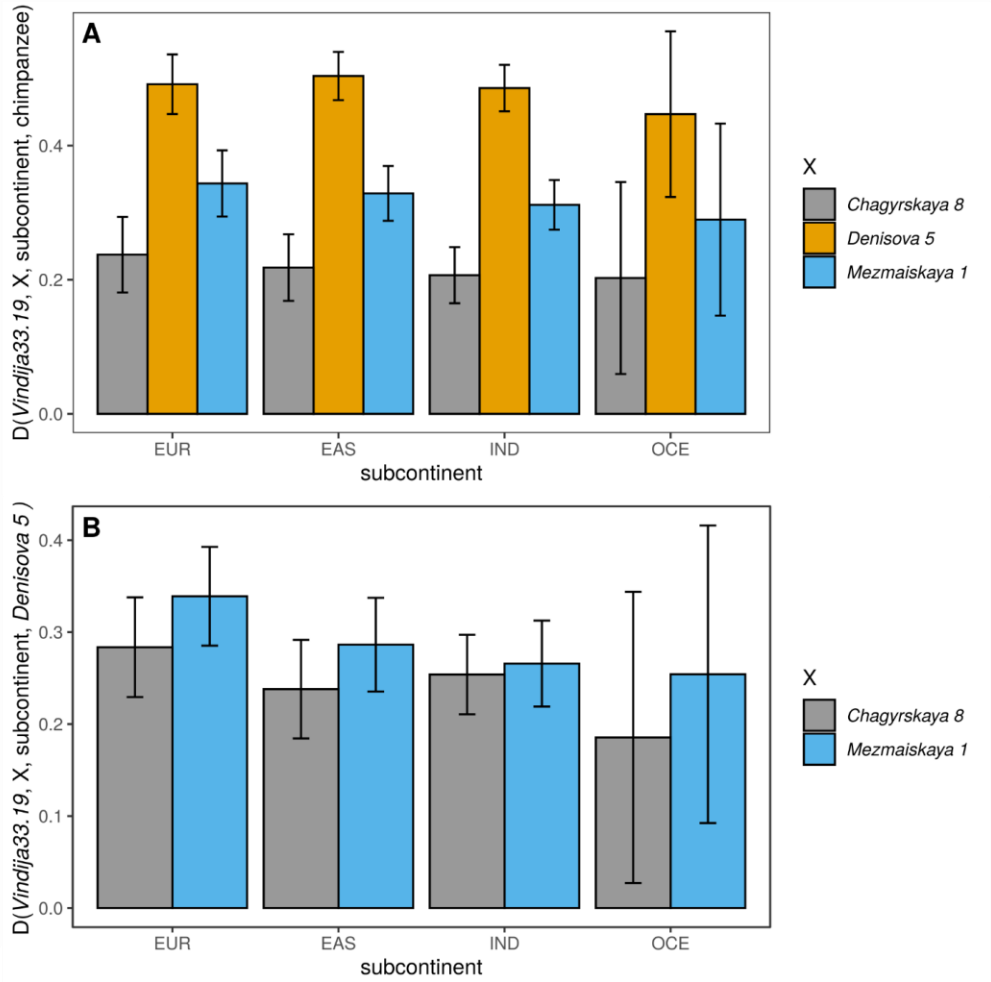


**Figure S7.7: *D*-statistic for population-specific introgressed fragments in East Asian (EAS), European (EUR), Indians (IND) and Oceanian populations (OCE).** The outgroups used are chimpanzee (A) or *Denisova 5* (B).

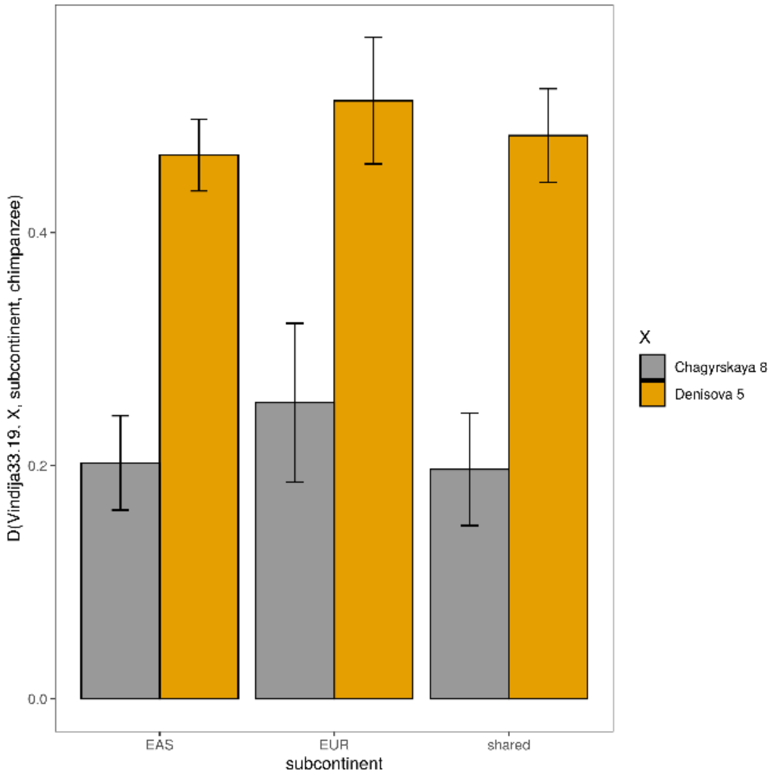


**Figure S7.8: *D*-statistic for introgressed fragments seen in East Asian (EAS), European (EUR), and fragments shared between Europeans and East Asians (shared).**

##### Joint estimation of split times, migration rates and age of the archaic genomes

We used the three high coverage Neandertal and the *Denisova 3* genomes, and the Yoruba, Han Chinese and Tuscani genomes from the SGDP dataset, to co-estimate effective population sizes, split time and ages of the archaic specimens using the software momi2 (17). For all genomes we masked sites that do not pass recommended quality filters (defined in SI3) (1) for all archaic genomes. The chimpanzee genome was used to define ancestral and derived alleles. The software momi2 computes the likelihood of a given model by fitting the joint site frequency spectrum of the different populations. We performed an optimization using the stochastic optimization function stochastic_optimize, subdividing the genome in batches of 20.000 SNPs. Confidence intervals were estimated for each model with 100 bootstraps. To estimate absolute dates we assumed a mutation rate of $1.45 \times 10^{-8}$ bp/generation and a generation time of 29 years (14).

We modeled each archaic genome as a representative of a separate population. We tested combinations of five admixture events: introgression from Neandertals to out-of-Africans, from Neandertals to Denisovans, from modern humans to Neandertals, from a superarchaic hominin (which separated from the lineage leading to modern humans more than $10^6$ years ago) to Denisovans, and from Europeans to Africans. The first introgression event, from a Neandertal population to out-of-Africa modern human populations, was included in all tested models and the relationship of the introgressing Neandertal population to the other Neandertal genomes was estimated as the split time from the lineage leading to *Vindija 33.19*. All split times, population sizes and ages of the samples were co-estimated from the data, using the constraints shown Supplementary Table 7.4. Effective population sizes were set as constant for each external and internal branch of the demography.

For all our models, estimates of split times, age of the archaic genomes and population sizes are not significantly different from the ones estimated with F(A|B), branch shortening and *PSMC* for the archaic genomes, respectively. We also estimate that 1.8-2% of the Denisova 3 genomes comes from Neandertals (2) (Fig.S7.9). In addition, we estimate a ~2% gene flow from Neandertals to modern humans, occurring from a population separating from the *Vindija 33.19* branch ~75kya.

We tested the robustness of these estimates to more complex scenarios (not shown). For example, Prüfer and colleagues (1), as well as more recent work (18), previously suggested that African populations harbor Neandertal DNA fragments, probably as the result of a back-migration from out-of-Africa populations, and that Neandertals received gene flow from modern humans (19,1). When we modeled these events, none of the split times estimates including Neandertals change significantly.

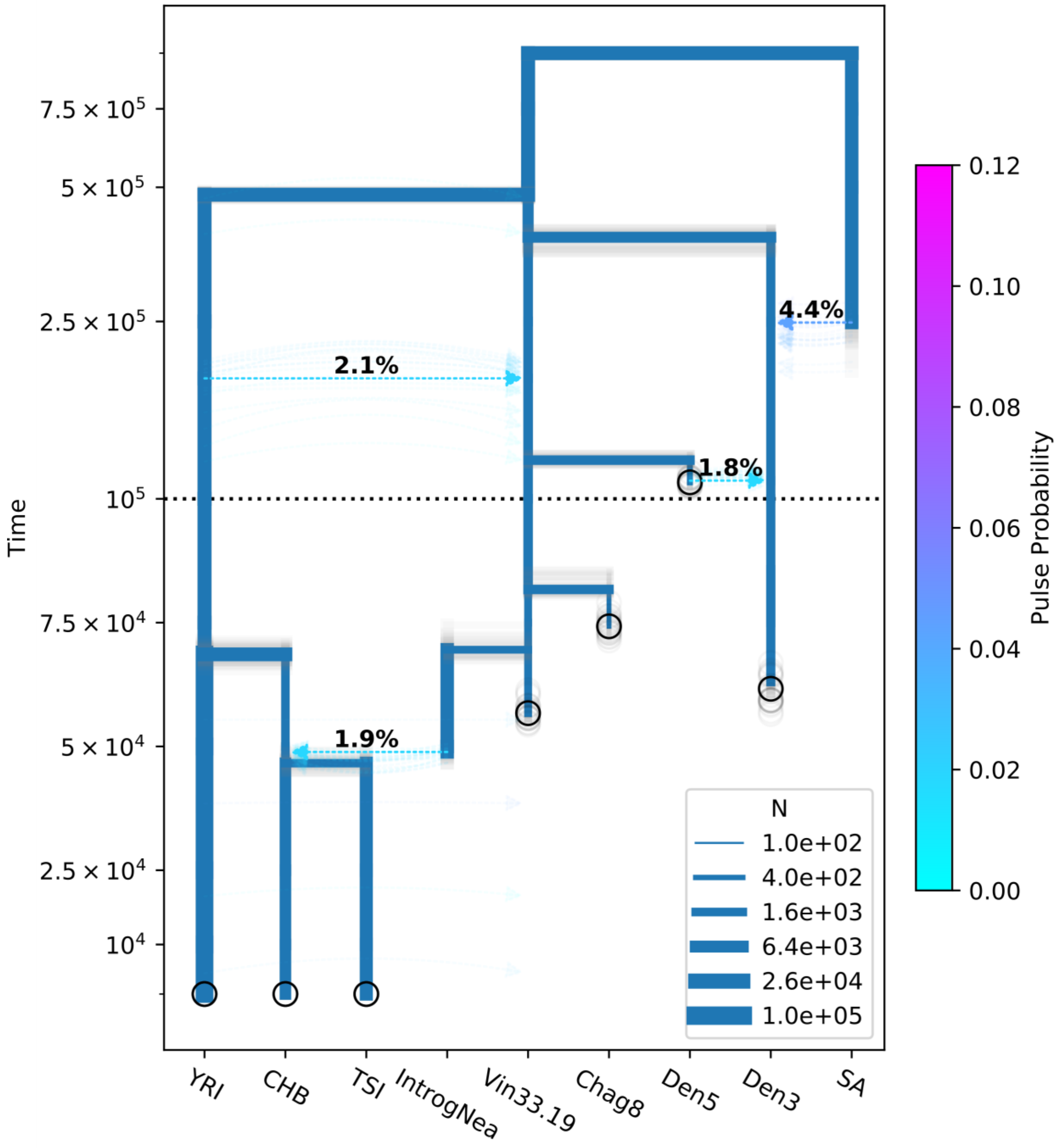


**Figure S7.9**: momi2 inferences in the presence of gene flow from Neandertals to modern humans, from Neandertals to *Denisova 3,* from a super-archaic hominin to *Denisova 3* and from Neandertals to modern humans. The thickness of the blue lines indicate population size. Colored arrows indicate introgression events and their colors the proportion of introgression. Transparent lines indicate individual bootstrap estimates. The populations reported are YRI: Yorubas, CHB: Han chinese, TSI: Tuscani, IntrogNea:

introgressing Neandertals into modern humans, Vin33.19: *Vindija 33.19*, Chag8: *Chagyrskaya 8*, Den5: *Denisova 5*, Den3: the Denisovan *Denisova 3*, SA: super-archaic hominin.

| Parameter | Lower limit | Upper limit |
|---|---|---|
| Split SA | Split Neandertals-Den3 | 1.1M |
| Split Den5-Vin33.19 | Age Den5, Age Vin33.19 | Split Neandertals-Den3 |
| Split Chag8-Vin33.19 | Age Den5, Age Chag8 | Split Neandertals-Den3 |
| Age archaic genomes | 40kya | 200kya |
| Split CHB-TSI | / | Split CHB/TSI - YRI |
| Split IntrogNea-Vin33.19 | Age Vin33.19 | / |
| Time Gene flow Nea → modern humans | Split Neandertals-modern humans, Split CHB-TSI | Split CHB/TSI - YRI |
| Time Gene flow Nea → Den3 | Age Vin33.19, Den3 | Split Neandertals-Den3 |
| Time Gene flow SA → Den3 | Split Neandertals-Den3 | / |
| Time Gene flow TSI → YRI | / | Split CHB/TSI - YRI |
| % Gene flow Nea → modern humans | 0% | 5% |
| % Gene flow Nea → Den3 | 0% | 20% |
| % Gene flow SA → Den3 | 0% | 10% |
| % Gene flow TSI → YRI | 0 | 20% |

**Table S7.4**: Parameters constraints used for the optimization of the momi2 models.

### Supplementary Information 8

#### Population structure of archaic hominins

**Summary:** We identify long tracts of homozygosity in archaic and modern human genomes. The abundance and length of homozygous tracts in the *Denisova 5* and *Chagyrskaya 8* Neandertals genomes suggests that these individuals lived in smaller groups than *Denisova 3* and modern humans. In particular, using coalescent simulations we estimate that Siberian Neandertals lived in groups smaller than 60 individuals. Heterozygosity outside homozygous tracts furthermore indicates that the long-term effective size of Neandertals was substantially smaller than that of modern humans.

##### Identification of tracts of homozygosity

Following a previously described method (1), we identify regions that are likely homozygous by descent (HBD), defined here as regions >2.5cM that are nearly devoid of heterozygous calls. To allow for some errors, we use non-overlapping 50kb windows to identify regions in which the large majority of such windows are devoid of heterozygous calls. The fraction of windows devoid of heterozygotes is given by the parameter $\pi$ (e.g. $\pi = 0.9$ indicates that at least 90% of 50 kb-windows in any called HBD tract contain no heterozygous sites). The optimal value of $\pi$ required for identifying HBD tracts is estimated separately for each genome as described elsewhere (1,2). A value of $\pi = 99\%$ was estimated for the four high-coverage archaic genomes.

To convert physical distance to genetic distance, we used a constant recombination rate of 1.3cM/Mb. We note that results do not change substantially when using a human recombination map from African-American individuals (3) (Figures S8.1, S8.2).

About 12.9% of the *Chagyrskaya 8* genome is covered by HBD tracts of size 2.5cM to 10cM. Such HBD tracts cover 5.7% and 6.2% of the *Denisova 5* and *Vindija 33.19* Neandertal genomes, respectively, and 2.6% of the *Denisova 3* genome. Longer HBD tracts (>10cM) account for 6.4% of the *Chagyskaya 8* genome, 10.5% of the *Denisova 5* genome (1) and are almost absent from the *Vindija 33.19* and *Denisova 3* genomes.

We estimated HBD tracts for the individuals in the Simons Genome Diversity Project (SGDP) (4) and four ancient modern humans (5-7) sequenced to high-coverage (Figures S8.1,

S8.2). Almost all modern humans have lower proportions of their genome covered by HBD tracts than Neandertals, especially for intermediate tracts. Exceptions include some individuals from the Karitiana and Surui, two Amazonian populations (Figure S8.3).

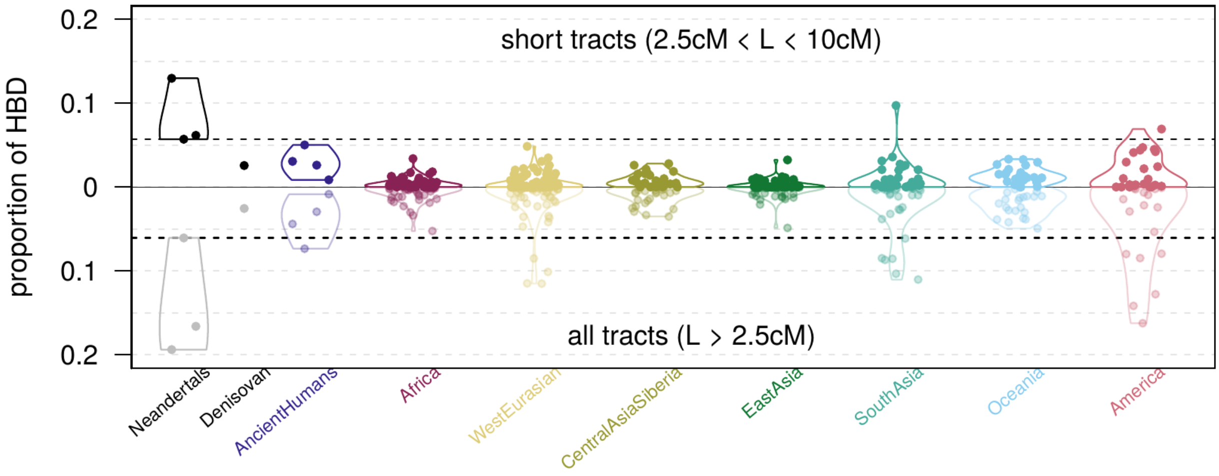


**Figure S8.1: Proportion of the genome within HBD tracts calculated using a constant recombination rate.** The dark (top) and light (bottom) panels denote the proportions of the genomes in HBD tracts between 2.5 and 10cM, and all HBD tracts >2.5cM, respectively. The dashed lines indicate the lowest value estimated for a Neandertal genome.

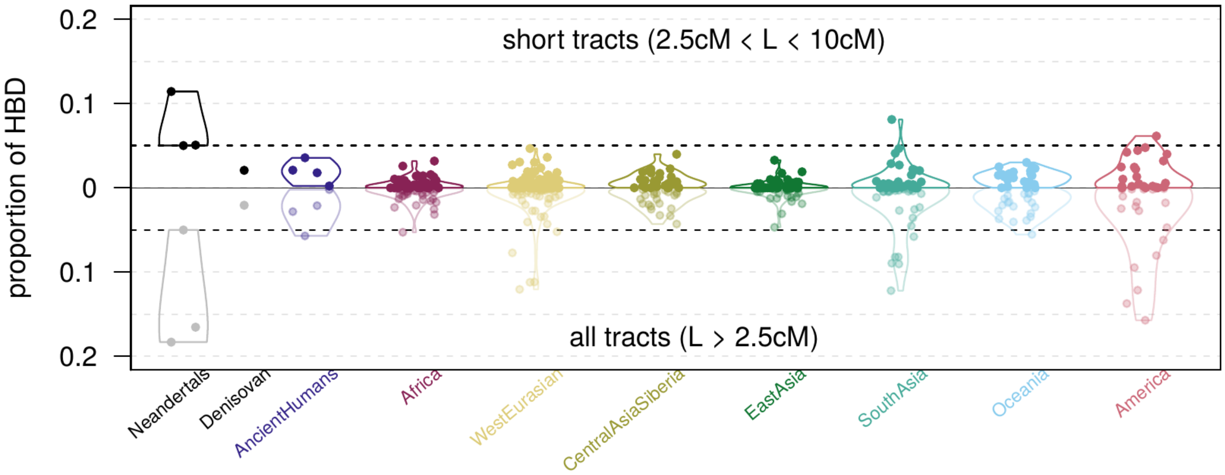


**Figure S8.2: Proportion of the genome in HBD tracts using an African-American recombination rate.** The dark (top) and light (bottom) panels denote the proportions of the genomes covered by HBD tracts between 2.5 and 10cM, and all HBD tracts with length >2.5cM, respectively. The dashed lines indicate the lower value estimated for a Neandertal genome.

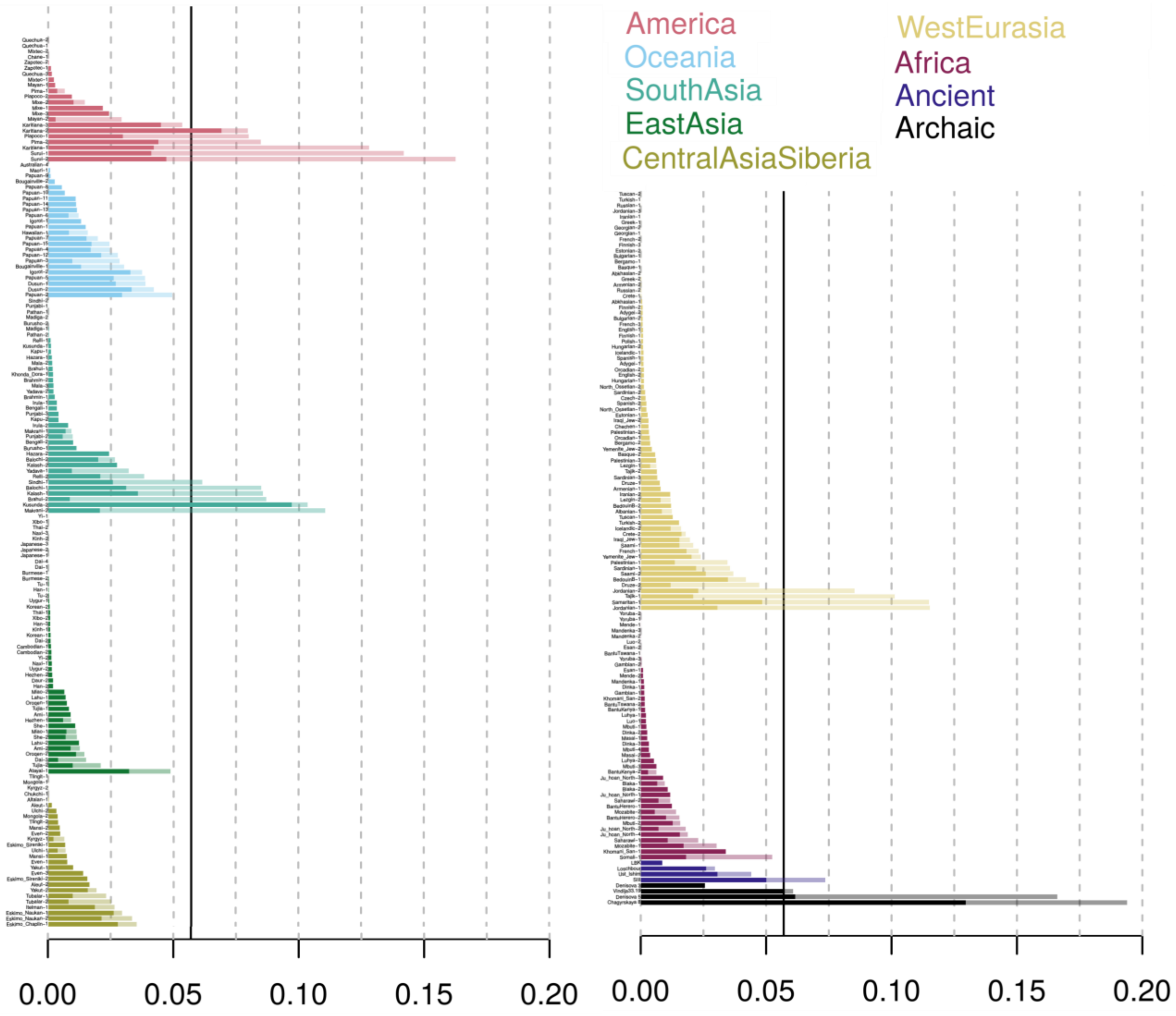


**Figure S8.3: Proportion of the genome in HBD tracts for each individual used in Figure S8.1.** Dark and light colors indicate the proportions of intermediate (between 2.5 and 10cM) and long (longer than 10cM) HBD tracts, respectively. The black vertical line indicates the lowest value estimated for a Neandertal genome for the proportion of intermediate HBD tracts.

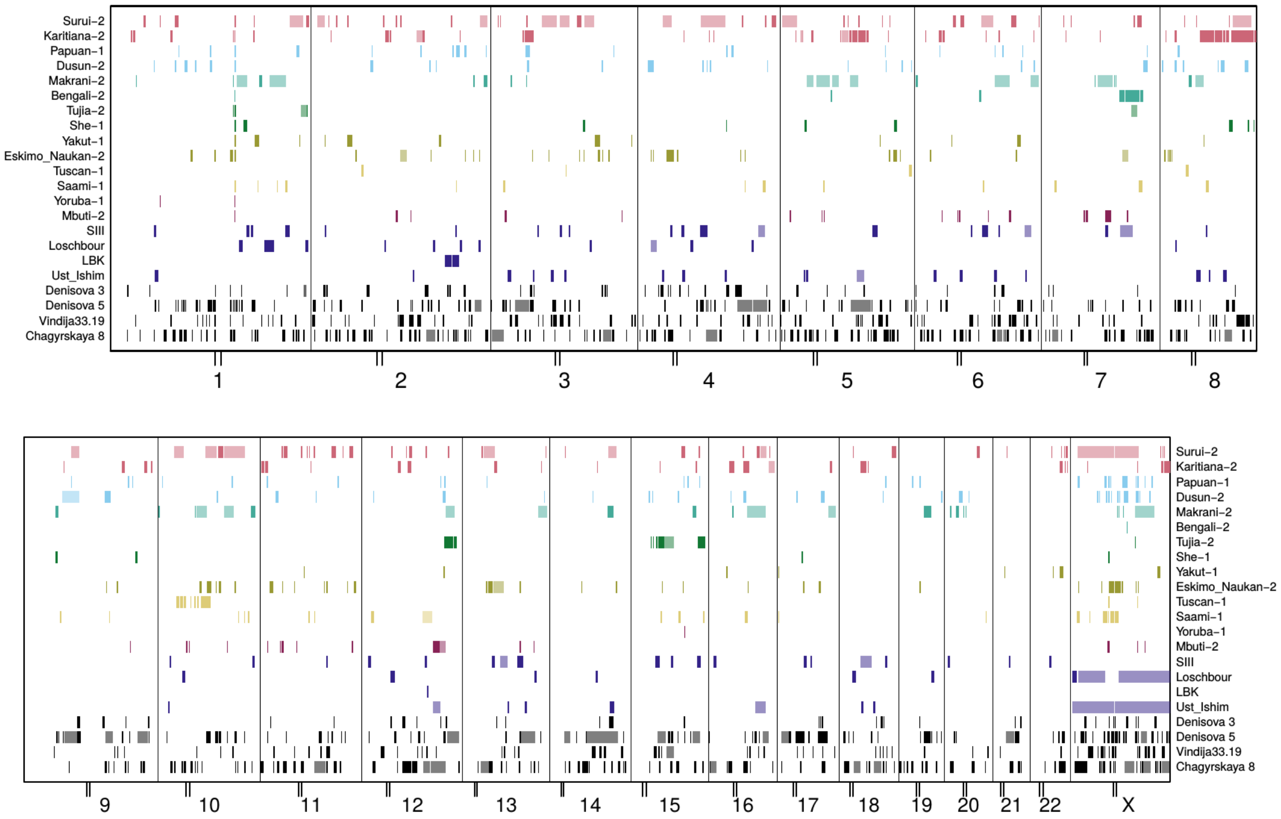


**Figure S8.4: Location of the HBD tracts along the genomes of archaic humans, ancient modern humans and 14 present-day humans** (two from each continent). The chromosomes are depicted in each box with the rectangles colored according to the groups and further divided in dark and light colors to represent intermediate (2.5-10cM) and long (>10cM) HBD tracts. Note that Loschour and Ust'Ishim are male so that no real heterozygous sites are expected on chromosome X.

#### Inference of population structure

Demographic reconstructions using Pairwise Sequentially Markovian Coalescent (PSMC)(8) indicate that the three high-coverage Neandertals experienced a low effective population size for the last ~200,000 years, including the time shortly before each individual lived (Fig. S6.2).

To test if the low effective population size could explain the large fraction of the genomes that are covered by HBD tracts, we ran coalescent simulations with the software *ms* (9) following the demography estimated by PSMC (Supplementary Information 6). The fraction of the genomes covered by HBD tracts longer than 2.5cM is much larger than in the simulations (Fig S8.5), indicating that the HBD tracts are not the direct result of population bottleneck common to all Neandertals, or a long-term low effective population size. In line with earlier results (1), we also find that the high fraction of HBD cannot be produced by a constant effective population size that would generate the heterozygosity observed in the Neandertal genomes.

Although we cannot exclude that some demographic scenarios in small panmictic populations might result in a similar abundance of HBD tracts, we here consider whether the proportion of HBD tract could be the result of highly subdivided and inter-connected populations. We model a metapopulations consisting of *D* subpopulations ("demes"), each consisting of *N* individuals. A fraction *m* of these individuals are replaced each generation by individuals migrating from other demes. Note that since the *Vindija 33.19*, *Chagyrskaya 8* and *Denisova 5* lived at different times and in different geographical locations, we do not make the assumption that they lived in the same metapopulation: we rather investigate whether the distinct populations they lived in differed in terms of structure from those of other hominins.

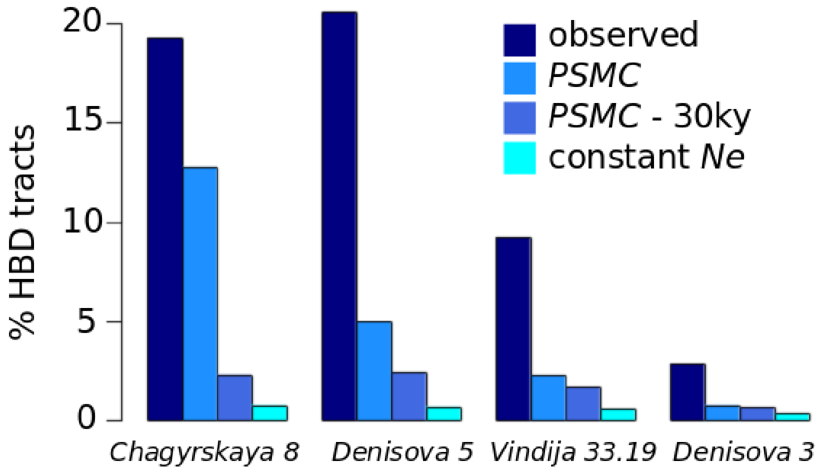


**Figure S8.5. Proportion of the genome covered by HBD tracts under different demographic scenarios.** The scenarios are: demography inferred by PSMC (Supplementary Information 6); demography inferred by PSMC up to 30,000 years before sample, and then constant, to account for the effects of recent inbreeding; constant population size with Ne to match the observed heterozygosity.

In such a scenario, the coalescent process can be modeled as occurring on two time scales (10): (i) First, when migration does not affect lineages, they coalesce within subpopulations of size N. Informally, this lasts until only one lineage is left per population, which happens when all lineages have coalesced, or when one of the remaining lineages leaves the deme by migration. (ii) When lineages migrate, the coalescent process is governed by the metapopulation with effective population size *ND*.

As the vast majority of long HBD tracts are due to recent coalescence events, these are informative about N, *i.e.* the size of the subpopulations (11). To infer the local Neandertal population structure, we use two complementary approaches: first, an intuitive analytical treatment of the first phase of coalescences within subpopulations; second, a more comprehensive simulation approach.

**Analytical approach**

*The proportion of HBD tracts in an island model*

To estimate *H(L)*, the probability that a DNA sequence of length *L* is HBD within an individual we assume that all HBD tracts are generated when they find a common ancestor within a deme uninterrupted by recombination and mutation.

Coalescent theory asserts that the two chromosomal segment an individual carries find a common ancestor with probability 1/2N per generation. The probability that one or the other segment entered the deme by migration is *2m*, where *m* is the migration rate per generation. For an HBD region to exist, it needs to remain unaffected by mutation and recombination, which we here assume to always break homozygosity. The chance segment of length *L* on one or the other chromosome to experience recombination in one generation is *2rL,* where *r* is the recombination rate per Morgan, and the probability to accumulate a mutation is *2uL,* where *u* is the mutation rate per Morgan.

With the probabilities for these events, we can now calculate *H(L),* the probability that a coalescent event occurred before migration and before recombination or mutation could affect a segment of length *L:*

$$H(L) = 1/2N / (1/2N + 2m + 2\ rL + 2uL), \qquad (1)$$

in which case the tract will be HBD. The time to the most recent common ancestor (TMRCA) for HBD tracts is exponentially distributed with mean 1/(1/2N + 2m + 2rL + 2uL).

Figure S8.6 shows that in almost all scenarios, for up to N=500 individuals, the 90%-quantile of the coalescence times for HBD tracts are shorter than 100 generations, and when the migration rate is high shorter than 10 generations, and largely independent of the length of the HBD tract. This reflects that for these scenarios, migration to the metapopulation is the primary reason for heterozygosity.

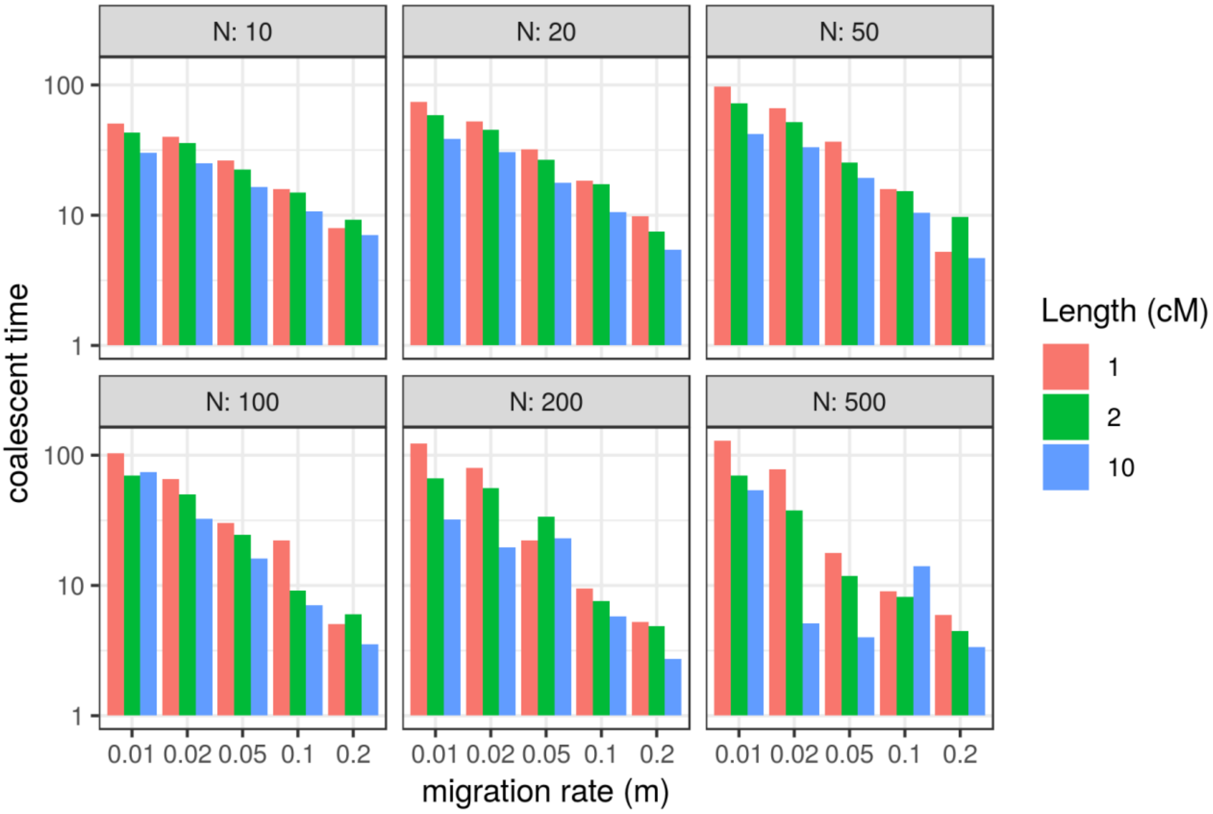


**Figure S8.6: Coalescent times of HBD tracts.** The 90%-quantile of the coalescent times of HBD fragments in simulations as a function of migration rate (*m*), DNA sequence length and deme size (N).

To validate equation (1), we compare numerical simulations based on this equation with coalescent simulations performed using *msprime* (12) (Figure S8.7). Specifically, we do $10^6$ simulations for each combination of parameters *N*, *m* and *L* in a grid of 168 parameter combinations for *L*, *N* and *m (L* from 1 to 10cM, *m* from 0.01 to 0.5, *N from 10 to 1000)*. To estimate standard errors, we performed a jackknife removing one chromosome at a time. We find that equation 1 provides a good fit to the coalescent simulations for most of the parameter space, and that the error is generally increasing with *L* and decreasing with *N* and *m* (Figure S8.7).

When we apply equation (1) to the data, we ignore mutations that break up HBD regions as described for the identification of HBD regions above. When this is implemented in our simulations, the results are qualitatively similar to the results generated by equation (1), except in cases where migration is very low (m<0.02), where the accuracy of the model decreases (Fig S8.8).

*Parameter estimation*

To estimate the population structure parameters for the Neandertals and modern humans, we need to estimate the probability that a random tract that with sequence of length *L* is homozygous by descent (HBD). We calculate this probability by counting all possible positions where a random tract may fall, and by counting the number of HBD tracts. To calculate the possible number of random regions, we consider the number of possible starting points of the tract: assume the genome has a total length of $L_{tot}$, and is subdivided into 22 autosomes, which are all longer than *L*. In this case, there are $L_{tot}$ - *22 L* possible starting points for the tract; the last *L cM* of each chromosome are invalid starting points because the end of the tract would be no longer overlapping the chromosome. The reasoning to calculate the probability that the random tract is HBD is analogous: assume we know the length of all HBD-fragments on the genome, and the length of the *i-th* tract is $L_i$. For the tract to be HBD, its starting point needs to fall in the $L_i$-*L* first bases of the HBD-fragment, otherwise it will overlap some region that are not HBD. Thus, the total number of starting points that lead to a HBD-tract is $\Sigma_{i \mid Li>L}$ ($L_i$ - L). Thus $H$(L) = $\Sigma_{i \mid Li>L}$ $(L_i – L)/( L_{tot} – 22\ L)$. As HBD-regions are non-overlapping, H(L) = Σi Hi (L), where Hi(L) is defined as in equation 1. We calculate *H* for the observed HBD regions by fitting numerically the values of L from 1 to 10 cM to numerical simulations, using the *L-BGFS-B* algorithm as implemented in the optim function in R.

The observed distribution of HBD-sequence lengths is well-captured for the *Denisova 5* and *Chagyrskaya 8* Neandertal genomes, but less so for *Vindija 33.19* genome, where the number of long HBD regions is overestimated (Figure S8.9). For the *Denisova 3* and modern human genomes the observed distribution of HBD-region lengths is inconsistent with a low effective deme size, and hence we do not estimate parameters for these genomes.

According to these analyses (Table S8.1), the *Denisova 5* genome comes from a population with effective deme sizes of ~30 individuals, with an upper 95% confidence interval of 55 individuals, and the *Charyrskaya 8* genome from a population with effective deme size between 30 and 110 individuals, with point estimate 70 individuals. Note that here we estimate the effective size of the demes, which could differ importantly from the census size in presence of high variance in reproductive success between individuals.

Since *N* and *m* have correlated effects, the composite parameter *Nm* is estimated more accurately, and is around 1 for *Denisova 5* (*Nm*=1.09, sd: 0.2) and about one half for *Chagyrskaya 8* (*Nm*=0.48, sd: 0.2), suggesting that around one migrant entered these populations per generation. For *Vindija 33.19*, we infer a large deme size and a migration rate not different from zero.

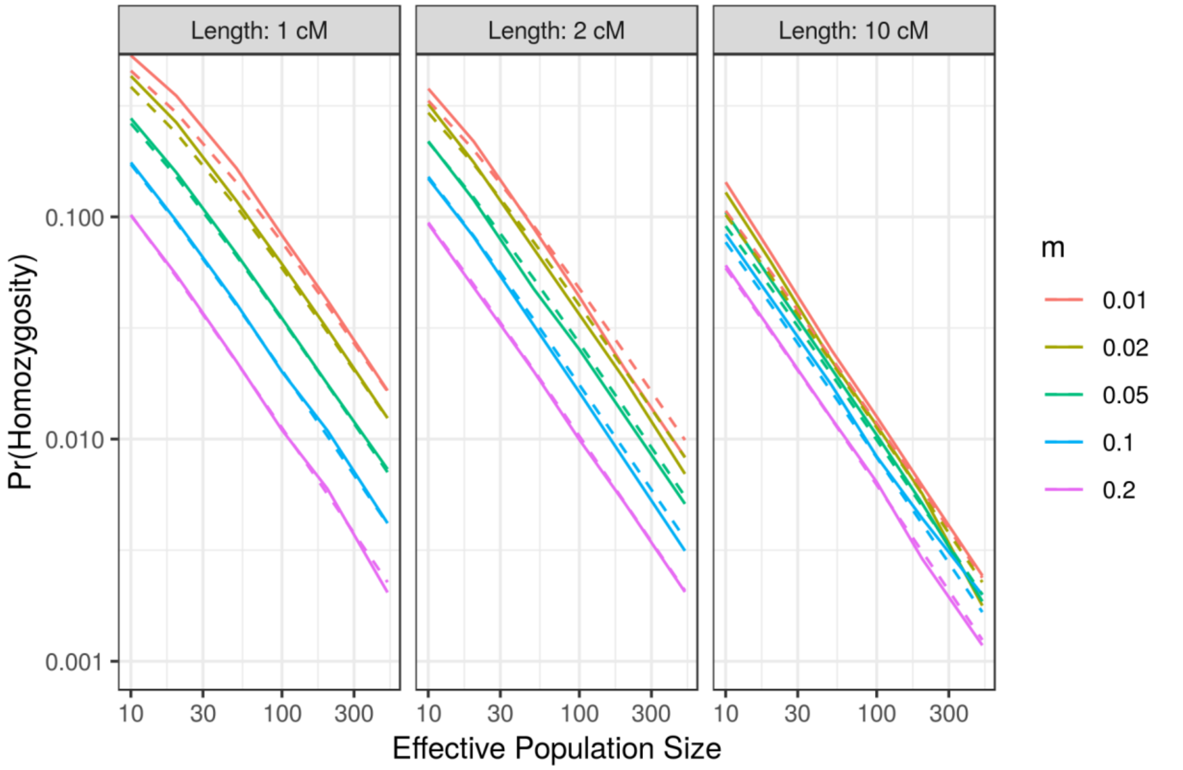


**Figure S8.7: Simulated vs. expected probability of HBD when including mutations breaking HBD regions:** The simulated (full line) and expected (dashed line, eq. 1) proportion of HBD. The fit generally decreases with sequence length, but increases with $N$ and $m$.

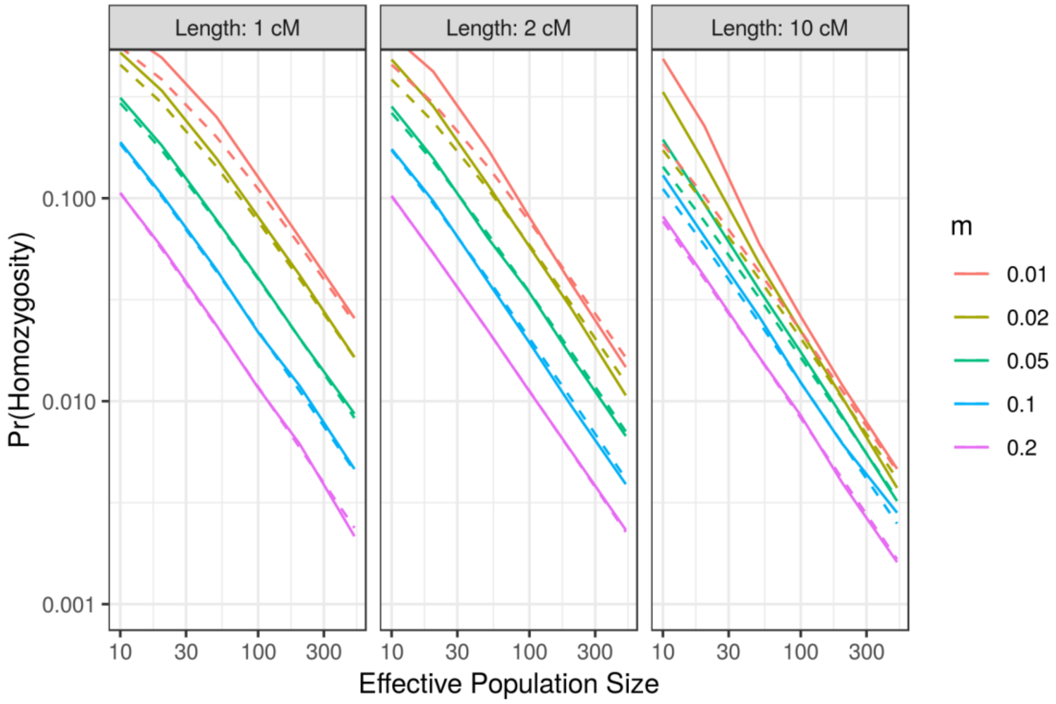

**Figure S8.8: Simulated vs. expected probability of HBD when ignoring mutations breaking HBD regions.** The simulated (full line) and expected (dashed line, eq. 1) proportion of HBD. The fit generally decreases with sequence length, but increases with *N* and *m*.

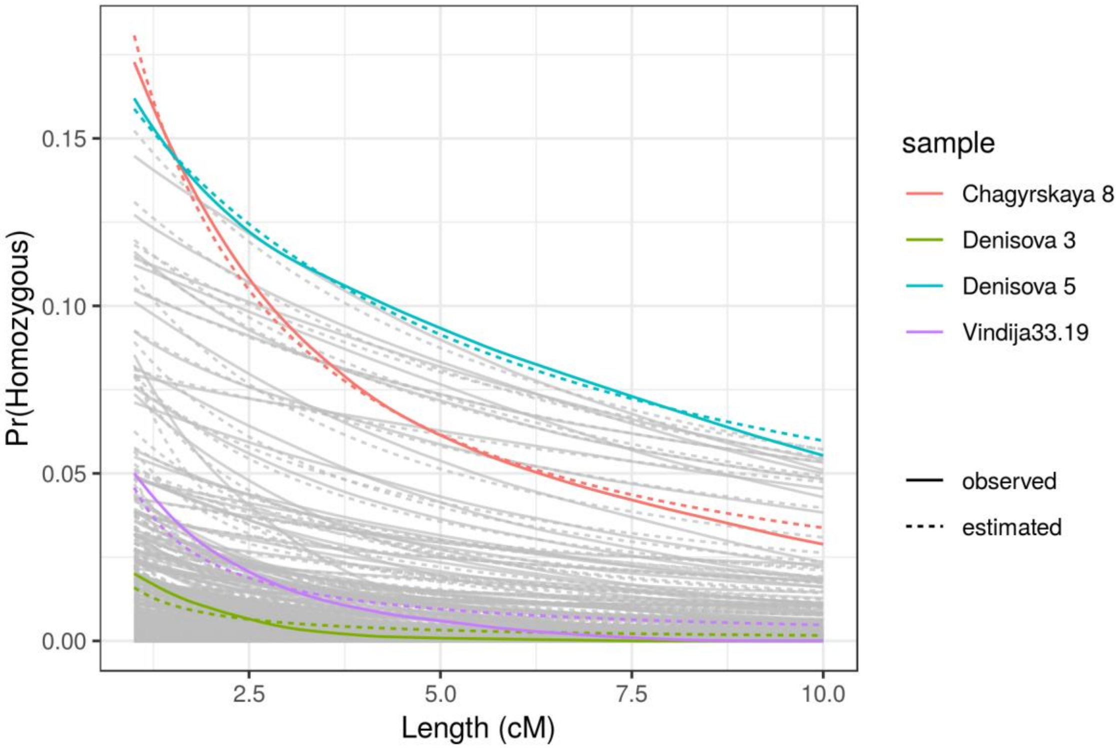


**Figure S8.9: Observed and expected probability of HBD in archaic and modern humans.** We plot the probability that a randomly chosen tract of length *L cM* is HBD. Full lines correspond to the observed data, dashed lines to the best-fit-estimate (Table S8.1).

| | **N** | **m** | **Nm** | **sd(N)** | **sd(m)** | **sd(Nm)** |
|---|---|---|---|---|---|---|
| *Denisova 5* | 30.5 | 0.036 | 1.09 | 12.8 | 0.011 | 0.20 |
| *Chagyskaya 8* | 69.8 | 0.007 | 0.48 | 20.7 | 0.004 | 0.192 |
| *Vindija 33.19* | 549.1 | 0.0 | 0 | 159 | - | - |

**Table S8.1: Estimates of deme sizes and migration rates.** Mean and standard deviation for the parameters *N*, *m*, and *Nm* for *Denisova 5*, *Chagyrskaya 8* and *Vindija 33.19*.

#### Simulation based approach

To infer the parameters *N*, *m* and *D* without the assumptions of the analytical approach, we use coalescent simulations of a finite island model. These coalescent simulations are used to approximate a likelihood surface for estimating the parameters.

We performed coalescent simulations using *ms* (9), fitting the proportion of the genome that is covered by HBD tracts of 2.5cM-10cM, or >2.5cM. We simulated $10^4$ full genome simulations for 50x50x50 combinations of values of the parameters *N*, *D*, and *m* and estimated the probability distribution *X* of observing a proportion *x* of HBD tracts of length 2.5cM-10cM, or >2.5cM, for given combinations of parameters. For large values of *m* and *N*, HBD tracts are rarely generated, so that *x* often equals *0*. To take this into account, we assumed that *x* follows a 0-inflated β-distribution (*i.e.* the random variable *X* can be written as $X= Q\,B + (1-Q)Z$, where *Q* follows a Bernoulli distribution, B a Beta distribution, and Z takes the value 0 with probability 1). To estimate the relationship between the parameters *N*, *D* and *m*, and the coefficients of the 0-inflated β-distribution, we used a 0-inflated β-regression framework (R package *gamlss*). The inferred model predicts simulated values well ($r^2$=98.6 and $r^2$=98.7 for 2.5cM<HBD<10cM and HBD>2.5cM, respectively).

To test whether ancient demography influences the proportion of HBD in a genome, we generated 100 whole-genome simulations for different combinations of *N*, *m* and *D* (Fig S8.10), and different scenarios of ancient population history. Specifically, we generated demographic scenarios in which an ancestral population of size $N_{anc}$ separates into *D* demes with migration rate *m* and size *N* for *t* generations. Simulations with *t*=100 generations and $N_{anc}$=1,000 or $N_{anc}$=10,000 could not be differentiated from each other (p>0.05, Wilcoxon rank test) or from the 0-inflated β-regression model for any of the combinations of parameters tested. This result indicates that our model is robust to differences in ancient demography.

We then estimated the likelihood of observing HBDs in proportions similar to what is observed in the archaic, ancient modern and present-day genomes. We notice that the deme size *N* and the migration rate *m* cannot be easily estimated independently because smaller groups have similar HBD patterns as more isolated ones (Fig S8.11). Long HBD are the result of coalescent events within a single deme and the total number of demes has only a modest effect, as can be observed comparing scenarios for D=20 and D=100 (Fig S8.12).

When all three high-coverage Neandertal genomes are considered together, we estimate that Neandertals had demes smaller than 50 individuals, regardless of the value of *m* and *D* (Fig S8.12). The estimated deme sizes *N* and migration rates *m* of ancient and present-day modern humans are larger (Fig S8.12, Fig S8.13, Table S8.2). Note, however, that when the Amazonian Karitiana are analyzed independently, and HBD tracts longer than 10 cM are used for the analyses (only HBD tracts > 10cM, likelihood ratio test p=0.45; all HBD tracts > 2.5cM, likelihood ratio test p=0.15) then Karitiana and Neandertals do not differ statistically significantly in their estimated parameters. Note, however, that Karitiana and Neandertals differ significantly in the absolute proportion of HBD tracts and in particular of intermediate ones (HBD tracts >2.5cM and <10cM). This pattern could be explained by a recent strong bottleneck in Karitiana.

For the intermediate tracts, we also tested a model in which Denisovans and Neandertals lived in populations with different parameters *N,D* and *m*, versus a model in which the three parameters are the same for both populations. Denisovans are statistically distinguishable from the three Neandertals (likelihood ratio test, degrees of freedom 3, *p*-values<0.05) (Fig S8.14, Table S8.2). This may indicate that deme sizes were smaller and migration rates smaller in Neandertals than in Denisovans. However, we caution that this inference is based on a single Denisovan genome (*Denisova 3)* and that long HBD tracts events are susceptible to stochasticity. At least one further Denisovan genome will be needed to infer differences in population structure between the Neandertals and Denisovans.

Note that when analyzing all HBD tracts longer than 2.5cM together, *Vindija33.19* shows a distinct pattern from the two Neandertals from the Altai region, although not significantly different (likelihood ratio test, degrees of freedom 3, *p*-values=0.054, HBD tracts>2.5cM, Table S8.2).

**Heterozygosity corrected for local population structure**

Table S8.3 gives the heterozygosity of the four archaic genomes and four ancient modern humans, both genome-wide and outside the HBD tracts.

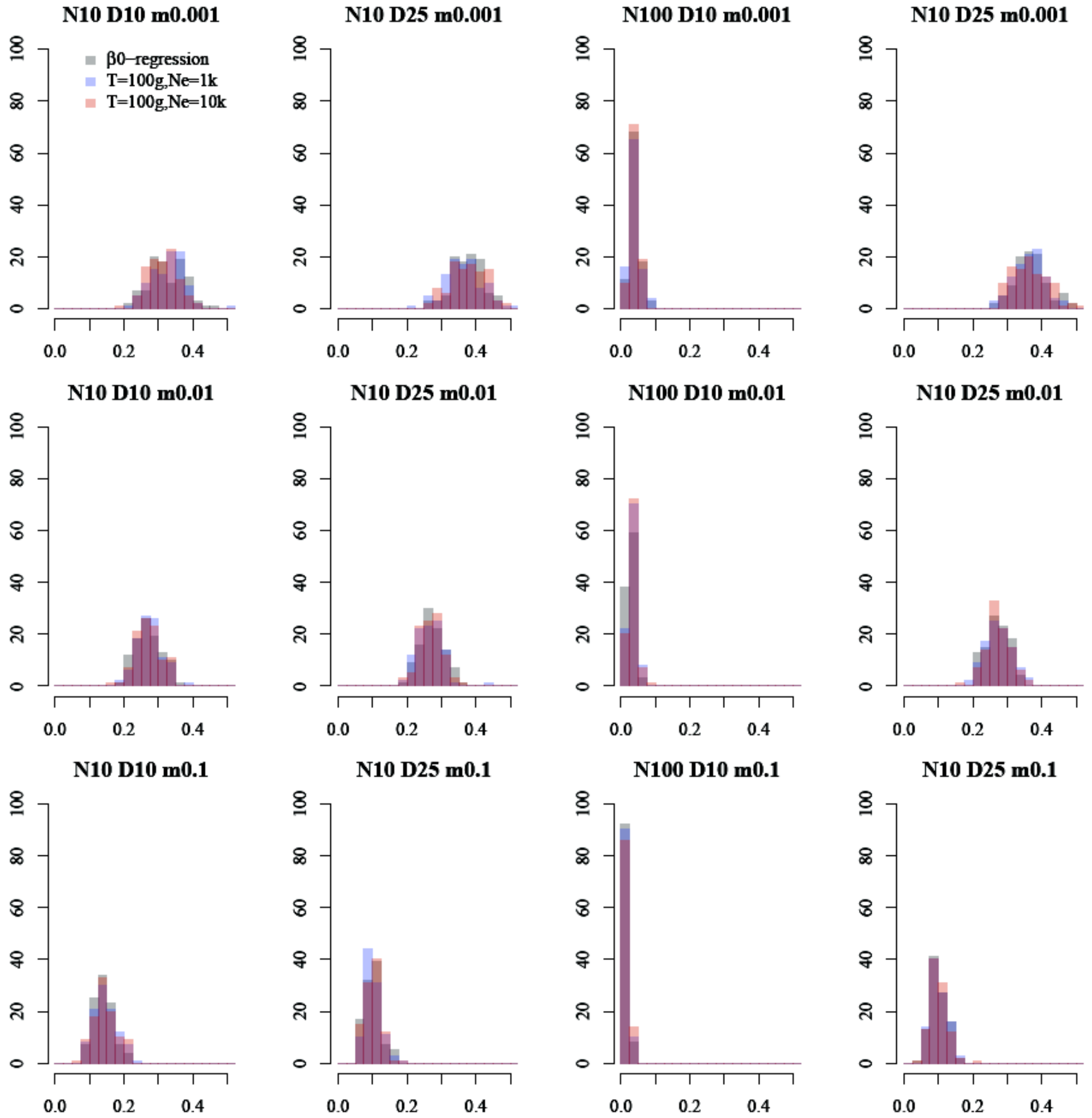


**Figure S8.10. Effects of ancient demography on HBD tracts of length >2.5cM and <10cM.** Simulations with identical metapopulation parameters (shown above) but different ancient demographies. A large ancient population size ($N_{anc}$=10,000, red) and a smaller one ($N_{anc}$=1,000, blue) are indistinguishable from the 0-inflated β-distribution model (gray) where population structure remains constant.

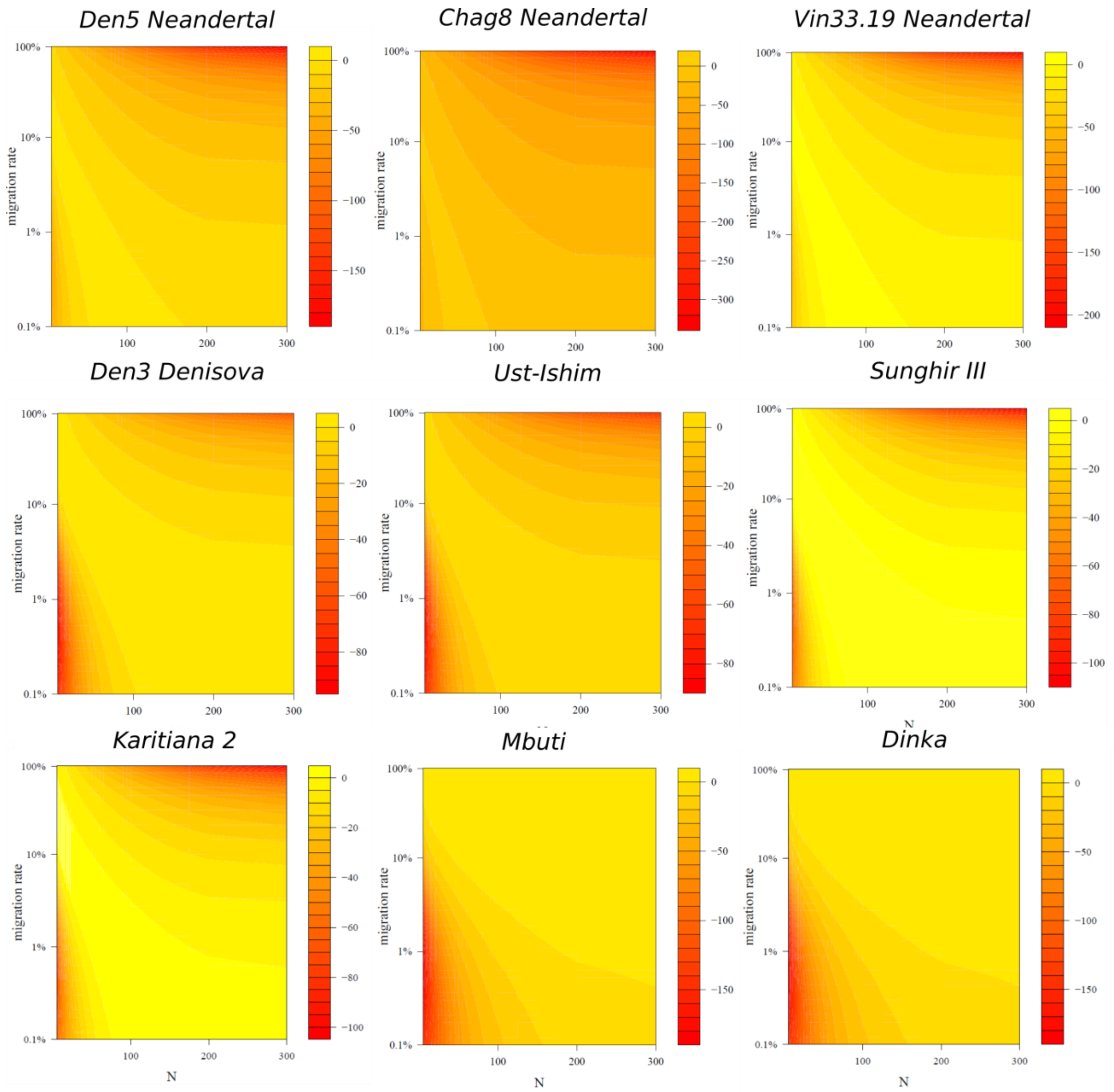


**Figure S8.11. Estimated *N* (x-axis) and *m* (y-axis) for single genomes.** Estimates are obtained using HBD tracts with length between 2.5cM and 10cM. Lighter colors indicate higher likelihoods. Note that the Karitiana individual estimates are similar to those of Neandertals.

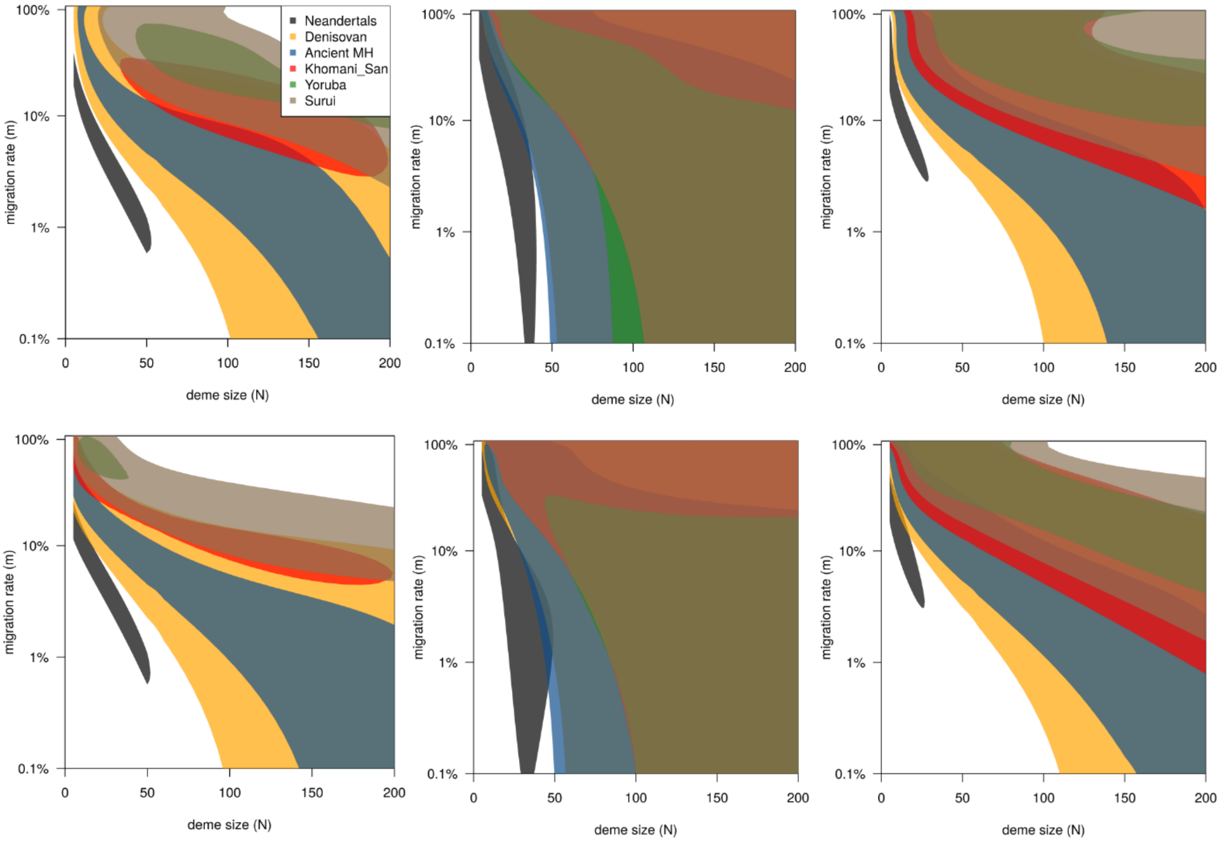


**Figure S8.12. Estimated *N* (x-axis) and *m* (y-axis) for Neandertals, Denisovans, ancient modern humans (Ancient MH) and present-day populations from the Simons Genome Diversity Project.** Estimates obtained using all HBD tracts between 2.5cM and 10cM (left), longer than 10cM (center), or longer than 2.5cM (right), for *D*=20 (top) and *D*=100 (bottom). The colored areas delimit the 95% likelihood ratio confidence intervals.

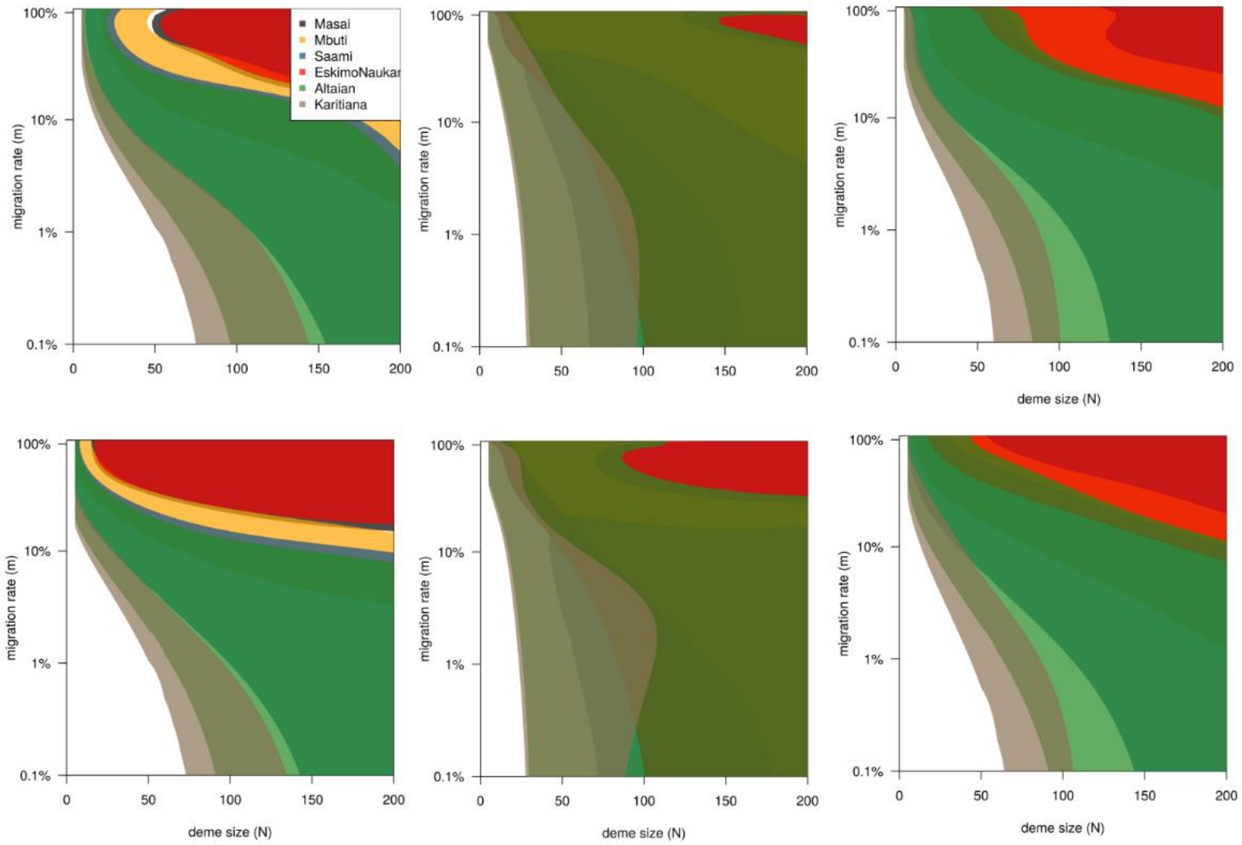


**Figure S8.13. Estimated *N* (x-axis) and *m* (y-axis) for different present-day populations from the Simons Genome Diversity Project.** Estimates obtained using all HBD tracts between 2.5cM and 10cM (left), longer than 10cM (center), or longer than 2.5cM (right), for *D*=20 (top) and *D*=100 (b,d). The colored areas delimit the 95% likelihood ratio confidence intervals.

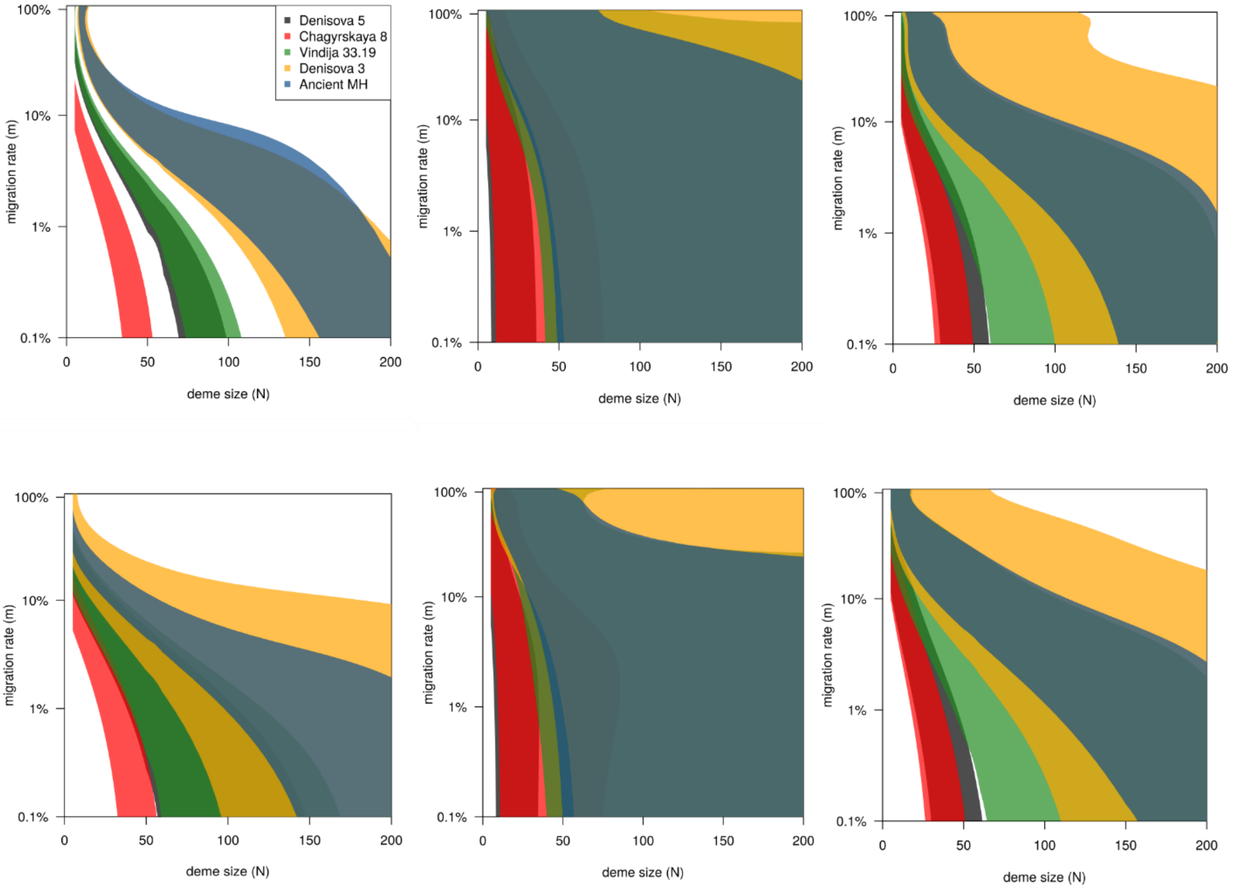


**Figure S8.14. Estimated *N* (x-axis) and *m* (y-axis) for individual archaic genomes and ancient modern humans (ancient MH).** Estimates obtained using all HBD tracts between 2.5cM and 10cM (left), longer than 10cM (center), or longer than 2.5cM (right), for *D*=20 (top) and *D*=100 (b,d). The colored areas delimit the 95% likelihood ratio confidence intervals.

| **Population 1** | **Population 2** | **2.5-10cM** | **>10cM** | **>2.5cM** |
|---|---|---|---|---|
| Neandertals | Present-day Humans | <0.001** | <0.001** | <0.001** |
| Neandertals | Karitiana | 0.0012 | 0.423 | 0.153 |
| Neandertals | Ancient Modern Humans | <0.001** | <0.001** | <0.001** |
| Neandertals | *Denisova 3* | 0.002* | 0.021* | 0.001* |
| *Denisova 3* | Ancient Modern Humans | 0.993 | 0.992 | 0.978 |
| *Vindija 33.19* | Eastern Nean. (*Chagyrskaya 8, Denisova 5)* | 0.999 | 0.154 | 0.0542 |

**Table S8.2: Likelihood ratio tests comparing different models of population structure.** The test compares a model with a single set of parameters (*N*, *m*, *D*) for two populations versus a model including different sets of parameters in the two populations (degrees of freedom=3).

| | Heterozygosity ($x10^{-4}$) | Heterozygosity non-HBD ($x10^{-4}$) |
|---|---|---|
| *Denisova 5 Neandertal* | 1.7 | 2.1 |
| *Chagyrskaya 8 Neandertal* | 1.5 | 2.0 |
| *Vindija 33.19 Neandertal* | 1.7 | 1.9 |
| *Denisova 3 Denisovan* | 2.0 | 2.0 |
| *Ust'Ishim* (6) | 7.8 | 8.1 |
| *Stuttgart (LBK)* (5) | 8.0 | 8.0 |
| *Loschbour* (5) | 5.8 | 6.0 |
| *Sunghir 3* (7) | 6.7 | 7.1 |

**Table S8.3: Heterozygosity including and excluding HBD tracts.** Heterozygosity as number of heterozygous positions per 10,000 base pair genome-wide and excluding HBD tracts longer than 2.5cM, for archaic hominins and four ancient modern humans who lived between ~8000 and ~45000 years ago in Eurasia (5-7).

### Supplementary Information 9

#### A catalog of Neandertal-specific derived changes

**Summary:** We identify genomic changes that arose and reached high frequencies on the lineage leading to Neandertals. Non-synonymous changes in protein-coding regions occur more often than expected in genes that are highly expressed during development in the striatum. Furthermore, striatum-expressed developmental genes in deserts of Neandertal ancestry in present-day people contain more non-synonymous changes than those outside of deserts. This suggests that these Neandertal changes may have been negatively selected in modern humans. An enrichment of Neandertal-specific changes can also be detected in the UTRs of striatum-expressed developmental genes. No other significant enrichment can be detected for coding regions, while other brain regions, in particular the posteroventral parietal cortex, are enriched for putatively regulatory changes.

##### Ascertainment of archaic genomic variants

We identified single nucleotide substitutions that are derived in at least one of the three high-coverage Neandertal genomes (*Denisova 5, Vindija 33.19, Chagyrskaya 8*) and ancestral in the *Denisova 3* genome. The ancestral state was inferred using the chimpanzee (*panTro4*), the gorilla (*gorGor3*) and the orangutan (*ponAbe2*) genomes. Specifically, we required that the chimpanzee and at least one of the two other ape genomes were available, that all available ape genomes shared the same state, and that the ancestral state was fixed among 37 Africans from the Simons Genome Diversity Project (1), excluding four African populations (Masai, Mozabite, Sahrawi, Somali) that have Neandertal ancestry due to European gene flow (2). This resulted in 679,437 Neandertal-specific variants. We verified that the results presented were not affected by allowing for up to 1% derived allele frequency in Africans to account for rare Neandertal introgressed alleles (data not shown).

#### Putative effects of archaic variants

We used Ensembl's Variant Effect Predictor (VEP) (3) to annotate these variants. When a variant was reported as being of a different type in different transcripts, we considered the type with the most severe consequence according to the VEP's functional hierarchy. We categorized Neandertal variants as polymorphic when at least one of the three high-coverage Neandertal genomes carried an ancestral allele or fixed when all carried the derived variant. We identified 204,933 fixed Neandertal variants, of which 2,058 overlap coding regions (CDS of protein coding genes according to annotations from Gencode (4), release 29) and 993 are fixed non-synonymous variants (Figure S9.1). The fixed non-synonymous variants affect 889 different genes.

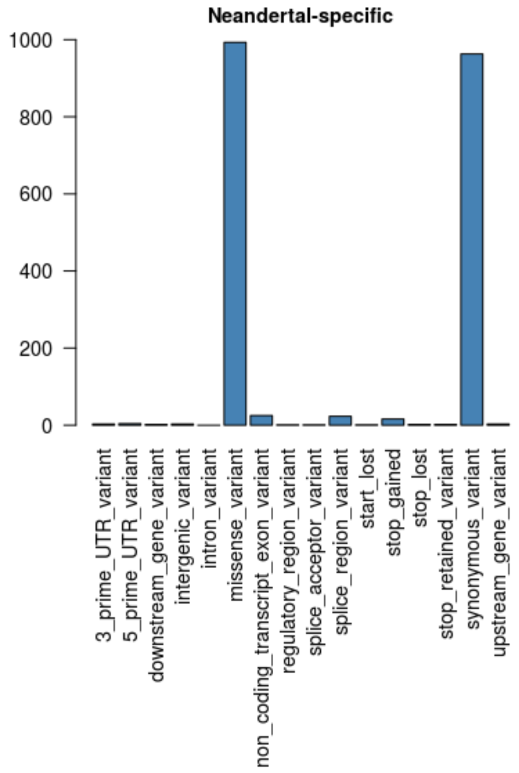


**Figure S9.1: Fixed genic Neandertal variants classified according to Ensembl's VEP.**

There are 19 Neandertal-specific loss-of-function variants (*ABHD1, AC140061.12, ANO2, ARPC4-TTLL3, C16orf96, CEP19, FCRL6, FEZ2, GJD3, MCF2L2, MMP19, OR7A5, P3H1, PLXDC1, RALGPS1, RAP1B, TRPT1, TTLL3, WDR49*).

#### Ontology analyses of protein coding variants

We performed McDonald-Kreitman tests (5) for groups of genes annotated using three different ontologies: Gene Ontology (6), Allen Brain Atlas (7,8) and Phenotype Ontology (9). The McDonald-Kreitman test uses the four counts of non-synonymous fixed differences (dN), synonymous fixed differences (dS), non-synonymous polymorphisms (pN) and synonymous polymorphisms (pS) per gene to search for outliers with a high (dN/dS) / (pN/pS) ratio that indicates positive selection on the lineage leading to the genomes analyzed.

To identify gene categories with elevated ratios, we compute the McDonald-Kreitman test for each category independently, by summing up the counts of dN, dS, pN and pS for all genes in that category and performing a 2x2 contingency table test using the R-package GOfuncR (10). To correct for multiple testing, GOfuncR computes the family-wise error rate (FWER), *i.e*. the probability that one or more of the categories with a FWER value lower than the focal one is a false positive (10). Below, we report the six categories with the lowest FWER for the Gene Ontology (Table S9.1) and Human Phenotype Ontology (HPO) (Table S9.2).

**Table S9.1: Top six categories from a GO-enrichment analysis of McDonald-Kreitman ratios of Neandertal variants.** The p-value and Family-Wise-Error Rates (FWER) are computed using the package GOfuncR.

| ontology | node id | node name | p-value | FWER |
|---|---|---|---|---|
| molecular_function | GO:0031406 | carboxylic acid binding | 14.6e-3 | 0.279 |
| biological_process | GO:0030195 | negative regulation of blood coagulation | 8.3e-3 | 0.58 |
| biological_process | GO:0050819 | negative regulation of coagulation | 8.3e-3 | 0.58 |
| biological_process | GO:1900047 | negative regulation of hemostasis | 8.3e-3 | 0.58 |
| biological_process | GO:0042339 | keratan sulfate metabolic process | 11.9e-3 | 0.738 |
| biological_process | GO:0010829 | negative regulation of glucose transmembrane transport | 11.9e-3 | 0.738 |

**Table S9.2: Top six categories from an HPO-enrichment analysis of McDonald-Kreitman ratios of Neandertal variants.** The p-value and Family-Wise-Error Rates (FWER) are computed using the package GOfuncR.

| node id | node name | p-value | FWER |
|---|---|---|---|

| | | | |
|---|---|---|---|
| HP:0007126 | Proximal amyotrophy | 1.6e-3 | 0.106 |
| HP:0002067 | Bradykinesia | 16.5e-3 | 0.719 |
| HP:0002722 | Recurrent abscess formation | 17.8e-3 | 0.758 |
| HP:0007334 | Generalized tonic-clonic seizures with focal onset | 18.2e-3 | 0.778 |
| HP:0011153 | Focal motor seizures | 19.7e-3 | 0.809 |
| HP:0002266 | Focal clonic seizures | 27.8e-3 | 0.935 |

To test for over-representation of genes with high McDonald-Kreitman ratios among genes expressed in different brain regions, we used the R-package ABAEnrichment (11) that implements the same algorithm as GOfuncR. ABAEnrichment combines the ontology of brain regions and gene expression data from the adult and the developing human brain, all provided by the Allen Brain Atlas (7,8), and annotates genes to the ontology categories, *i.e.* the different brain regions, on the basis of the gene expression in that regions. Specifically, to define whether a gene is expressed in a given brain region, ABAEnrichment considers whether the expression of a gene falls above a certain expression cutoff, *e.g.* at a 90% cutoff, a gene is considered expressed if it is among the 10% quantile of genes with the highest expression. ABAEnrichment tests expression cutoffs in 10% intervals by default, between 10% and 90%. We considered the results for these cutoffs.

For the Allen Brain Atlas, no regions had a significantly higher McDonald-Kreitman ratio than other regions in the adult brain (not shown). However, for the developing human brain, the striatum had a significantly higher (dN/dS) / (pN/pS) ratio (FWER=0.035) at the 90% expression cutoff than other regions in the brain (Table S9.3) in the age category 12-19 years (age category 4). At this cutoff, 39 genes with Neandertal-specific non-synonymous variants are expressed in the striatum (Table S9.4). To test the stability of this signal at different expression cutoffs, we repeated the ABAEnrichment test for different gene expression cutoffs between 80% and 95% in steps of 1% (Fig. S9.2). The only expression cutoff for which the striatum has FWER<0.05 is 90%. However, the striatum also shows the lowest FWER among brain regions at age category 1, for expression cutoffs between 88%-90%. In addition, the striatum does not have a lower number of genes at that cutoff compared to other brain areas (Fig. S9.3).

Note that the ratios (dN/dS) / (pN/pS) are significantly lower for Neandertals than for Africans (Wilcoxon paired rank test $<10^{-6}$) (Fig. S9.4). In contrast to Africans, for Neandertals no regions show a ratio significantly higher than 1, including the striatum at age category 4 (ratio = 1.002, p=1 using Fisher's exact test). This is possibly explained by the larger presence of slightly deleterious polymorphic variants in Neandertals (12,13), which lower (dN/dS) / (pN/pS) ratios and mask signals of positive selection (14,15).

In conclusion, while we caution that more genomes would be required to better assess the targets of positive selection in Neandertals, we suggest that genes expressed in the striatum show an intriguing increase of non-synonymous changes in Neandertals.

**Table S9.3: Top six combinations of brain-region/age-category from the ABAEnrichment analysis of Neandertal protein coding variants.** The analysis scores combinations of brain-regions and age-category on the basis of the McDonald-Kreitman ratio of the genes expressed for that combination. Age categories are defined as follows: 1=prenatal, 2=0-2 years, 3=3-11 years, 4=12-19 years, 5= older than 19 years. Results are analogous when derived variants with a frequency of up to 1% in Africans are considered (data not shown).

| age category | structure id | structure | FWER |
|---|---|---|---|
| **4 (12-19 yrs)** | Allen:10333 | STR_striatum | 0.029 |
| **1 (prenatal)** | Allen:10333 | STR_striatum | 0.341 |
| **1 (prenatal)** | Allen:10294 | HIP_hippocampus (hippocampal formation) | 0.422 |
| **1 (prenatal)** | Allen:10155 | Br_brain | 1 |
| **1 (prenatal)** | Allen:10157 | FGM_gray matter of forebrain | 1 |
| **1 (prenatal)** | Allen:10158 | Tel_telencephalon | 1 |

**Table S9.4: List of genes with changes in Neandertals and expressed in the striatum at age category 4 (12-19 years), for the 90% expression cutoff.**

| Gene | dN | dS | pN | pS | Gene | dN | dS | pN | pS | Gene | dN | dS | pN | pS |
|---|---|---|---|---|---|---|---|---|---|---|---|---|---|---|
| A2M | 0 | 1 | 1 | 0 | EVI2A | 0 | 0 | 1 | 0 | PDZD4 | 0 | 0 | 1 | 0 |
| ABCA2 | 0 | 0 | 1 | 0 | F3 | 1 | 0 | 0 | 0 | PHLPP1 | 1 | 0 | 1 | 2 |
| ACTN2 | 0 | 0 | 0 | 1 | FAM171A1 | 0 | 0 | 0 | 1 | PIGT | 0 | 0 | 0 | 1 |
| ADCY5 | 0 | 0 | 0 | 1 | FBXO44 | 0 | 0 | 1 | 0 | PINK1 | 0 | 0 | 1 | 0 |
| ADD1 | 0 | 0 | 0 | 1 | FGF1 | 0 | 0 | 0 | 1 | PJA2 | 0 | 0 | 1 | 0 |
| AHSA1 | 0 | 0 | 1 | 0 | FKBP8 | 0 | 0 | 1 | 0 | PKN1 | 0 | 0 | 2 | 0 |
| AK5 | 0 | 0 | 0 | 1 | GAS7 | 0 | 0 | 1 | 0 | PLEKHH1 | 0 | 0 | 1 | 0 |
| AKAP11 | 0 | 0 | 2 | 0 | GHITM | 0 | 0 | 1 | 0 | PLK2 | 0 | 0 | 0 | 1 |
| AKAP12 | 0 | 0 | 0 | 1 | GNAS | 1 | 0 | 0 | 0 | PMEPA1 | 0 | 0 | 0 | 1 |
| ALAD | 0 | 0 | 1 | 1 | GOT1 | 0 | 0 | 0 | 1 | PMP2 | 0 | 0 | 1 | 0 |
| ALDH2 | 0 | 1 | 0 | 0 | GPI | 0 | 0 | 1 | 1 | PPDPF | 0 | 0 | 1 | 0 |
| ALDH4A1 | 0 | 0 | 0 | 2 | GPR37 | 0 | 0 | 1 | 1 | PPP1CB | 0 | 0 | 0 | 1 |
| AMFR | 0 | 0 | 0 | 1 | GPR37L1 | 0 | 0 | 0 | 1 | PPP1R3C | 0 | 0 | 1 | 0 |
| AMPD2 | 1 | 0 | 0 | 0 | GPR6 | 0 | 0 | 0 | 1 | PPP2CA | 0 | 1 | 0 | 0 |
| ANK2 | 0 | 0 | 2 | 3 | GPX3 | 0 | 0 | 1 | 0 | PPP2R1A | 0 | 0 | 0 | 1 |
| ANLN | 0 | 0 | 1 | 1 | GRM3 | 0 | 0 | 0 | 1 | PRDX2 | 0 | 0 | 1 | 0 |
| APLP1 | 0 | 0 | 2 | 0 | GSK3A | 1 | 0 | 0 | 0 | PRKCB | 0 | 0 | 0 | 1 |
| APP | 0 | 0 | 0 | 1 | GSN | 0 | 0 | 2 | 0 | PRKCG | 0 | 0 | 0 | 1 |
| ARCN1 | 1 | 1 | 0 | 1 | GSTP1 | 0 | 0 | 0 | 1 | PRNP | 0 | 0 | 1 | 0 |
| ARHGEF9 | 0 | 1 | 0 | 0 | HADHA | 0 | 0 | 1 | 0 | PRRC2B | 1 | 0 | 1 | 0 |
| ARNT2 | 0 | 0 | 0 | 1 | HAGH | 0 | 0 | 2 | 1 | PSD2 | 1 | 0 | 1 | 0 |
| ARPC4-TTLL3 | 0 | 0 | 0 | 1 | HDLBP | 0 | 0 | 0 | 1 | PSMC4 | 0 | 0 | 0 | 2 |
| ASAH1 | 1 | 0 | 0 | 0 | HERPUD1 | 0 | 0 | 1 | 0 | PTPN5 | 0 | 0 | 0 | 1 |
| ATP1A1 | 0 | 2 | 0 | 0 | HIVEP2 | 1 | 1 | 0 | 1 | PTPRA | 0 | 1 | 0 | 0 |
| ATP1A2 | 0 | 0 | 0 | 1 | HK1 | 0 | 0 | 0 | 1 | PTPRN | 0 | 0 | 1 | 0 |
| ATP1A3 | 0 | 1 | 0 | 0 | HNRNPH2 | 0 | 0 | 0 | 1 | PTPRZ1 | 1 | 1 | 1 | 0 |
| ATP2A2 | 0 | 0 | 1 | 0 | HNRNPM | 0 | 0 | 1 | 0 | PTTG1IP | 0 | 0 | 1 | 0 |
| ATP2B1 | 0 | 0 | 0 | 1 | HNRNPUL2 | 0 | 0 | 1 | 0 | QDPR | 0 | 0 | 0 | 1 |
| ATP8A1 | 0 | 0 | 0 | 1 | HSPA12A | 0 | 0 | 1 | 0 | RAB5C | 0 | 0 | 0 | 1 |
| ATXN10 | 0 | 0 | 0 | 1 | HSPA2 | 0 | 0 | 4 | 3 | RAB7A | 0 | 0 | 0 | 1 |
| BAALC | 0 | 1 | 0 | 0 | HSPH1 | 0 | 0 | 0 | 2 | RAPGEF5 | 0 | 0 | 0 | 1 |
| BABAM1 | 0 | 1 | 0 | 0 | HTATSF1 | 0 | 1 | 0 | 0 | RBFOX1 | 0 | 0 | 0 | 1 |
| BAG6 | 0 | 0 | 2 | 2 | HTRA1 | 0 | 0 | 2 | 0 | RHBDD2 | 0 | 0 | 0 | 2 |
| BEST1 | 0 | 0 | 1 | 0 | HYOU1 | 0 | 0 | 1 | 0 | ROGDI | 1 | 0 | 0 | 0 |
| BIN1 | 0 | 1 | 0 | 0 | IDH2 | 0 | 0 | 0 | 1 | RPL3 | 0 | 0 | 1 | 0 |
| BTBD1 | 0 | 0 | 0 | 1 | IDH3G | 0 | 0 | 1 | 1 | RTN1 | 0 | 0 | 0 | 1 |
| C16orf45 | 0 | 0 | 0 | 1 | IL6ST | 0 | 0 | 1 | 0 | SAP18 | 0 | 0 | 1 | 0 |
| C17orf49 | 0 | 1 | 0 | 0 | INF2 | 0 | 0 | 0 | 3 | SDC3 | 0 | 0 | 1 | 0 |
| C1orf43 | 0 | 1 | 0 | 0 | ITFG1 | 0 | 0 | 1 | 0 | SDC4 | 0 | 1 | 0 | 0 |
| C1orf61 | 0 | 0 | 1 | 0 | ITPK1 | 0 | 0 | 0 | 1 | SDHB | 0 | 0 | 1 | 2 |
| C2orf74 | 1 | 0 | 0 | 0 | ITPR1 | 1 | 0 | 0 | 2 | SEPT5 | 0 | 0 | 1 | 0 |
| CAMK1 | 0 | 0 | 1 | 0 | KCTD17 | 0 | 0 | 3 | 0 | SERINC1 | 0 | 0 | 1 | 0 |
| CAMK1D | 0 | 1 | 0 | 0 | KIAA0100 | 0 | 1 | 0 | 0 | SERINC3 | 0 | 0 | 0 | 1 |
| CAMKK1 | 0 | 0 | 0 | 1 | KIAA0513 | 0 | 0 | 0 | 1 | SF3B1 | 0 | 1 | 0 | 0 |
| CAPNS1 | 1 | 0 | 0 | 0 | KIAA0930 | 0 | 0 | 1 | 0 | SGTA | 0 | 0 | 0 | 1 |
| CAPZB | 0 | 1 | 0 | 0 | KIF1A | 0 | 0 | 0 | 2 | SLC1A3 | 0 | 0 | 0 | 1 |
| CBR1 | 0 | 0 | 2 | 0 | KIF1B | 0 | 0 | 1 | 0 | SLC25A23 | 1 | 0 | 0 | 0 |
| CCNI | 0 | 0 | 0 | 1 | KIF3C | 0 | 0 | 1 | 1 | SLC44A2 | 0 | 0 | 0 | 1 |
| CELSR2 | 0 | 0 | 2 | 1 | KIF5A | 0 | 0 | 0 | 1 | SLC4A3 | 0 | 1 | 0 | 0 |
| CERCAM | 0 | 0 | 0 | 1 | KIFC2 | 1 | 0 | 0 | 0 | SLC9A3R1 | 0 | 0 | 0 | 1 |
| CERS2 | 0 | 1 | 0 | 0 | LANCL1 | 0 | 0 | 1 | 0 | SMARCA2 | 0 | 2 | 1 | 1 |
| CETN2 | 0 | 0 | 0 | 1 | LARP1 | 0 | 0 | 0 | 1 | SNAPC5 | 0 | 0 | 0 | 1 |
| CHRM1 | 0 | 0 | 0 | 1 | LGALS3BP | 0 | 0 | 1 | 0 | SORBS1 | 0 | 0 | 1 | 1 |
| CKAP5 | 0 | 1 | 0 | 1 | LGI1 | 0 | 0 | 0 | 1 | SPAG9 | 0 | 0 | 0 | 1 |
| CKB | 0 | 0 | 1 | 0 | LIPA | 0 | 0 | 1 | 0 | SPTAN1 | 1 | 0 | 0 | 0 |
| CLCN3 | 0 | 0 | 1 | 0 | LMBRD1 | 0 | 0 | 0 | 1 | SPTBN1 | 0 | 1 | 0 | 3 |
| CLCN4 | 0 | 1 | 0 | 0 | LPAR1 | 0 | 0 | 0 | 1 | SRRM2 | 1 | 0 | 1 | 0 |
| CLIP3 | 0 | 0 | 0 | 3 | LRP4 | 1 | 0 | 0 | 0 | SRSF2 | 0 | 0 | 0 | 1 |
| CLSTN3 | 0 | 1 | 0 | 0 | LRPAP1 | 0 | 0 | 0 | 1 | STIM1 | 1 | 0 | 0 | 0 |
| CMAS | 0 | 0 | 1 | 0 | LRRC4B | 0 | 1 | 0 | 0 | STMN2 | 0 | 0 | 1 | 0 |
| CNDP1 | 0 | 0 | 0 | 1 | LRRN1 | 0 | 0 | 1 | 0 | STMN3 | 0 | 0 | 0 | 1 |
| COCH | 0 | 0 | 2 | 0 | MAP1A | 0 | 0 | 1 | 1 | STOM | 0 | 0 | 1 | 0 |
| COPA | 0 | 1 | 0 | 0 | MAP1B | 0 | 0 | 1 | 1 | STXBP1 | 0 | 0 | 0 | 1 |
| COX4I1 | 0 | 1 | 0 | 0 | MAP2 | 1 | 0 | 0 | 1 | SYNPR | 0 | 0 | 1 | 0 |

| | | | | | | | | | | | | | | |
|---|---|---|---|---|---|---|---|---|---|---|---|---|---|---|
| CPNE5 | 0 | 0 | 0 | 1 | MAP4 | 2 | 0 | 1 | 0 | TALDO1 | 0 | 0 | 1 | 2 |
| CREB3 | 0 | 0 | 0 | 1 | MAPK4 | 0 | 0 | 1 | 0 | TAX1BP1 | 1 | 0 | 0 | 0 |
| CRTAP | 1 | 0 | 1 | 0 | MAZ | 0 | 0 | 1 | 0 | TBC1D17 | 0 | 0 | 0 | 1 |
| CTNNB1 | 0 | 1 | 0 | 0 | MBP | 0 | 0 | 0 | 1 | TCF25 | 0 | 0 | 1 | 0 |
| DAAM2 | 0 | 1 | 0 | 0 | MFN2 | 0 | 1 | 0 | 0 | TEX2 | 1 | 0 | 1 | 0 |
| DCTN1 | 1 | 0 | 0 | 0 | MPRIP | 0 | 0 | 1 | 1 | TF | 1 | 0 | 0 | 1 |
| DDX17 | 0 | 0 | 0 | 1 | MTMR10 | 0 | 0 | 1 | 0 | TJP1 | 0 | 0 | 0 | 1 |
| DDX24 | 1 | 0 | 0 | 0 | MTSS1L | 1 | 0 | 0 | 0 | TM9SF2 | 0 | 0 | 0 | 1 |
| DDX5 | 0 | 0 | 0 | 3 | MYH10 | 1 | 0 | 0 | 1 | TMEFF1 | 0 | 0 | 0 | 1 |
| DGCR2 | 0 | 0 | 1 | 0 | MYO5A | 0 | 0 | 1 | 1 | TMEM219 | 0 | 0 | 1 | 0 |
| DIRAS2 | 0 | 0 | 0 | 1 | NAT14 | 0 | 0 | 0 | 1 | TOP2B | 0 | 0 | 1 | 0 |
| DNAJA4 | 0 | 0 | 1 | 0 | NCAM1 | 0 | 0 | 0 | 1 | TPP1 | 0 | 0 | 1 | 0 |
| DPP6 | 0 | 0 | 1 | 0 | NCOA4 | 0 | 0 | 1 | 0 | TRAK2 | 0 | 0 | 0 | 1 |
| DTNA | 0 | 0 | 0 | 1 | NDEL1 | 0 | 1 | 0 | 0 | TRAP1 | 1 | 0 | 0 | 0 |
| DUSP1 | 0 | 0 | 0 | 2 | NDRG1 | 0 | 0 | 1 | 0 | TRIM44 | 0 | 0 | 0 | 1 |
| DYNC1H1 | 0 | 0 | 0 | 2 | NDRG4 | 0 | 1 | 0 | 0 | TSPAN7 | 0 | 0 | 0 | 1 |
| ECH1 | 0 | 0 | 1 | 0 | NDUFS2 | 0 | 0 | 0 | 1 | TSPYL4 | 0 | 0 | 0 | 1 |
| EEF2 | 0 | 1 | 0 | 0 | NDUFV1 | 0 | 0 | 1 | 0 | TTC1 | 1 | 0 | 0 | 0 |
| EIF2AK1 | 1 | 0 | 0 | 0 | NECAB1 | 0 | 0 | 1 | 0 | TTYH1 | 0 | 0 | 0 | 1 |
| EIF3A | 1 | 0 | 0 | 1 | NECAB2 | 0 | 0 | 2 | 0 | TUFM | 0 | 0 | 0 | 2 |
| EIF3L | 0 | 1 | 0 | 0 | NFE2L1 | 0 | 0 | 0 | 1 | TXNIP | 0 | 0 | 1 | 0 |
| EIF4G1 | 0 | 0 | 1 | 0 | NME1-NME2 | 0 | 0 | 1 | 1 | UBXN6 | 0 | 0 | 0 | 1 |
| EIF6 | 0 | 0 | 0 | 1 | NME2 | 0 | 0 | 1 | 0 | UGT8 | 0 | 0 | 0 | 1 |
| ELMO1 | 0 | 0 | 0 | 2 | NPC1 | 0 | 0 | 0 | 2 | UNC13A | 0 | 0 | 0 | 1 |
| ELMOD1 | 0 | 0 | 1 | 0 | NPDC1 | 0 | 0 | 0 | 1 | UQCRC2 | 0 | 0 | 0 | 1 |
| ELOVL5 | 0 | 1 | 0 | 0 | NR1D1 | 0 | 0 | 2 | 0 | UROD | 0 | 1 | 0 | 1 |
| ENTPD3 | 0 | 0 | 0 | 1 | NTSR2 | 0 | 0 | 1 | 0 | USP11 | 0 | 1 | 0 | 0 |
| EPAS1 | 0 | 1 | 0 | 0 | NUDCD3 | 0 | 0 | 1 | 0 | USP5 | 0 | 0 | 0 | 1 |
| EPB41L1 | 0 | 0 | 0 | 1 | OLFM1 | 0 | 0 | 1 | 0 | VAT1 | 0 | 1 | 1 | 0 |
| EPHB6 | 0 | 0 | 0 | 1 | OLFM2 | 0 | 1 | 0 | 0 | VCP | 0 | 0 | 0 | 1 |
| EPHX1 | 0 | 1 | 0 | 0 | PAM | 0 | 1 | 1 | 1 | WNK1 | 1 | 1 | 0 | 1 |
| EPN1 | 2 | 0 | 1 | 0 | PCDHGB2 | 0 | 0 | 1 | 0 | XPO6 | 0 | 0 | 0 | 1 |
| ESD | 0 | 0 | 1 | 0 | PCDHGC3 | 0 | 0 | 0 | 1 | ZFAND5 | 0 | 0 | 0 | 1 |
| ETFB | 0 | 0 | 0 | 1 | PDE2A | 0 | 1 | 0 | 1 | ZMAT2 | 0 | 0 | 0 | 1 |
| | | | | | | | | | | ZSCAN18 | 0 | 1 | 0 | 0 |

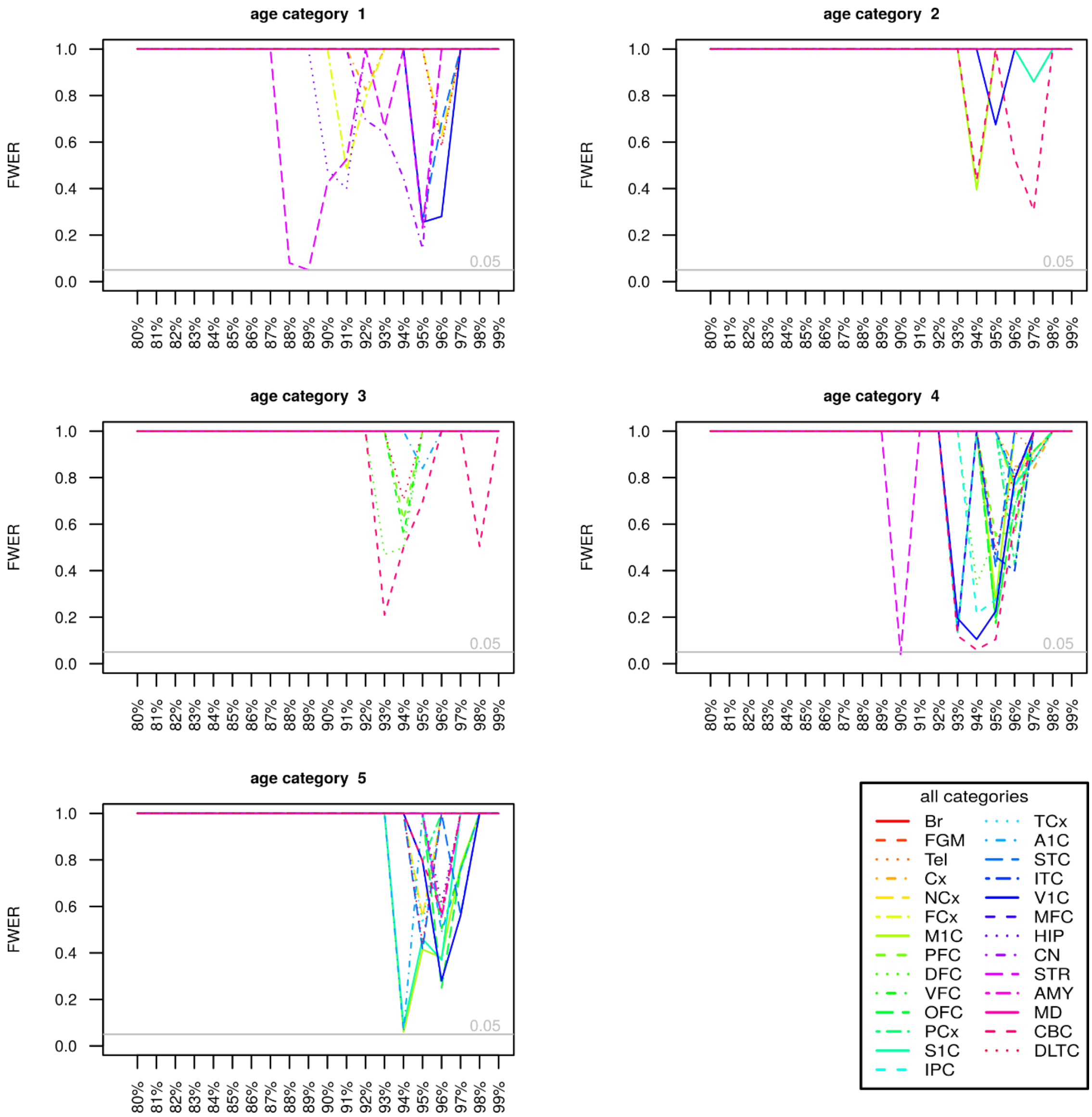


**Figure S9.2: Family-wise error rates (FWER) of the McDonald-Kreitmann test for Neandertal-specific variants, comparing different brain regions and developmental states (age categories, labelled as in Table S9.3) at different expression cutoffs (x-axis).** The number of genes per brain regions are reported in Fig.S9.3. The brain regions are indicated as: STR, striatum; CBC cerebellar cortex; M1C, primary motor cortex (area M1, area 4); A1C, primary auditory cortex (core); S1C, primary somatosensory cortex (area S1, areas 3,1,2); V1C, primary visual cortex (striate cortex, area V1/17); MFC, anterior (rostral) cingulate (medial prefrontal) cortex; CN, cerebral nuclei; VFC, ventrolateral prefrontal cortex; OFC orbital frontal cortex; IPC, posteroventral (inferior) parietal cortex; STC, posterior (caudal) superior temporal cortex (area 22c); DFC, dorsolateral prefrontal cortex; ITC, inferolateral temporal cortex (area TEv, area 20); HIP, hippocampus (hippocampal formation); PCx, parietal neocortex; DLTC, dorsolateral temporal neocortex; FCx, frontal neocortex; TCx, temporal neocortex; PFC, prefrontal cortex; AMY, amygdaloid complex; NCx, neocortex (isocortex); Tel, telencephalon; Cx, cerebral cortex; Br, brain; FGM, gray matter of forebrain; MD, mediodorsal nucleus of thalamus.

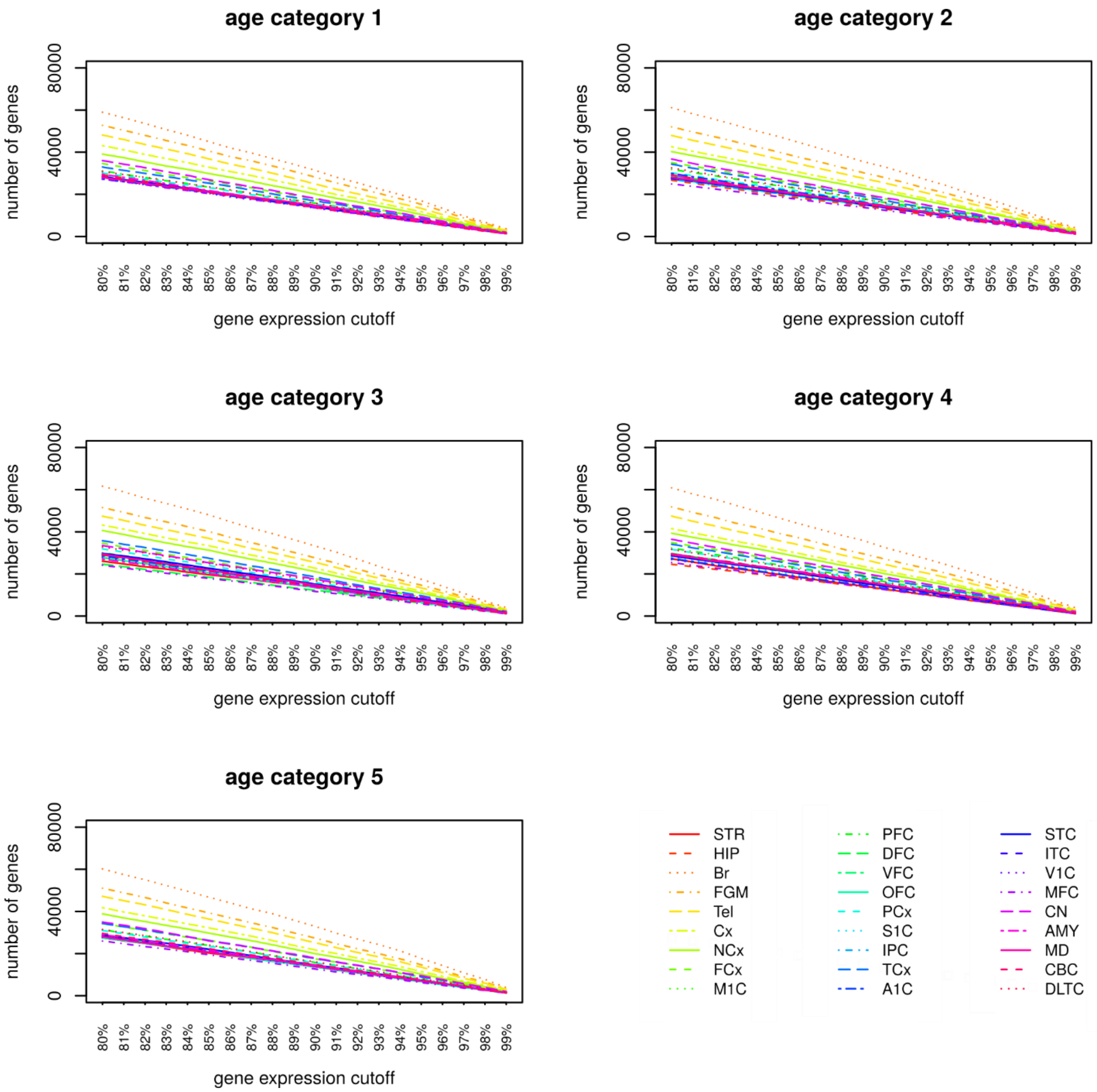


**Figure S9.3: Number of genes expressed in the different brain regions at the different gene expression cutoffs.** Brain regions are denoted as in Figure S9.2.

We note that it has previously been shown that "Neandertal deserts" (16), genomic regions among present-day humans that are depleted of Neandertal and Denisovan introgressed fragments, are enriched in genes expressed in the striatum in the adult (>19 years) and adolescent (12-19 years) brain (16,17). We find that three genes out of 34, present in deserts and expressed in the striatum at the 90% cutoff in the adolescent brain, have fixed non-synonymous changes (*MAP2, AMPD2* and *PTPRZ1)* (Figure S9.5). The proportion of genes carrying non-synonymous changes in non-deserts is ~5 times lower (23 genes out of 1403, Fisher's exact test p=0.0217). This could suggest that in the striatum, variants that evolved either neutrally or under positive selection in Neandertals, would have been selected against once introgressed in modern humans.

Finally, we investigated whether genes associated with specific phenotypes have elevated McDonald-Kreitman ratios compared to other genes. We find no significant enrichment for any category (not shown).

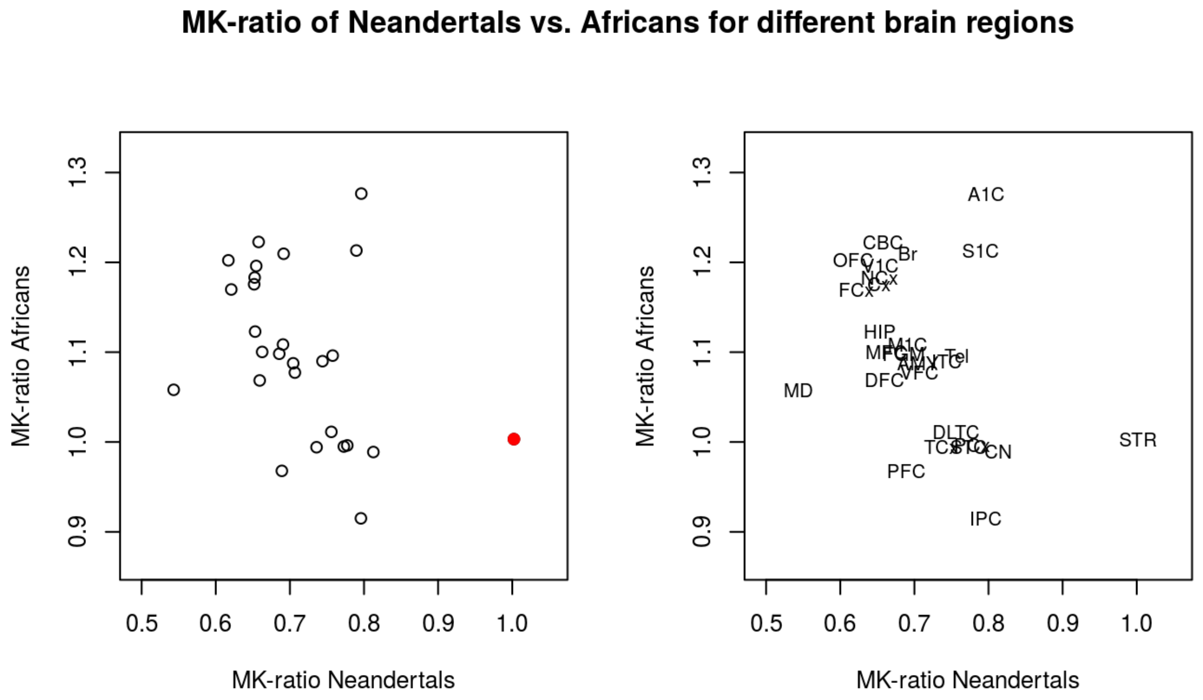


**Figure S9.4: Relationship between McDonald-Kreitman ratios in Africans of the SGDP dataset (y-axis) and Neandertals (x-axis) per brain region.** Striatum is highlighted in red in the left panel, while labels are reported in the right one following the Allen Brain Atlas nomenclature reported in Figure S9.2.

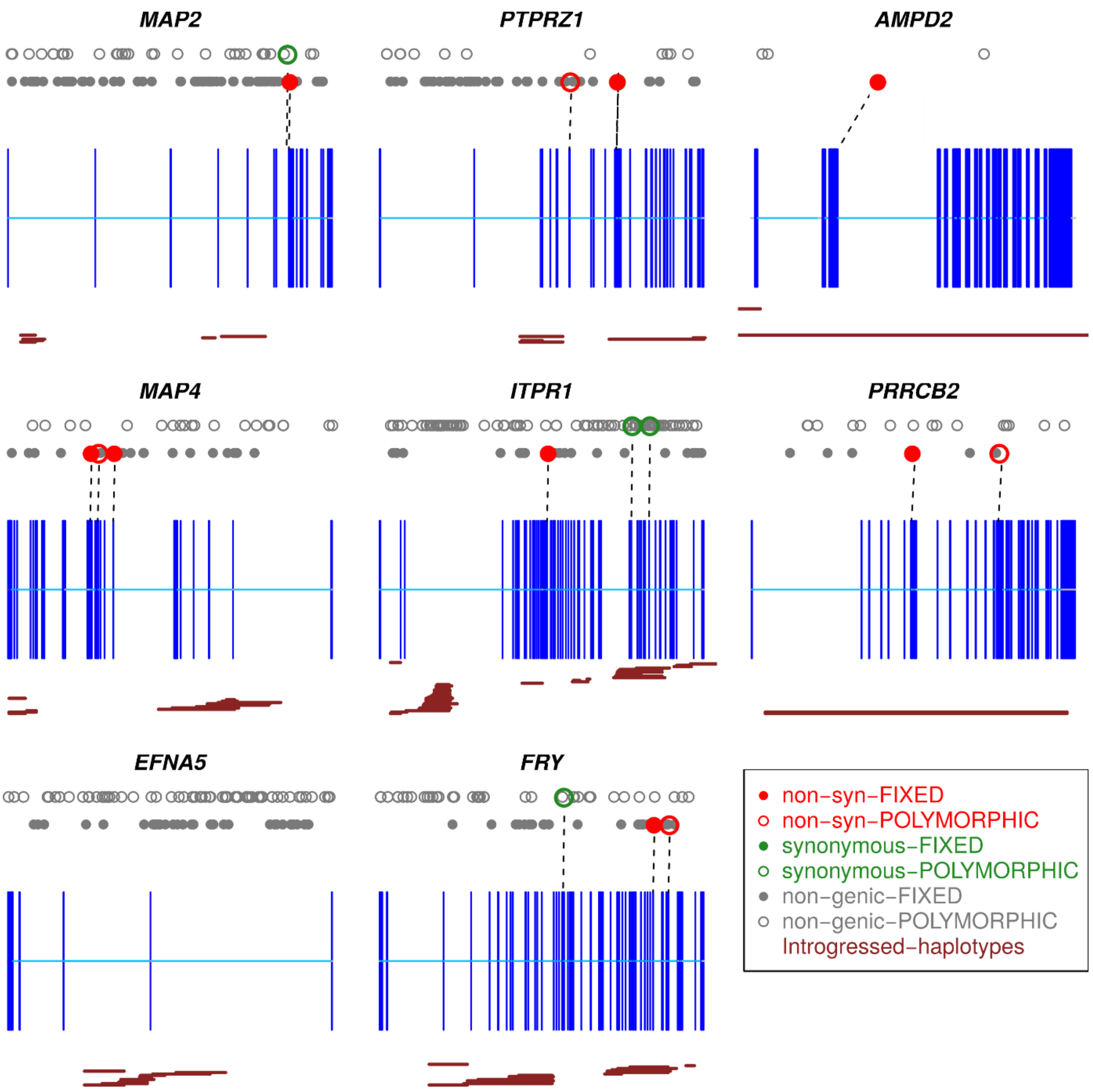


**Figure S9.5: The three genes expressed in the striatum and present in the deserts (top row) and three random genes expressed in the striatum but not present in deserts (middle row) are shown, compared to two genes sampled at random from the list of genes not expressed in the striatum (bottom row).** Exons are represented in blue, while Neandertal haplotypes detected in modern human populations (18) in dark red. Changes present in all Neandertals are shown as full dots in the bottom row. Changes polymorphic in Neandertals are shown as empty circles in the top row. Red and green indicate non-synonymous and synonymous changes, respectively. Random genes were sampled without replacement from the list of genes with at least 30% of the exon sequences passing all general filters (Supplementary Information 3).

**Putatively regulatory regions**

We considered promoters and untranslated regions (UTRs) annotated by ENSEMBL as putatively regulatory regions and tested them separately. We associate each regulatory region to the closest gene to define the regulatory target. We excluded sites that showed extreme values of coverage, low genotype quality and mappability as described in Supplementary Information 3. Within these regions, we counted the number of derived variants shared by three Neandertal genomes as described above. We normalized the number of fixed derived Neandertal variants by the number of polymorphic variants in Neandertals. For each of the three ontologies described above, we then performed a binomial test as implemented in the R-package GOfuncR, comparing the density of regulatory changes per each gene category.

Neandertal-specific variants were not associated with any particular GO or HPO category in neither the UTRs or promoters (Table S9.5 and Table S9.6). We also did not detect any significant enrichment in the adult brain, for any of the tests (Table S9.7). In the developing brain, we detect a significant enrichment for regions regulating genes expressed in the posteroventral (inferior) parietal cortex, in both UTRs (FWER=0.022) and promoters (FWER=0.019) (Table S9.8). Two other regions appear among the top six regions with the lowest FWER for UTRs and promoters: the posterior (caudal) superior temporal cortex (area 22c) and the cerebellar cortex. In addition, the striatum is significant for age category 1 (prenatal) for UTRs (FWER=0.049).

**Table S9.5: Top six categories from a GO-enrichment analysis of Neandertal-specific regulatory changes, normalizing the number of fixed Neandertal changes by the number of polymorphic changes.** A low FWER indicates categories where this ratio is comparatively high.

| UTRs | | | | |
|---|---|---|---|---|
| **ontology** | **node id** | **node name** | **p-value** | **FWER** |
| cellular_component | GO:0044432 | endoplasmic reticulum part | 0.2e-3 | 0.336 |
| molecular_function | GO:0061650 | ubiquitin-like protein conjugating enzyme activity | 0.2e-3 | 0.378 |
| cellular_component | GO:0030992 | intraciliary transport particle B | 0.3e-3 | 0.44 |
| cellular_component | GO:0005789 | endoplasmic reticulum membrane | 0.4e-3 | 0.479 |
| cellular_component | GO:0042175 | nuclear outer membrane-endoplasmic reticulum membrane network | 0.4e-3 | 0.496 |
| cellular_component | GO:0098827 | endoplasmic reticulum subcompartment | 0.4e-3 | 0.507 |

| Promoters | | | | |
|---|---|---|---|---|
| molecular_function | GO:0005089 | Rho guanyl-nucleotide exchange factor activity | 4.0e-3 | 0.57 |
| cellular_component | GO:0044853 | plasma membrane raft | 8.0e-3 | 0.634 |
| molecular_function | GO:0042578 | phosphoric ester hydrolase activity | 8.6e-3 | 0.721 |
| molecular_function | GO:0017048 | Rho GTPase binding | 10.6e-3 | 0.795 |
| biological_process | GO:1901566 | organonitrogen compound biosynthetic process | 3.46e-3 | 0.83 |
| cellular_component | GO:0005901 | caveola | 20.9e-3 | 0.948 |
| biological_process | GO:0034109 | homotypic cell-cell adhesion | 3.9e-3 | 0.953 |

**Table S9.6: Top six categories from an HPO-enrichment analysis of Neandertal-specific regulatory changes, normalizing the number of fixed Neandertal changes by the number of polymorphic changes.**

| **UTRs** | | | |
|---|---|---|---|
| **node id** | **node name** | **p-value** | **FWER** |
| HP:0002476 | Primitive reflex | 0.3e-3 | 0.729 |
| HP:0002127 | Abnormal upper motor neuron morphology | 2.1e-3 | 0.994 |
| HP:0002193 | Pseudobulbar behavioral symptoms | 2.8e-3 | 0.99 |
| HP:0011347 | Abnormality of ocular abduction | 2.8e-3 | 0.99 |
| HP:0000482 | Microcornea | 5.2e-3 | 0.99 |
| HP:0010994 | Abnormality of the striatum | 5.3e-3 | 0.99 |
| **Promoters** | | | |
| HP:0011968 | Feeding difficulties | 5.5e-3 | 0.656 |
| HP:0000426 | Prominent nasal bridge | 6.7e-3 | 0.85 |
| HP:0010013 | Abnormality of the 5th metacarpal | 11.2e-3 | 0.938 |
| HP:0006712 | Aplasia/Hypoplasia of the ribs | 19.7e-3 | 0.989 |
| HP:0008368 | Tarsal synostosis | 20.1e-3 | 0.998 |
| HP:0009834 | Abnormality of the proximal phalanges of the hand | 20.1e-3 | 0.998 |
| HP:0000253 | Progressive microcephaly | 20.1e-3 | 0.998 |

**Table S9.7 Top six categories for the ABAEnrichment analysis of the Neandertal specific changes regulating genes expressed in the adult brain, normalizing the number of fixed Neandertal changes by the number of polymorphic changes.**

| UTRs | | |
|---|---|---|
| **Structure id** | **Region** | **Minimum FWER** |
| Allen:4743 | He-VIIIB_VIIIB, Left Lateral Hemisphere | 0.094 |
| Allen:4703 | Ve-V_V | 0.118 |
| Allen:4734 | He-III_III, Left Lateral Hemisphere | 0.13 |
| Allen:4726 | PV-VIIIA_VIIIA, Left Paravermis | 0.132 |
| Allen:4707 | Ve-VIIAt_VIIAt | 0.14 |
| Allen:4724 | PV-Crus II_Crus II, Left Paravermis | 0.147 |
| **Promoter** | | |
| Allen:4751 | PV-III_III, Right Paravermis | 0.258 |
| Allen:4752 | PV-IV_IV, Right Paravermis | 0.27 |
| Allen:4701 | Ve-III_III | 0.301 |
| Allen:4775 | He-VIIIA_VIIIA, Right Lateral Hemisphere | 0.308 |
| Allen:4700 | Ve-I-II_I-II | 0.344 |
| Allen:4706 | Ve-VIIAf_VIIAf | 0.367 |

**Table S9.8: Top six categories for the ABAEnrichment analysis of the Neandertal-specific changes regulating genes expressed in the developing brain, normalizing the number of fixed Neandertal changes by the number of polymorphic changes.**

| UTRs | | | |
|---|---|---|---|
| **age_category** | **Structure id** | **structure** | **FWER** |
| 1 (prenatal) | Allen:10225 | IPC_posteroventral (inferior) parietal cortex | 0.022 |
| 1 (prenatal) | Allen:10208 | PCx_parietal neocortex | 0.026 |
| 1 (prenatal) | Allen:10209 | S1C_primary somatosensory cortex (area S1, areas 3,1,2) | 0.027 |
| 1 (prenatal) | Allen:10333 | STR_striatum | 0.049 |
| 1 (prenatal) | Allen:10243 | STC_posterior (caudal) superior temporal cortex (area 22c) | 0.051 |
| 1 (prenatal) | Allen:10657 | CBC_cerebellar cortex | 0.066 |
| **Promoters** | | | |
| 1 (prenatal) | Allen:10225 | IPC_posteroventral (inferior) parietal cortex | 0.019 |
| 1 (prenatal) | Allen:10185 | VFC_ventrolateral prefrontal cortex | 0.036 |
| 1 (prenatal) | Allen:10236 | A1C_primary auditory cortex (core) | 0.078 |
| 1 (prenatal) | Allen:10208 | PCx_parietal neocortex | 0.08 |
| 1 (prenatal) | Allen:10243 | STC_posterior (caudal) superior temporal cortex (area 22c) | 0.103 |
| 1 (prenatal) | Allen:10252 | ITC_inferolateral temporal cortex (area TEv, area 20) | 0.104 |
| 5 (>19 years old) | Allen:10657 | CBC_cerebellar cortex | 0.113 |

### Supplementary Information 10

#### Selection scan

**Summary:** We applied two different statistics, the Population Branch Statistics (PBS) (1) and HKA (2), to identify targets of natural selection in the three high-coverage Neandertal genomes, *Vindija 33.19* (3), *Denisova 5* (4) and *Chagyrskaya 8*.

##### Methods

For the HKA test, we computed the ratio between fixed and polymorphic derived variants. We restricted our analyses to sites in which either the bonobo or chimpanzee genomes (*panTro4*) and at least one more ape genome was available, among gorilla (*gorGor3*) and orangutan (*ponAbe2*), and at which all ape genomes or all but one shared the same state. To restrict the HKA test to derived variants which evolved in the Neandertal lineage we also excluded variants that showed allele frequencies higher than 5% in Yorubas from the 1000 Genomes dataset (5).

The PBS statistic aims at identifying regions for which allele frequencies changed more in one population compared to two other populations, measuring differences in allele frequencies among populations in terms of Fst. Besides Neandertals we considered Yoruba and Denisovans, the latter represented by the single genome of *Denisova 3* (6). To account for the small sample sizes and the divergence time of more than 400 ky, we modified the PBS statistic using an Fst estimator developed by Reich and colleagues (7), which accounts for inbreeding and is more accurate for small sample sizes (8). In addition, we added to all population a dummy heterozygote genome to be able to apply this statistic also to sites that are fixed in one population for an allele absent in the other two, a category of sites usually discarded in PBS analyses.

We computed both statistics in sliding windows of 25kb (with steps of 6250bp, respectively) across the genomes, excluding windows with less than 4 informative sites for HKA and PBS. We computed neutral expectations by performing 200 whole-genome coalescent simulations following the demographic histories inferred in Supplementary Information 7 and Prüfer and colleagues (3), and applying the same masks used for the real data to obtain comparable datasets. For the X chromosome we generated independent simulations, setting the population size to ¾ that of the autosomes. To control for the different amount of informative sites among windows, we stratify windows obtained from real data and simulated genomes in bins containing different number of informative sites. For each observed window we then computed a p-value, calculating the proportion of simulated windows in the same bin, hence with a comparable proportion of informative sites, showing a value of the focal statistic equal or more extreme. For all analyses, we also compute an empirical False Discovery Rate (FDR), by calculating for each p-value *p* the ratio between the number of observed windows with p-values equal or lower than simulated ones (9). Note that the exact p-values depend on the specific simulated demography. Thus, we also adopt an outlier approach by reporting the top 500 candidate windows, that constitute the top 0.1 of the total 25kb autosomal windows, respectively.

We contrasted candidate regions with GWAS associated variants and coding regions downloaded from the UCSC website (10), the GTEX (11) website, and genomic regions reported in Peyrégne and colleagues (12).

**Results**

On the autosomes we identified 36 and 38 windows with FDR<5% using HKA and PBS, respectively (Figure 4, Supplementary Table 1-4), while only one region was detected using HKA on the X-chromosome (Supplementary Figure 10.1). When we merge contiguous windows, the numbers of disjoint autosomal regions are 20 and 17 for the two statistics, respectively. Among them, a single region identified by PBS partially overlaps two smaller candidates regions identified by HKA, which are less than 10kb apart on chromosome 5. To estimate the probability that such overlap would occur by chance we generated random candidate regions with the same length as the observed ones in terms of contiguous windows, and calculated the proportion of cases in which higher or lower overlaps would be observed. We estimate that such overlaps would occur with probabilities smaller than $4x10^{-4}$.

Interestingly, 5 HKA and 2 PBS candidate regions overlap previously reported candidates of positive selection in modern humans identified by extended lineage sorting (ELS) (12), i.e. long genomic regions (>0.1centiMorgan) for which Neandertals fall outside the variation of modern humans. The overlap between ELS and HKA/PBS is higher than would be expected by chance (p-value 0.010, 0.056 for HKA and PBS). Note that this signal is even stronger when the top 500 candidates for HKA and PBS are considered (p-value $<10^{-4}$ for both tests). The overlap between ELS and HKA/PBS could be potentially explained in two different ways: first, diversifying selection or other mechanisms could lead to the rapid evolution of certain genomic regions independently in Neandertals and modern humans (13); second, the genomic signatures identified by ELS and the methods here used are not entirely independent, and thus selection on one lineage might be mistaken as selection on another lineage by one of the statistics. For example, extended lineage sorting could be generated by the replacement of Neandertal haplotypes falling within the variation of modern humans by a second Neandertal haplotype falling out of the variation of modern humans, because of drift or selection. We thus performed the PBS scan performed on Neandertals also on modern humans, and compared the two sets of candidates. As expected, regions identified in modern humans overlap the regions identified by Peyrégne et al. (14.3%) more extensively than regions identified in Neandertals (7.5%). However, the two sets of candidates show no overlap, indicating that a subset of regions previously detected with ELS might have undergone selection in Neandertals.

We then tested the overlap of the candidate windows identified in the Neandertals with genomic functional annotations. Interestingly, coding regions and genes are not overrepresented among candidate windows. Rather, coding regions (p-values=0.355,0.046 for HKA and PBS, respectively) and GTEX genes (11) (p-values=0.104,0.009 for HKA and PBS) appear as generally depleted among candidate regions.  We report the genes whose coding regions are spanned by candidate windows in Supplementary Table 10.3. Among these genes we find HTN1, which determine tooth enamel and resistance to caries (14), or EXOC6B, which has effects on neural development (15).

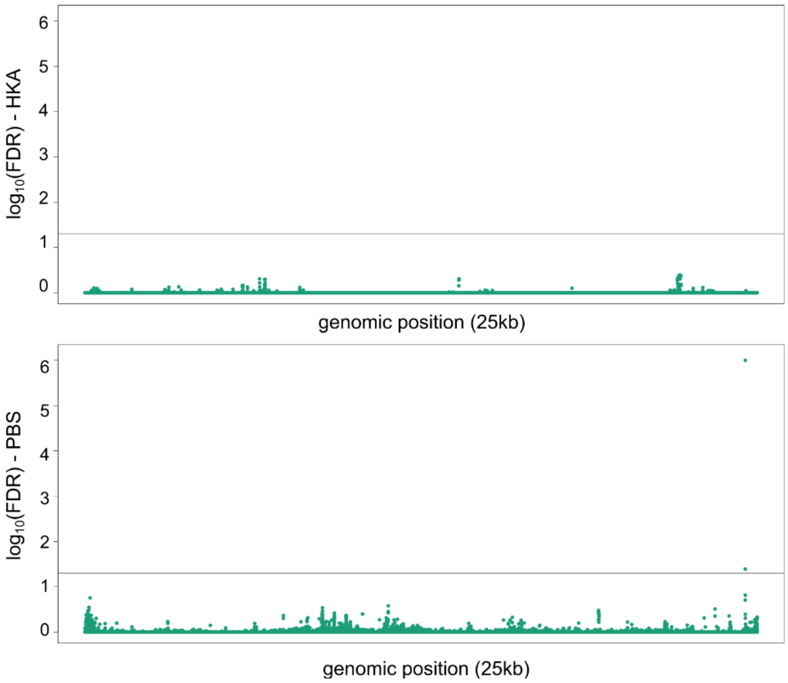


**Figure S10.1:** Manhattan plots of False Discovery Rates for different neutrality tests applied on the Neandertal chromosome X: HKA (top panel), PBS (bottom panel). The y-axis indicates the -log10 of the False Discovery Rate (FDR) up to 6, i.e. FDR=$10^{-6}$. The black continuous line indicates FDR=5%.

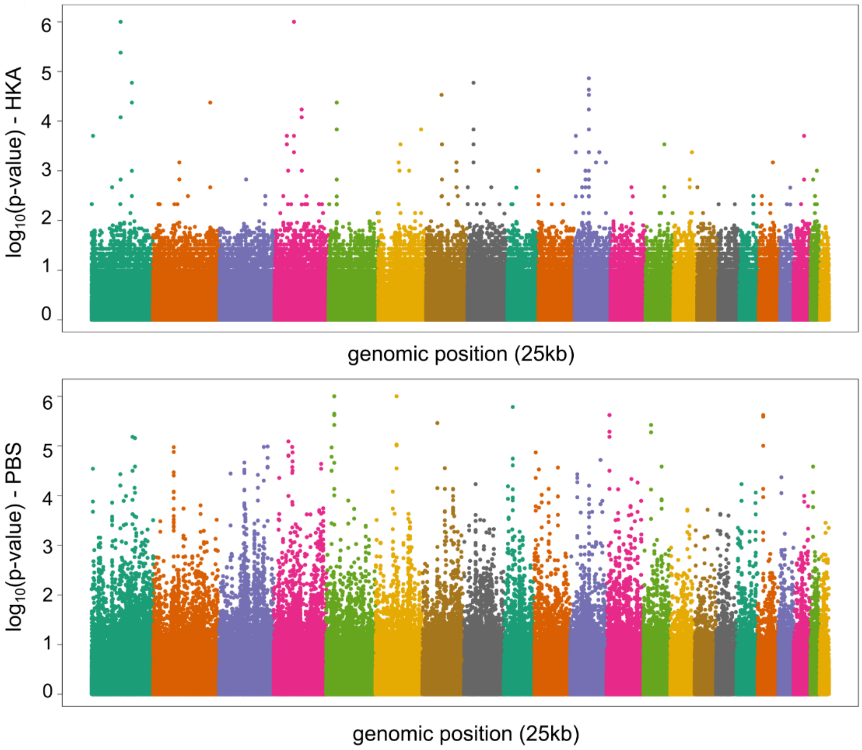


**Figure S10.2:** Manhattan plots showing the distribution of p-values for different neutrality tests applied on the Neandertal autosomes: HKA (top) and PBS (bottom). The y-axis indicates the -log10 of the p-values up to 6, i.e. FDR=$10^{-6}$.

**Table S10.1:** Table of windows with FDR<0.05 for the HKA neutrality test. The number of polymorphisms (P) and fixed differences (D) are used to compute the HKA ratio.

| Chromosome | INIT | END | P | D | FDR | p-value |
|---|---|---|---|---|---|---|
| 1 | 5410502 | 5435502 | 2 | 32 | 0.034 | 0.000 |
| 1 | 105010502 | 105035502 | 3 | 56 | 0.004 | 0.000 |
| 1 | 105016752 | 105041752 | 2 | 57 | 0.000 | 0.000 |
| 1 | 105041752 | 105066752 | 2 | 34 | 0.022 | 0.000 |
| 1 | 175229252 | 175254252 | 2 | 39 | 0.010 | 0.000 |
| 1 | 175235502 | 175260502 | 2 | 36 | 0.014 | 0.000 |
| 2 | 211680199 | 211705199 | 2 | 36 | 0.014 | 0.000 |
| 4 | 42850710 | 42875710 | 2 | 31 | 0.041 | 0.000 |
| 4 | 42856960 | 42881960 | 2 | 32 | 0.034 | 0.000 |
| 4 | 70919460 | 70944460 | 3 | 43 | 0.034 | 0.000 |
| 4 | 70925710 | 70950710 | 2 | 55 | 0.000 | 0.000 |
| 4 | 70931960 | 70956960 | 2 | 59 | 0.000 | 0.000 |
| 4 | 70938210 | 70963210 | 3 | 40 | 0.047 | 0.000 |
| 4 | 97506960 | 97531960 | 2 | 34 | 0.022 | 0.000 |
| 4 | 97513210 | 97538210 | 2 | 35 | 0.017 | 0.000 |
| 5 | 28828757 | 28853757 | 2 | 33 | 0.031 | 0.000 |
| 5 | 28860007 | 28885007 | 2 | 36 | 0.014 | 0.000 |
| 6 | 82677585 | 82702585 | 2 | 31 | 0.041 | 0.000 |
| 6 | 153865085 | 153890085 | 2 | 33 | 0.031 | 0.000 |
| 7 | 54122170 | 54147170 | 2 | 37 | 0.013 | 0.000 |
| 7 | 55540920 | 55565920 | 2 | 31 | 0.041 | 0.000 |
| 7 | 55547170 | 55572170 | 2 | 31 | 0.041 | 0.000 |
| 7 | 55578420 | 55603420 | 2 | 31 | 0.041 | 0.000 |
| 8 | 21033254 | 21058254 | 3 | 41 | 0.039 | 0.000 |
| 8 | 21039504 | 21064504 | 2 | 39 | 0.010 | 0.000 |
| 8 | 21045754 | 21070754 | 2 | 33 | 0.031 | 0.000 |
| 11 | 4791086 | 4816086 | 2 | 32 | 0.034 | 0.000 |
| 11 | 55678586 | 55703586 | 3 | 40 | 0.047 | 0.000 |
| 11 | 55691086 | 55716086 | 2 | 38 | 0.012 | 0.000 |
| 11 | 55697336 | 55722336 | 2 | 35 | 0.017 | 0.000 |
| 11 | 55716086 | 55741086 | 2 | 40 | 0.011 | 0.000 |
| 11 | 55722336 | 55747336 | 3 | 40 | 0.047 | 0.000 |
| 11 | 55816086 | 55841086 | 2 | 37 | 0.013 | 0.000 |
| 11 | 55834836 | 55859836 | 2 | 33 | 0.031 | 0.000 |
| 13 | 84352797 | 84377797 | 2 | 31 | 0.041 | 0.000 |
| 20 | 41098070 | 41123070 | 2 | 32 | 0.034 | 0.000 |

**Table S10.2:** Table of windows with FDR<0.05 for the PBS neutrality test.

| Chromosome | INIT | END | PBS | FDR | p-value |
|---|---|---|---|---|---|
| 1 | 103898002 | 103923002 | 0.642 | 0.044 | 0.000 |
| 1 | 103904252 | 103929252 | 0.642 | 0.044 | 0.000 |
| 1 | 103910502 | 103935502 | 0.642 | 0.044 | 0.000 |
| 2 | 73036449 | 73061449 | 0.589 | 0.039 | 0.000 |
| 3 | 93823294 | 93848294 | 0.601 | 0.038 | 0.000 |
| 3 | 93829544 | 93854544 | 0.620 | 0.035 | 0.000 |
| 3 | 162123294 | 162148294 | 0.639 | 0.043 | 0.000 |
| 3 | 175698294 | 175723294 | 0.614 | 0.032 | 0.000 |
| 3 | 175710794 | 175735794 | 0.651 | 0.039 | 0.000 |
| 3 | 175717044 | 175742044 | 0.624 | 0.022 | 0.000 |
| 4 | 56050710 | 56075710 | 0.631 | 0.030 | 0.000 |
| 4 | 56056960 | 56081960 | 0.662 | 0.022 | 0.000 |
| 4 | 69438210 | 69463210 | 1.123 | 0.009 | 0.000 |
| 4 | 69444460 | 69469460 | 1.123 | 0.009 | 0.000 |
| 4 | 69450710 | 69475710 | 1.123 | 0.009 | 0.000 |
| 4 | 69456960 | 69481960 | 1.114 | 0.007 | 0.000 |
| 5 | 28816257 | 28841257 | 0.776 | 0.000 | 0.000 |
| 5 | 28822507 | 28847507 | 0.748 | 0.040 | 0.000 |
| 5 | 28828757 | 28853757 | 0.721 | 0.032 | 0.000 |
| 5 | 28847507 | 28872507 | 0.725 | 0.020 | 0.000 |
| 5 | 28853757 | 28878757 | 0.767 | 0.000 | 0.000 |
| 5 | 28860007 | 28885007 | 0.734 | 0.027 | 0.000 |
| 5 | 28866257 | 28891257 | 0.745 | 0.021 | 0.000 |
| 6 | 79352585 | 79377585 | 0.710 | 0.000 | 0.000 |
| 7 | 54274249 | 54299249 | 0.752 | 0.004 | 0.000 |
| 8 | 47183254 | 47208254 | 0.630 | 0.035 | 0.000 |
| 9 | 31760450 | 31785450 | 0.740 | 0.002 | 0.000 |
| 9 | 31766700 | 31791700 | 0.626 | 0.031 | 0.000 |
| 12 | 10914574 | 10939574 | 0.716 | 0.044 | 0.000 |
| 12 | 10920824 | 10945824 | 0.722 | 0.018 | 0.000 |
| 12 | 10927074 | 10952074 | 0.734 | 0.023 | 0.000 |
| 12 | 10933324 | 10958324 | 0.703 | 0.049 | 0.000 |
| 13 | 46965297 | 46990297 | 0.687 | 0.007 | 0.000 |
| 13 | 46971547 | 46996547 | 0.741 | 0.007 | 0.000 |
| 17 | 18269104 | 18294104 | 0.630 | 0.026 | 0.000 |
| 18 | 24454529 | 24479529 | 0.746 | 0.032 | 0.000 |
| 18 | 24460779 | 24485779 | 0.720 | 0.006 | 0.000 |
| 19 | 11034911 | 11059911 | 0.645 | 0.038 | 0.000 |

**Table S10.3:** List of contiguous regions overlapping the coding exons of the reported genes.

| HKA | | | |
|---|---|---|---|
| *Chromosome* | *Init* | *End* | *Gene* |
| 4 | 70919460 | 70963210 | *HTN1* |
| 7 | 55540920 | 55572170 | *VOPP1* |
| 11 | 4791086 | 4816086 | *OR51F1* |
| 11 | 55678586 | 55747336 | *OR5W2* |
| **PBS** | | | |
| *Chromosome* | *Init* | *End* | *Gene* |
| 2 | 73036449 | 73061449 | EXOC6B |
| 3 | 93823294 | 93854544 | NSUN3 |
| 12 | 10914574 | 10958324 | TAS2R7 |
| 17 | 18269104 | 18294104 | EVPLL |
| 19 | 11034911 | 11059911 | YIPF2, TIMM29 |
